# A Combined Behavioral and Neuroimaging Battery to Test Positive Appraisal Style Theory of Resilience in Longitudinal Studies

**DOI:** 10.1101/470435

**Authors:** Miriam Kampa, Anita Schick, Kenneth Yuen, Alexandra Sebastian, Andrea Chmitorz, Victor Saase, Michèle Wessa, Oliver Tüscher, Raffael Kalisch

**Author notes:** **Correspondence:** Miriam Kampa, Raffael Kalisch, Johannes Gutenberg University Medical Center, Langenbeckstr. 1, 55131 Mainz, Germany.

## Abstract

Resilience is the maintenance or rapid recovery of mental health during and after stressor exposure. It is becoming increasingly clear that resilience results from a complex and dynamic process of adaptation to stressors involving the biological, psychological and social levels. Positive appraisal style theory of resilience (PASTOR) claims that the common final pathway to maintained mental health lies in the non-negative (non-catastrophizing, non-pessimistic) appraisal of potential stressors, permitting the organism to fine-tune stress responses to optimal levels, thus avoiding unnecessary stress, inefficient deployment of resources and concomitant deleterious allostatic load effects. Successful adaptation consists in developing or strengthening a positive appraisal style. Another key element of PASTOR is that a positive appraisal style is strongly determined by the efficacy and efficiency of the neural and cognitive processes that underlie positive stressor appraisal. We here present a combined behavioral and functional magnetic resonance imaging (fMRI) battery designed to assess such processes repeatedly in longitudinal settings. The battery includes tests of stress reactivity and recovery, reward sensitivity, safety learning and memory in the context of fear conditioning and extinction, volitional situation-focused reappraisal, volitional self-focused reappraisal, and emotional interference inhibition, along with structural MRI and resting-state MRI scans. A detailed description of the battery methods is provided. The feasibility of the battery was successfully tested in N=55 healthy subjects; group results of the individual tasks largely replicate existing literature.

## Introduction

The prevention of mental disease and the promotion of mental health are currently major challenges for Western societies. So, in Europe alone, it is estimated that about a third of the population each year suffer from a mental disease such as anxiety, depression, post-traumatic stress disorder (PTSD) or also chronic pain or addiction, whose etiology or maintenance is at least in part related to the occurrence of stress (Wittchen et al., 2011). On the positive side, many individuals exposed to adversity are resilient to the development of lasting stress-related mental dysfunctions. Even though up to 90% of people in Europe and the US experience at least one potentially traumatic event in their lives, the life time prevalence of PTSD is low, with a rate of approximately 8% (de Vries & Olff, 2009; Kilpatrick et al., 2013). Also, the life time prevalence of any mental disorder is considerably lower than the occurrence of significant life events (approximately 20 to 50%; Kessler et al., 2007). Furthermore, prospective studies have shown that of those subjects who were confronted with adversity around two thirds did either not show any psychological or functional impairment or recovered quickly (Bonanno, Kennedy, Galatzer-Levy, Lude, & Elfström, 2012; Bonanno, Mancini, et al., 2012; Mancini, Bonanno, & Sinan, 2015; Werner, 1992). These data make the study of resilient individuals and the mechanisms that protect them from lasting impairments a promising strategy to find new ways of reducing stress-related suffering also in vulnerable individuals and of alleviating the burden on health care systems.

### Background for battery development: resilience theory

An emerging consensus is that resilience should be defined as an outcome, that is, as the maintenance and/or quick recovery of mental health during and after times of adversity, such as trauma, difficult life circumstances, challenging life transitions, or physical illness (Kalisch et al., 2017). It is also increasingly accepted that resilience - as opposed to lasting stress-induced mental and functional impairments - is the result of a dynamic process of successful adaptation to stressors (Bonanno, Romero, & Klein, 2015; Kalisch et al., 2017; Kalisch, Müller, & Tüscher, 2015; Martha Kent, Davis, & Reich, 2014; Rutter, 2012; Sapienza & Masten, 2011).. Indeed, there is now ample evidence that individuals *change* while they successfully cope with stressors, whether this manifests at the level of altered perspectives on life (Joseph & Linley, 2006; Tedeschi, 2011), as emergence of new strengths or competences (Luthar, Cicchetti, & Becker, 2000), as partial immunization against the effects of future stressors (Seery, Holman, & Silver, 2010; Seery, Leo, Lupien, Kondrak, & Almonte, 2013), or also as epigenetic alterations and modified gene expression patterns (Boks et al., 2015; Breen et al., 2015). Furthermore, neurobiological studies suggest that such organismic adjustments are causal for the preservation of mental health (e.g. Friedman et al., 2014; Krishnan et al., 2007; Maier, 2015; Wang, Perova, Arenkiel, & Li, 2014). Hence, resilience is not simply inertia, or insensitivity to stressors, or merely a passive response to adversity (Russo, Murrough, Han, Charney, & Nestler, 2012). In the same vein, resilience can no longer be understood simply as a stable, fixed personality trait or predisposition (the “resilient personality”) that guarantees long-term mental health whatever stressor the organism is exposed to. Rather, resilience research must attempt to understand the complex, interactive and time-varying processes that lead to a positive long-term outcome in the face of adversity (Kalisch et al., 2017).

While above results indicate that beneficial adaptation processes occur at biological, psychological and social levels, it has also been argued that these various processes converge in an individual learning or being able to optimally regulate his/her stress responses (Kalisch et al., 2015). This tenet is derived from a functional analysis of stress according to which stress responses are primarily adaptive reactions to potential threats to an organism’s needs and goals that serve to protect the organism from harm or damage and to preserve physiological homeostasis (Sterling & Eyer, 1988; Weiner, 1992). Albeit in principle protective, stress responses are also costly, by consuming energy, time, and cognitive capacity; by interfering with the pursuit of other important goals; and by often placing a burden on individual social, monetary and health resources. This implies that, if very intense, repeated or chronic, stress responses can become harmful themselves, as exemplified in the concept of “allostatic load” (McEwen, 1993). For these reasons, the organism needs regulatory mechanisms or “brakes” that fine-tune stress responses to optimal levels and, thus, preserve their primary adaptive function while at the same time assuring maximally efficient deployment of resources. Stress-regulatory mechanisms prevent a response over-shoot in amplitude or duration; shut off stress responses once the source of threat has vanished (“response termination” or “recovery”); and counteract response generalization. Rather than acting on the acute stress response, stress-regulatory mechanisms may also have an influence on how individuals respond to future exposures to the same or other stressors, by affecting, for instance, post-exposure evaluation or memory formation processes. Such flexible and adjustable responding to stressors (Ragland & Shulkin, 2014) limits resource consumption and maintains general functioning, thereby also allowing for the concurrent pursuit of other goals. Ultimately, it prevents the accumulation of allostatic costs and reduces the likelihood of developing lasting dysfunctions under stressor exposure (Kalisch et al., 2015). Hence, any biological, psychological and social adaptation processes most likely promote resilience in so far as they promote optimal stress response regulation. While some individuals may enter adverse life situations with already very efficient regulation capacities, most individuals presumably still considerably improve or even only develop such capacities in the confrontation with stressors (Kalisch et al., 2015).

This functional analysis permits to focus the investigation of protective adaptation processes to adaptations in the cognitive and neural mechanisms that underpin stress response regulation. A useful theoretical framework to approach these mechanisms is appraisal theory, which holds that the type, extent, and temporal evolution of emotional reactions, including acute and chronic stress responses, are not determined by simple, fixed stimulus-response relationships but by subjective and context-dependent appraisal (evaluation, analysis, interpretation) of the relevance of a stimulus or situation for the needs and goals of the organism (Arnold, 1969; Lazarus & Folkman, 1984; Scherer, 2001). Stress or threat responses, in particular, result from the appraisal of a situation as potentially harmful and as exceeding coping resources (Lazarus & Folkman, 1984). Both unconscious and conscious processes can contribute to this meaning analysis. Unconscious, non-verbal appraisal is presumably at the heart of phylogenetically old threat processing that also exists in animals. Conscious and explicit appraisal may be more dominant in unfamiliar and ambiguous situations and be restricted to humans (Leventhal & Scherer, 1987; Robinson, 1998). Appraisal processes have a neurobiological foundation (Kalisch & Gerlicher, 2014; Sander, Grandjean, & Scherer, 2005).

Positive appraisal style theory of resilience (PASTOR) proposes that individuals with a generalized tendency to appraise potentially threatening stimuli or situations in a nonnegative (non-pessimistic, non-catastrophizing) fashion are less likely to produce exaggerated, repeated and persistent stress responses and may thus be better protected against many long-term deleterious effects of trauma or chronic stressors (Kalisch et al., 2015). A positive appraisal style on average reduces the values that an individual attributes to stressors on the key threat appraisal dimensions of threat magnitude or cost, threat probability, and coping potential to levels that realistically reflect the threat or even slightly underestimate it. In mildly aversive situations, positive appraisal is easily achieved by a class of neuro-cognitive processes or mechanisms that we have termed “positive situation classification”, consisting in a comparison of a current situation with earlier, successfully managed situations or in the recurrence to positive cultural stereotypes that eventually lead to the relatively effortless activation of pre-existing positive appraisal patterns. However, in many or most aversive situations, negative appraisals are triggered automatically and are therefore largely unavoidable, presumably reflecting an evolutionarily determined preference for protection and defence. In such situations, positive appraisal and concomitant stress response regulation depend on the individual’s ability to positively reappraise (re-evaluate) a situation. Reappraisal processes/mechanisms can range from unconscious, automatic/effortless, implicit, non-verbal and non-volitional to conscious, effortful, explicit, verbal and volitional. They may reflect decreases in the actual threat value of a situation, for example in fear extinction, when a fear-conditioned stimulus (CS) that originally predicted threat (the unconditioned stimulus, UCS) is no longer followed by the UCS. Two other such “safety learning” processes are discrimination (e.g., between a threat-predictive CS+ and a non-predictive and hence non-dangerous CS−), and recovery after stressor termination. The function of these processes is to avoid unnecessary, costly stress responses. Another class of reappraisal processes changes the relative weighting of the negative and positive aspects present in any situation towards a more positive interpretation. One example is volitional (“cognitive”) reappraisal (Gross, 1998, 2001). Reappraisals do not necessarily have to occur at the time of stressor exposure, but may also be achieved in retrospect, thereby counteracting the consolidation or overgeneralization of aversive memories or generating competing positive memories. Finally, the positive adjustment of appraisals in strongly aversive situations (“reappraisal proper”) also requires a capacity to inhibit the interference resulting from competing negative appraisals and from the accompanying aversive emotional reactions (Kalisch et al., 2015). Hence, in addition to positive situation classification, positive reappraisal (proper) and interference inhibition are two other broad classes of neuro-cognitive processes whose efficiency and effectiveness together fashion an individual’s appraisal style. Via the ensuing shaping of stress responses, they ultimately determine the differences that individuals with comparable stressor exposure show in their long-term mental health outcomes. For a more extended discussion, see Kalisch et al. (2015).

### A battery for testing PASTOR

The efficiency and effectiveness of positive (re)appraisal processes can be indirectly inferred from the subjective-experiential, behavioral, physiological and neural reactions an individual shows to aversive stimuli, in particular when these stimuli change meaning towards the positive (i.e., terminate, or lose their threat predictiveness), when they carry ambiguous (both positive and negative) meaning, or when they are irrelevant (meaningless) distractors (Kalisch et al., 2015). In the present battery, we therefore assembled a range of tasks covering the three process classes described above (positive situation classification, positive reappraisal (proper) and interference inhibition). See Figure 1 for an overview. A purely behavioral stress test (Task 1) is intended to assess individual differences in the immediate stress response to a combined cognitive-sensory-emotional stressor (stress reactivity) as well as in stress response recovery when the stressor is over. All other tasks are fMRI tasks. A differential fear conditioning task (Task 3) assesses the reaction to a UCS-predictive CS+ as well as subjects’ discrimination of the CS+ from non-predictive CS-s, a case of safety learning. The subsequent extinction phase tests safety learning in the sense of ceasing to respond aversively to a no-longer predictive CS+. Whether the ensuing safety memory (extinction memory) is retrieved at re-confrontation with the CS+ and then inhibits the CS+ associated fear memory is tested the next day. In two tasks, subjects are explicitly invited to find and volitionally use a conscious reappraisal strategy (self-focused in Task 4, situation-focused in Task 5). Inhibition of goal-irrelevant emotional interference is tested in Task 6. A reward sensitivity test (Task 2) is included to index the functionality of the reward system as one major source of stress system inhibition, following opponent-systems theory (Dickinson & Pearce, 1977; Konorski, 1967; Nasser & McNally, 2012). These assessments are complemented by anatomical and diffusion-weighted MRI scans serving to measure brain structure and structural connectivity as well as several resting-state fMRI scans serving to measure functional connectivity and its modulation by stressor exposure as well as the dynamics of safety memory consolidation. Subjects also provide sociodemographic, life style, mental health and life history data and report on their appraisal and coping styles using established questionnaire instruments as well as a newly developed instrument specifically designed to index a generalized positive appraisal style. Validation data and results from this latter questionnaire will be presented in a separate publication.

**Figure 1.**
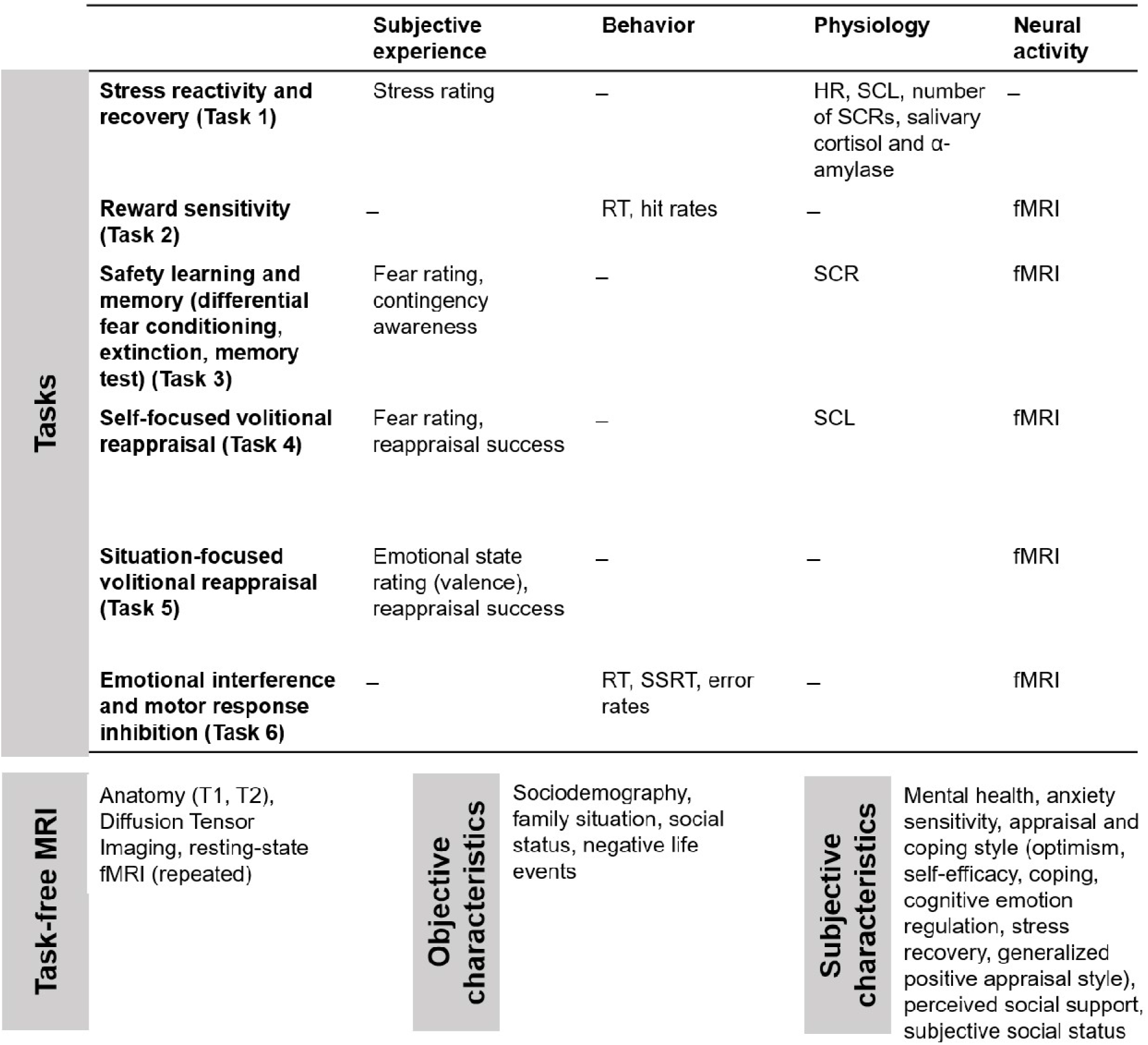
Overview of the test battery. HR, heart rate; SCL, skin conductance level; SCR, skin conductance response.

We assume that this multi-modal battery will tap into key positive (re)appraisal process, including their underpinnings in brain architecture, and can be analyzed using factor-analytic methods. Ensuing battery factors or components that can be interpreted to reflect (re)appraisal processes can then be tested in prospective-longitudinal settings for their capacity to predict whether individuals overcome a period of adversity in good mental health (i.e., whether they are resilient). More so, we expect that changes in relevant battery factors observed with repeated battery employment (such as several times during an extended period of life adversity or before and after a more circumscribed stressor exposure) will explain inter-and intra-individual variance in mental health time courses of stressed individuals, thereby revealing the key adaptations these individuals undergo to maintain or regain functioning (cf. Kalisch et al., 2015). For instance, it is conceivable that individuals start to adopt volitional reappraisal strategies when being confronted with challenging life situations, in an effort to improve their well-being and optimize their behavior, and with time also become increasingly efficient and effective in using these strategies, further reducing the likelihood of them developing mental problems. Similar scenarios can be imagined for other processes reflected in battery factors. Hence, the battery is also intended to reveal resilience-promoting training or plasticity effects.

We emphasize that the purpose of the battery is to measure the effectiveness/efficiency of neurobehavioral functions hypothesized to be important resilience mechanisms, i.e., to contribute to the long-term maintenance of mental health under adversity. The battery does explicitly not serve to measure resilience. Resilience as an outcome can only be measured in a longitudinal setting where individuals are exposed to real-life stressors and their ensuing mental health changes are put into relation with stressor exposure. Individuals who are less affected than other individuals experiencing comparable stressor exposure can then be classified as more resilient (Kalisch et al., 2017). In such a setting, the battery can be employed either once (e.g., at study inclusion, to ask whether the measured functions predict a resilient outcome) or repeatedly during the course of the study (to ask whether changes in the measured functions – in the sense of adaptation processes - relate to resilient outcome).

### The current report

A repeated employment of the battery is currently practiced in the prospective-longitudinal Mainz Resilience Project (MARP), where the battery is also complemented at each repetition with further measures of life history, hair cortisol, genetic background, epigenetic marks, immune factors, gut microbiome composition and others. A detailed description of the MARP study rationale and design as well as of the outcome-based measurement of resilience operationalized in MARP will be provided elsewhere. In the current report, we focus on an exhaustive description of the battery methods and procedures and provide group results for the different tasks and other battery elements (except brain anatomy, structural and resting-state connectivity) from a separate cross-sectional cohort of N=55 healthy young subjects, in which the battery was first applied for the purpose of testing its feasibility and usefulness (“discovery sample”). Resource limitations prohibited immediate repetition of the battery in this sample, so we cannot report on test-retest reliability. We here assess whether dropout rates are acceptable (to test feasibility) and whether the fMRI group-level task activations conceptually replicate results from previous studies with similar tasks, as derived from meta-analyses or reviews, to estimate the robustness and thereby usefulness of our battery tasks. In a follow-up publication, we will report on the exact replication of the group results in the first 103 MARP subjects at the time of their study inclusion (“replication sample”), to provide another indication of the battery’s robustness. On this basis, we can then proceed to find a factor solution for the battery in the discovery sample, attempt to confirm it in the replication sample, and then formulate hypotheses about the effects of relevant battery factors (potentially reflecting key resilience mechanisms) on resilience outcomes in the prospective-longitudinal MARP investigation.

## Method

### Participants

Fifty-six right-handed, healthy, Caucasian subjects with normal or corrected-to-normal vision and command of German volunteered in the study. Subjects were recruited via postings on notice boards on the campus and via social media channels. One male subject was excluded after study completion since his answers in one of our self-report instruments indicated potential presence of a mental disorder. All other subjects were free of psychiatric or neurological conditions according to their self-report and none took any neuro-pharmacological or psycho-pharmacological medication. One subject dropped out before MRI measurement because of claustrophobia. The remaining N=54 subjects (30 women) had a mean age of 25 years (age range: 18 to 31 years). The majority of our sample were university students (72.2%), one subject was still in secondary school (1.9%), 13.0% had a full-time job and 3.7% were unemployed.

Table 1 and Table 2 give further sample characteristics. All subjects gave written informed consent. The study was approved by the local ethics committee (Medical Board of Rhineland-Palatinate, Mainz, Germany) and conducted in accordance with the Declaration of Helsinki. Subjects were reimbursed with 90 € for their participation.

**Table 1.** Sample demographics

|  |  |  |  |  |
| --- | --- | --- | --- | --- |
| Country of birth | Germany (94.4) | other (5.6) |  |  |
| Mother tongue | German (98.1) | other (1.9) |  |  |
| Adoption | yes (1.9) | no (98.1) |  |  |
| Living situation in childhood | living with both parents (81.5) | living with one parent only (16.7) | alternatingly living with mother or father (1.9) |  |
| Father | alive (90.7) | dead (9.3) |  |  |
| Mother | alive (100) | dead (0) |  |  |
| Number of siblings | 0 (2) | 1 (42.9) | 2 (26.5) | >2 (16.3) |
| Twin | yes (1.9) | no (98.1) |  |  |
| Mental disease in family<sup>1</sup> | yes (31.5) | no (68.5) |  |  |
| Monthly household income | >2000 (17.1) | 1000-2000 (27.7) | 750-999 | <750 |
| Negative life events<sup>2</sup> | 6.8 ± 2.7 |  |  |  |

Percentages are given in brackets. <sup>1</sup>examples given: anxiety disorders, depression, schizophrenia, addiction to alcohol, illicit drugs or medicaments; <sup>2</sup>ad. from Caspi, Moffit, & Thorton (1996), translated to German, Likert scoring from 0-.

**Table 2.** Psychological characteristics of the sample

| Measure | *M* | *SD* |
| --- | --- | --- |
| Mental health (GHQ-28)<sup>1</sup> | 48.5 | 8.41 |
| Anxiety sensitivity (ASI-3)<sup>2</sup> | 19.1 | 10.5 |
| Stress recovery ability (BRS)<sup>3</sup> | 3.7 | 0.8 |
| Optimism (LOT-R)<sup>4</sup> | 15.4 | 4.5 |
| Self-efficacy (SWE)<sup>5</sup> | 30.1 | 4.2 |
| Coping style (BC)<sup>6</sup> |  |  |
| Self-distraction (BC1) | 5.6 | 1.3 |
| Denial (BC2) | 3.2 | 1.3 |
| Emotional support (BC3) | 5.9 | 1.5 |
| Behavioral disengagement (BC4) | 3.0 | 0.9 |
| Positive reframing (BC5) | 5.2 | 1.6 |
| Humor (BC6) | 4.2 | 1.8 |
| Active coping (BC7) | 5.7 | 1.3 |
| Substance use (BC8) | 2.4 | 0.9 |
| Instrumental support (BC9) | 5.6 | 1.6 |
| Living out emotions (BC10) | 4.3 | 1.3 |
| Planning (BC11) | 5.9 | 1.3 |
| Acceptance (BC12) | 6.0 | 1.5 |
| Self-blame (BC13) | 4.4 | 1.6 |
| Religion (BC14) | 3.1 | 1.7 |
| Cognitive emotion regulation (CERQ)<sup>7</sup> |  |  |
| Acceptance (CERQ1) | 7.7 | 1.9 |
| Rumination (CERQ2) | 6.2 | 2.0 |
| Positive reappraisal (CERQ3) | 7.3 | 2.0 |
| Self-blame (CERQ4) | 4.8 | 2.1 |
| Positive refocusing (CERQ5) | 4.7 | 1.7 |
| Catastrophizing (CERQ6) | 4.4 | 2.3 |
| Blaming others (CERQ7) | 3.6 | 1.6 |
| Planning (CERQ8) | 7.1 | 1.8 |
| Putting into perspective (CERQ9) | 6.7 | 2.1 |
| Distancing<sup>8</sup> | 5.3 | 2.1 |
| Social support (OSS-3)<sup>9</sup> | 10.6 | 1.8 |
| Subjective social status<sup>10</sup> |  |  |
| Country | 6.5 | 1.4 |
| Community | 7.3 | 1.3 |

N=54. <sup>1</sup>German version: Klaiberg, Schumacher, & Brähler (2004), original: Goldberg & Hillier (1979); <sup>2</sup>Kemper, Ziegler, & Taylor (2009), Taylor et al. (2007), N=53; <sup>3</sup>Brief Resilience Scale; Chmitorz, Wenzel, et al. 2018); Smith et al. (2008); <sup>4</sup>Glaesmer, Hoyer, Klotsche, & Herzberg (2008); Scheier & Carver (1985); <sup>5</sup>Schwarzer & Jerusalem (1999); <sup>6</sup>Knoll, Rieckmann, & Schwarzer (2005); Carver (1997); <sup>7</sup>Loch, Hiller, & Witthöft (2011); Garnefski, Kraaij, & Spinhoven (2001); <sup>8</sup>Two items reflecting distancing/detachment were added to the original questionnaire (English translation: “I try to observe the situation from a detached perspective, like from outside.”, “I try to distance myself from the situation and my feelings.”); <sup>9</sup>ad. from Dalgard, Bjork, & Tambs (1995), translated to German. <sup>10</sup>Euteneuer, Süßenbach, Schäfer, & Rief (2014), Adler, Epel, Castellazzo, & Ickovics (2000), community: N=53; M, mean; SD, standard deviation. Results from a newly developed generalized positive appraisal style questionnaire are not reported here.

### Procedure

The study was conducted in 2015. All tests took place at University Medical Center Mainz. Subjects were invited to one behavioral testing session (visit 1) and two MRI sessions (visits 2 and 3). Visit 1 and visit 2 were maximally four weeks apart, while visits 2 and 3 were on two consecutive days. See Table 3 for an overview of the testing schedule. The main experimenter was female, supplemented by one to two assistants (male or female).

**Table 3.** Procedure

| Visit | Schedule | Duration (min) |
| --- | --- | --- |
| Behavioral testing session (visit 1) | Study information, consent | 30 |
|  | Questionnaires | 45 |
|  | Task 1: Stress reactivity and recovery |  |
|  | - Set-up | 10 |
|  | - Rest phase (baseline) | 6 |
|  | - Break (instructions, saliva sample 1) | 10 |
|  | - Stress phase | 5 |
|  | - Break (saliva sample 2) | 10 |
|  | - Rest phase (recovery) | 6 |
|  | Instructions on and practice of fMRI tasks, psychoeducation on the impact of thoughts on emotion, saliva samples 3-5 | 60 |
|  | Confirmation of MRI suitability by study physician | 15 |
|  |  | Total: 200 |
| First MRI session (visit 2) | Set-up (instructions, placement in scanner, electrode attachment, pain calibration, localizer scans) | 20 |
|  | Resting-state fMRI, run 1 (Baseline) | 6 |
|  | Task 2 (fMRI): Reward sensitivity | 8-9<sup>1</sup> |
|  | Calibration of pain stimulus for Task 3 | 10 |
|  | Task 3 (fMRI), run 1: Differential fear conditioning (CD) (+Habituation, HB) | 10.5 (+1.5) |
|  | Resting-state fMRI, run 2 | 6 |
|  | Task 3 (fMRI), run 2: Extinction (EX) | 6 |
|  | Resting-state fMRI, run 3 | 6 |
|  | Structural MRI: MPRAGE (T1), T2, DTI | 20 |
|  | Resting-state fMRI, run 4 | 6 |
|  |  | Total: 100 |
| Second MRI session (visit 3) | Set-up (instructions, placement in scanner, electrode attachment, localizer scans) | 20 |
|  | Task 3 (fMRI), run 3: Memory test (spontaneous recovery, SR; renewal, RN) | 9 |
|  | Instructions on self-focused reappraisal and calibration of pain stimulus for Task 4 | 10 |
|  | Task 4 (fMRI): Self-focused volitional reappraisal | 10 |
|  | Task 5 (fMRI): Situation-focused volitional reappraisal | 14 |
|  | Task 6 (fMRI), runs 1-3: Emotional interference and motor response inhibition | 24 (3 x 8) |
|  |  | Total: 90 |

<sup>1</sup>Exact duration varies due to an adaptive algorithm that set target durations according to subject performance. MPRAGE, Magnetization-Prepared Rapid Acquisition Gradient-Echo sequence; DTI, Diffusion Tensor Imaging.

#### Behavioral testing session (visit 1)

After filling in the questionnaires, subjects performed Task 1 (stress reactivity and recovery; for details on the tasks, see the description of the tasks below). Subjects then received instructions for the fMRI tasks to be presented at visits 2 and 3 and were given the opportunity to practice Tasks 2, 3, 5 and 6. To prepare volitional reappraisal at visit 3 (Tasks 4 and 5), subjects received a psychoeducation about the impact of thoughts on emotion and about emotion-related brain regions. Subjects received a short paper form of the task instructions to take home. At the end of visit 1, MRI suitability was confirmed for each subject individually by a study physician.

#### MRI sessions (visits 2 and 3)

Visits 2 and 3 each lasted approximately 1.5 to 2 h and were conducted in the afternoon and evening (between 2 and 6 pm). At the beginning of each visit and before being placed in the MRI scanner, subjects were again instructed on the tasks.

During visit 2, subjects performed Task 2, assessing reward sensitivity, which lasted 8 to 9 minutes depending on subject performance. To prepare Task 3 on safety learning, the current amplitude for electro-tactile stimulation was then determined in a pain stimulus calibration procedure. This was followed by the first two runs of Task 3, involving differential fear conditioning, which was directly preceded by a 1.5-minutes habituation phase (HB+CD, task run 1, 10.5 min), and extinction (EX, task run 2, 6 min). Four 6-minutes resting-state fMRI scans were collected: at baseline (before Task 2), immediately after fear conditioning (after Task 3, run1), immediately after extinction (after Task 3, run 2) and at the end of visit 2. For these resting-state scans, subjects were instructed to remain eyes open, to fixate the white cross on the black background in the center of the screen and not to think of anything particular. At visit 2, we also acquired structural MRI scans (MPRAGE/T1, T2, DTI; seeTable 3). During the acquisition of the structural scans (20 min), subjects were presented with a movie clip that showed moving shapes, in order to reduce subject movement (Vanderwal, Kelly, Eilbott, Mayes, & Castellanos, 2015).

At visit 3, approximately 24 h later, subjects’ fear and extinction memories were tested (Task 3, run 3, 9 min). Each memory test (spontaneous recovery [SR], renewal [RN]) lasted 4.5 min. Memory tests were conducted directly following each other in one experimental run. Thereafter, subjects received instructions on self-focused volitional reappraisal on the screen while lying in the scanner (Task 4), and the to-be-used pain stimulus was calibrated. Subjects then had to apply self-focused (Task 4, 10 min) and situation-focused reappraisal (Task 5, 14 min), in order to deliberately regulate their emotional reactions to different affective stimuli. The second MRI session ended with three runs of a combined emotional interference and motor response inhibition task (Task 6, 24 min). During both MRI sessions, subjects never left the scanner.

### Tasks

All tasks were programmed in Presentation software (NBS, San Francisco, USA). Tasks 1, 5 and 6 all make use of emotional pictures; we assured that each picture was only used in one of the tasks. Note that the repeated use of the battery in longitudinal settings such as MARP involves different stimulus sets and trial orders at battery repetition, to reduce potential learning effects. These modifications will be described in the context of a future MARP study protocol.

#### Task 1: stress reactivity and recovery

To induce stress, a slightly modified version of the Mannheim Multicomponent Stress Task (MMST; Reinhardt, Schmahl, Wüst, & Bohus, 2012) was applied (Figure 2). The MMST consists of multiple stressors of different modalities: a cognitive, an emotional, an acoustic and a motivational stressor and was here extended to also include a social-evaluative stress component. As cognitive stressor, subjects had to perform a mental arithmetic task (PASAT-C; Lejuz, Kahler, & Brown, 2003), consisting in summing two sequentially presented numbers in the range of 0 to 20. Numbers were presented for 250 ms each. Time latency between numbers was reduced from 3 s to 2 s in the second half of the task. Answers were given via mouse by marking the result on a numerical keypad presented in the lower part of an emotional picture on the screen. Subjects received online performance feedback on the total number of correct answers shown below the keypad. The PASAT-C lasted approximately 3.5 min. As emotional stressor, negative emotional photographs were used. Reinhardt et al. (2012) had selected 44 negative and nine positive pictures from the International Affective Picture Set (IAPS; Lang, Bradley, & Cuthbert, 2008) and from the internet. Negative images were fear- and disgust-related, while positive images showed animals and babies. Negative images were presented in blocks of five, always followed by one positive picture, to avoid habituation. Picture duration was 5 s for negative and 3 s for positive pictures; there was no delay between the presentations of two consecutive pictures and between blocks of pictures. To direct subjects’ attention to the pictures, the task started with a 1-minute exposure to thirteen pictures (two picture blocks + one additional negative picture) in the absence of the cognitive stressor. To further ensure attention, subjects were asked to detect pictures that were presented twice during this phase of the task and to indicate this using the computer mouse after the phase (10 s). For the rest of the task, the images were used as background pictures for the PASAT-C. As acoustic stressor, white noise and 60 randomly timed explosion sounds were presented over headphones. During the first minute of the task, the noise volume was kept constant (75 dB), while with the onset of the PASAT-C the volume started to continuously rise to a maximum of 93 dB, and additionally explosion sounds were given. In the original study, explosion sounds indicated incorrect answers. However, since we wanted to keep the sensory input equal for all subjects, we decided to uncouple explosion sounds from performance. As motivational stressor, subjects were told that their reimbursement might be reduced by 10 € if they performed too poorly. Performance-related punishment on reimbursement was never actualized. We added an element of social evaluation to the original design by giving three negative, social-comparative performance feedbacks on the screen, one after the first minute of the PASAT-C, one after 2.5 min and one at the end of the task (see Figure 2). The order of feedbacks was identical across subjects.

**Figure 2.**
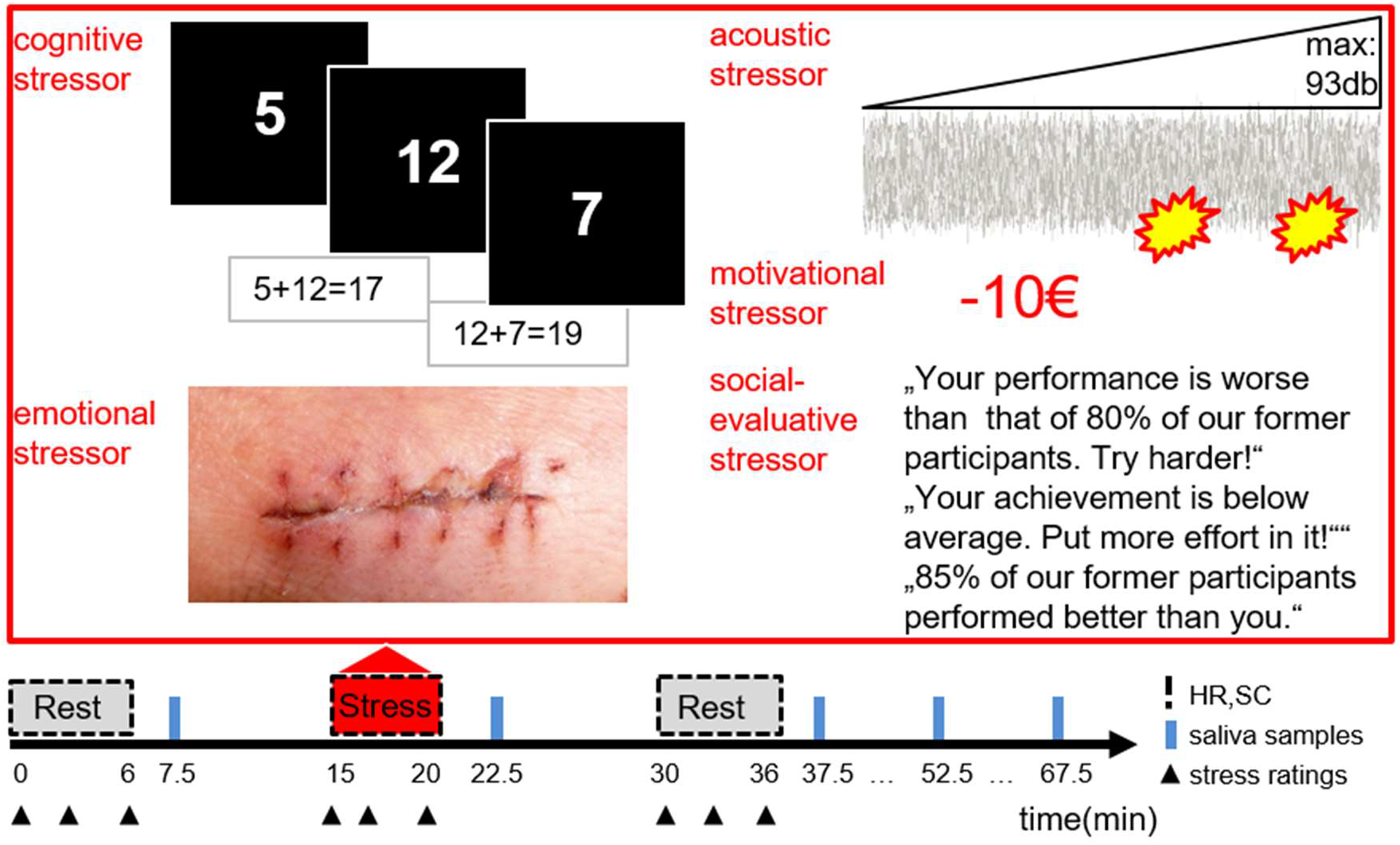
Design of Task 1: Stress reactivity and recovery. The task consisted of three experimental phases: rest (baseline, 5.75 min), stress (5.15 min), and rest (recovery, 5.75 min), during which continuous heart rate (HR) and skin conductance (SC) as well as discrete stress ratings (triangles) were recorded. Saliva samples (blue bars) were taken in the intervals and after the phases. To stress subjects, five different types of stressors were applied. As cognitive stressor, subjects had to perform a mental arithmetic task displayed on a background of negative pictures, which served as an additional emotional stressor. As acoustic stressor, increasing white noise and explosion sounds were given. As motivational stressor, subjects were threatened by potential reduction of their reimbursement. As social-evaluative stressor, negative social evaluative performance feedback was presented. The image displayed for the emotional stressor is a placeholder for the category of disgust-related pictures used as emotional stressors (kindly provided by Pavel Ševela / Wikimedia Commons).

Stressor exposure lasted altogether 5.15 min and was preceded and followed by two 5.75-minutes resting phases (baseline, recovery), during which subjects had to fixate a white cross on a black computer screen and to give three stress ratings (Figure 2). In the 10-minutes interval between the baseline rest and the stressor exposure phases, subjects received instructions on the upcoming task, and a first saliva sample was collected at 1.5 min into the interval. In the 10-minutes interval between the stressor exposure and the recovery rest phases, a second saliva sample was collected at 2.5 min, and subjects filled in a questionnaire on the preceding stress experience. Three more saliva samples were collected during the subsequent phase of instructions on the future fMRI tasks to be performed at visits 2 and 3 (see Figure 2). Sampling started 1.5 min after the end of the recovery rest phase and continued in intervals of 15 min. Hence, a total of five saliva samples were collected throughout the task. The time between successive samples was always approximately 15 min.

We assessed stress ratings, heart rate (HR), skin conductance level (SCL), and number of skin conductance responses (nSCRs) during the three experimental phases as well as salivary cortisol and salivary α-amylase levels from the samples collected after the phases. For the stress ratings, subjects were asked to indicate how stressed they felt in this moment on a visual analog scale (VAS) ranging from 0 (not stressed at all) to 9 (very stressed), presented on the screen for 11 s. Ratings were taken at the beginning and at the end of each experimental phase. One additional rating per phase was taken at 2.5 min (baseline, recovery) and 1 min (stressor exposure) into the respective phase. The stress experience questionnaire applied after the stress phase asked subjects to name maximally three components of the test that were most stressful to them.

#### Task 2: reward sensitivity

The anticipation and receipt of gains and losses reliably activates structures of the reward system in the brain (Knutson, Adams, Fong, & Hommer, 2001; Wu, Samanez-Larkin, Katovich, & Knutson, 2014). We applied a shortened and simplified version of the monetary incentive delay (MID) task as described by Wu et al. (2014). There were five different experimental conditions of varying incentive magnitude (+3 €, +0.5 €, ±0 €, −0.5 €, −3 €). We used one condition less than Wu et al. (2014) who distinguished between +0 € and −0 € amounts. The stimulus material consisted of the numeric Gain cues (+3 €, +0.5 €) presented within a circle, the Loss cues (−3 €, −0.5 €) presented within a square, the Zero cue presented within a diamond and the target, which was a white star on black background (see Figure 3). Subjects had practiced the task during the behavioral session (visit 1), wherein they did not gamble for money, but could instead increase a numeric score.

**Figure 3.**
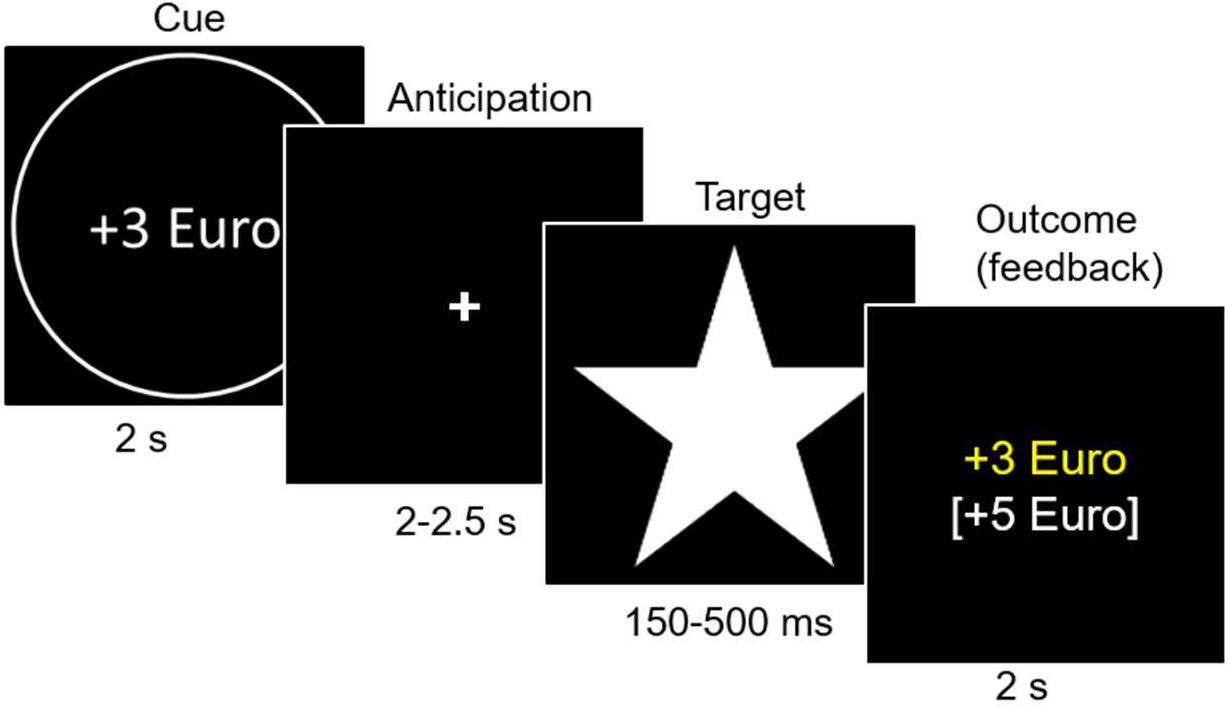
Design of Task 2: Reward sensitivity. Each trial started with a 2-seconds cue indicating the incentive condition (+3 €, +0.5 €, ±0 €, −0.5 €, −3 €), followed by an anticipation phase of 2 to 2.5 s. Subjects had to press the right button when the target (a white star) appeared on the screen. Target duration was adapted based on past task performance in a range from 150 and to 500 ms. Each trial ended with a 2-seconds numeric feedback on subjects’ trial outcome (gained amount in yellow or lost amount in red) and overall outcome (accumulated total amount). As an example, a correct +3 € trial is shown.

The task started with a 14-seconds task instruction. Each trial lasted approximately 6 to 7 s and started with the presentation of a numeric cue for 2 s, indicating the amount of money subjects could gain or avoid losing. Each cue was followed by a 2 to 2.5-seconds anticipation phase, during which a white fixation cross was on the black screen until target onset. Duration of the target ranged from 150 to 500 ms depending on past task performance. Subjects’ task was to press a button on a response pad with their right middle finger as soon as the target appeared, in order to gain or to avoid losing the indicated amount. Each trial ended with a 2-seconds numeric feedback on subjects’ trial outcome (gained amount in yellow or lost amount in red) and the overall outcome (accumulated total amount).

The mean inter-trial-interval (ITI) was 4 s with a range of 2 to 6 s. To assure that the experience of reward did not differ between subjects depending on task performance, we tried to achieve a hit rate of 66% within each subject and condition. We therefore applied an algorithm that adaptively changed target duration for a given subject within each condition based on past performance. If the subjects’ hit rate was below 66%, target duration was increased by 25 ms; else, it was reduced by 25 ms. Initial target duration was entered based on mean target duration in the practice session at visit 1, but had to be at least 300 ms.

Due to time restrictions, we limited our trial number to 9 trials per condition, so that the total trial number was 45. Two pseudo-random trial orders were used in which maximally two consecutive trials were of the same kind. The second trial order was created by interchanging the Gain and Loss trials of the first trial order, to exclude ordering effects. Subjects were instructed that their final overall outcome would be paid out to them. The average achieved total score was 9.78 € (SD=5.13 €). Nonetheless, every subject received the same amount (20 €) on top of their reimbursement, to keep up motivation for the rest of fMRI testing in case of low performance.

Reaction times (RT) and hit rates were used as behavioral outcome measures.

#### Task 3: safety learning and memory

Task 3 investigated safety learning by means of differential Pavlovian fear conditioning (run 1, CD), extinction (run 2, EX) and a memory retrieval test (run 3) that involved spontaneous recovery (SR) and renewal (RN) (Kalisch, 2006; Milad et al., 2007)(see Figure 4). Testing was conducted on two consecutive days, to separate learning and memory consolidation (visit 2) from memory retrieval (visit 3). Subjects had practiced the trial-by-trial fear ratings and the post-run contingency questions in the behavioral session (visit 1).

**Figure 4.**
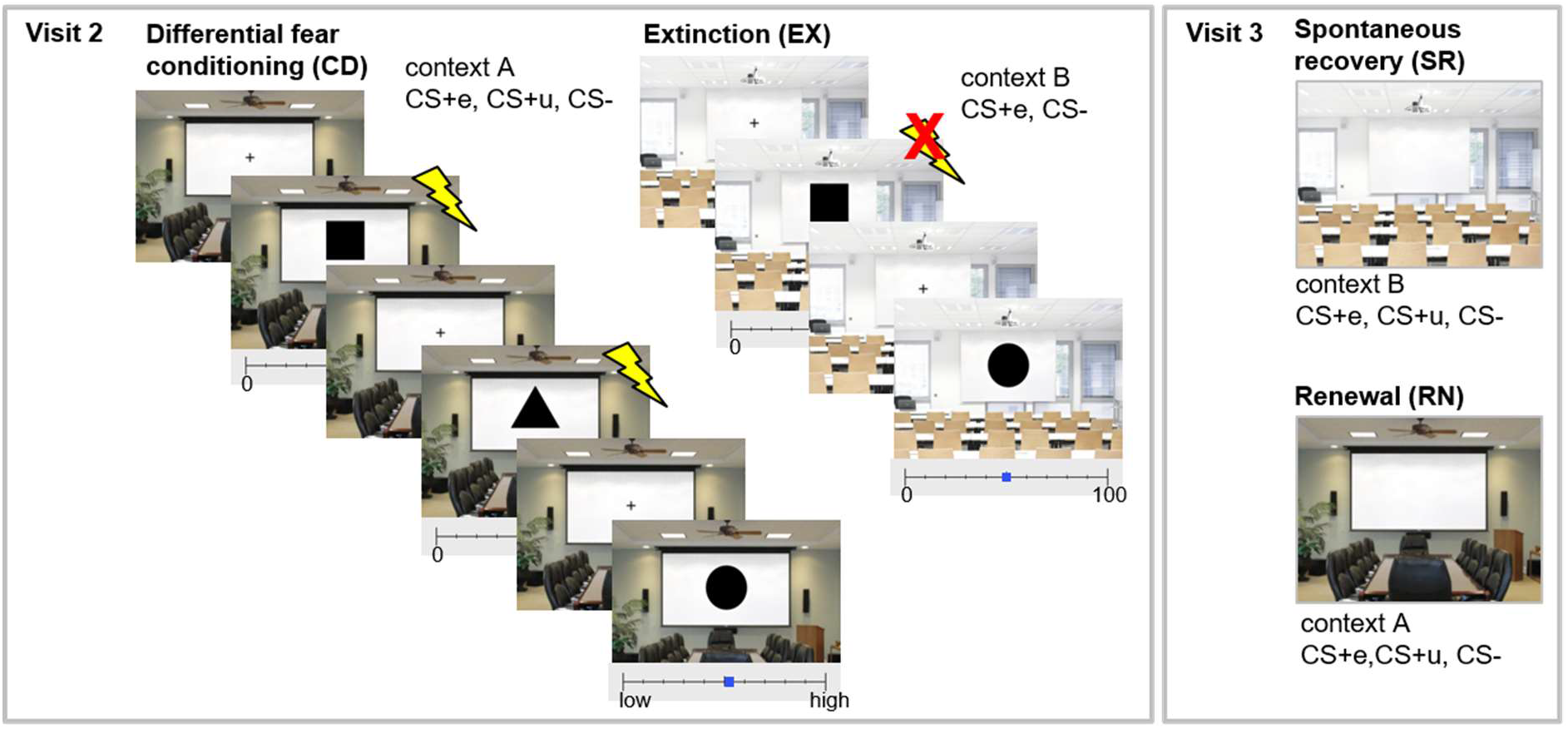
Design of Task 3: Safety learning and memory. Differential fear conditioning (CD) was conducted in context A (here: conference room) with two CS+s (CS+e, Cs+u) and one CS- (black symbols on screen). CS+s were coupled with the UCS in 100% of trials, while the CS− was always safe. Extinction (EX) took place in context B (here: seminar room), using only one of the CS+s (CS+e). No UCS was administered during extinction. On the following day, two memory retrieval tests were performed, again using all three CSs. Spontaneous recovery (SR) was tested in the extinction context B and renewal (RN) in the conditioning context A. Each CS trial lasted 6 s. During the first 4.5 s of a trial, subjects rated their fear of receiving a pain stimulus using a VAS at the bottom of the screen. ITIs lasted 9 to 15 s.

Three black geometric symbols (triangle, circle, square) served as conditioned stimuli (CSs). The CSs were super-imposed on background pictures of either a seminar room or a conference room, which served as contexts A and B. Both pictures contained a screen in the middle, so that the CS could be presented naturalistically as being projected onto the screen. The assignment of stimuli and contexts was counterbalanced across subjects.

A painful electrical stimulus delivered on the right dorsal hand was employed as the aversive unconditioned stimulus (UCS). During conditioning, two CSs were coupled with the UCS in 100% of trials, to become CS+s. The third CS (CS−) was never followed by the UCS and therefore safe. Safety learning in a differential conditioning paradigm consists in discriminating the safety of the CS− from the threat associated with the CS+. Accordingly, subjects should develop a safety memory associated with the CS− and fear memories associated with the two CS+s. During extinction, only one CS+ (henceforth termed CS+e) as well as the CS− were presented. There was no UCS. Safety learning here consists in recognizing the safety of the CS+e and in developing an associated safety memory (or “extinction memory”). During the memory test one day later, we presented all three CSs, assuming that the comparison of an extinguished CS+ (i.e., CS+e) with an unextinguished CS (termed CS+u) reveals genuine information about the CS+e associated safety memory, expressed in reduced CRs to the CS+e relative to the CS+u. Also, the comparison of CS− responses with the CS+s can reveal information about learned safety.

Conditioning was conducted in context A, whereas extinction was conducted in B. During the memory test, spontaneous recovery (SR) of CRs was tested by presenting the three CSs in the same context as extinction (B). Fear renewal (RN) was then tested in the original conditioning context (A). The latter situation normally favors fear memory retrieval, due to threat associations of the context, and therefore represents a more demanding test for safety memory retrieval than spontaneous recovery.

Directly before conditioning, the UCS current amplitude was determined in a calibration procedure to be maximally painful, but tolerable (M=6 mA, SD=3.3 mA, range: 2 to 20 mA, N=51). Subjects rated the painfulness of the chosen UCS on a scale from 0 (not painful) to 10 (maximally imaginable focal pain from such type of electrode) (M=8.2, SD=1, range: 6 to 10). Subjects were then instructed, first via headphones, then in written form on the screen, that they might or might not receive electric stimulation during the following experimental phases, that electric stimulation would only be administered at the offset of a symbol, and that they should be attentive. These instructions were repeated before each run. In all runs, CS duration was always 6 s and subjects had to rate their fear of a pain stimulus on a VAS at the bottom of the screen that ranged from 0 (no fear) to 100 (high fear). The scale froze after 4.5 s, after which the rating could no longer be changed, and remained on the screen until the next trial. The average ITI was 12 s (range 9 to 15 s), except during habituation (2 s). During ITIs, subjects saw a black cross on the background picture (A or B). The background picture was continuously shown throughout the respective run to provide a global context. After each run, subjects had to answer a range of contingency questions presented on the screen, using the response pad^1^.

The first run started with a habituation phase of 1.5 min, in which every CS was shown twice on each background picture. In the immediately following conditioning phase (9 min) each CS (CS+e, CS+u, CS−) was shown 8 times. UCSs were delivered at CS+ offset. After conditioning, subjects had to indicate again the level of pain experienced during UCSs (0-10) (M=7.7, SD=1.45, range: 1 to 10). Pain ratings habituated slightly relative to before conditioning (t=2.18, p=0.03). During the extinction run, the CS+e and the CS− were shown 8 times each; there was no UCS. Before the memory tests at visit 3 the next day (run 3), the pain electrode was again attached to the hand and subjects were asked to remind themselves of what they had learned in the runs on the previous day. During the two memory retrieval tests (spontaneous recovery, renewal), all three CSs were shown again, 4 times each. Each retrieval test lasted 4.5 min. Renewal directly followed upon spontaneous recovery without any interruption. Concerning trial order, we used two groups to balance out possible order effects for the two CS+s. Additionally, there were two possible trial orders for CS+e and CS− during extinction. Trial order was construed such that not more than two stimuli of one kind would follow each other.

Behavioral and physiological outcome measures were SCRs, fear ratings and contingency responses.

#### Task 4: self-focused volitional reappraisal

Distancing or detaching oneself from a threatening situation or from one’s own thoughts or feelings is a way to volitionally reappraise (reduce) the self-relevance of salient external or internal stimuli and thereby to dampen emotional reactions (Kalisch et al., 2005). We employed an adapted version of the task used by Paret et al. (2011). As preparation, subjects had received psychoeducation on the influence of thoughts on emotion and on brain regions involved in emotion and emotion regulation at visit 1.

We used a classical instructed fear paradigm (also known as “anticipatory anxiety” or “threat of shock”), which consisted in fore-warning subjects that they might receive a painful electric stimulus at any time during a trial (Threat condition, T). In a control condition (No Threat, NT), subjects were told they would not be stimulated during the trial. In a fully balanced, two-by-two factorial design, subjects either employed self-focused reappraisal (Reappraisal condition, R) or not (No Reappraisal comparison condition, NR). The stimulus material consisted in a yellow triangle with a black exclamation mark on black background that served as threat cue (T) and a blue triangle with a black minus on black background that signaled safety (NT) (see *Figure 5*).

**Figure 5.**
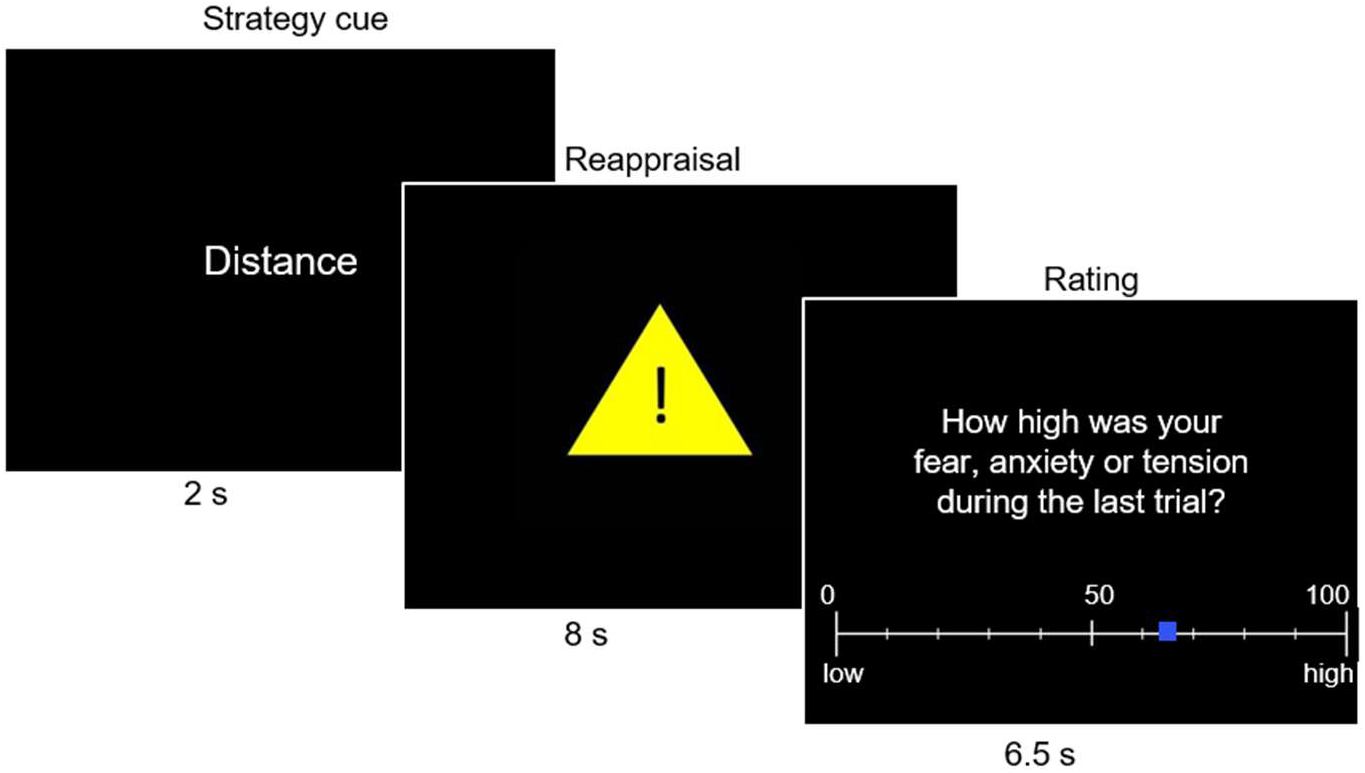
Design of Task 4: Self-focused volitional reappraisal. Subjects had to either downregulate their tension by distancing from the experimental situation (Reappraisal or R trials) or not (No Reappraisal or NR trials). Each trial started with a 2-seconds strategy cue word (“distance” in R or “leave” in NR trials), followed by an 8-seconds threat (exclamation mark in yellow triangle for threat trials, T) or no-threat cue (minus in blue triangle for No Threat trials, NT), during which the strategy had to be applied. Each trial ended with a rating of fear, anxiety and tension from 0 (low fear) to 100 (high fear) for 6.5 s. As an example, a T/R trial is shown.

Task instructions were given in written form on the screen while subjects were lying in the scanner. Subjects were reminded of the psychoeducation at visit 1 and told they should use their thoughts to reduce tension. Specifically, subjects were asked to take the perspective of a neutral and detached observer. To this purpose, they were shown four short self-statements describing how one can distance oneself from a situation^2^. After assuring subjects had understood the statements, they were asked to choose one statement and to repeat it five times in their minds, to be able to properly recall it during the task. To further enhance commitment, subjects were told that they might later be asked to recall the phrase. The chosen sentence was assessed verbally after the task.

To prepare the instructed fear task, we applied a pain stimulus calibration procedure. We started with the current amplitude used in Task 3 and repeated the calibration procedure (final amplitude: M=8.1 mA, SD=6.3 mA, range: 2 to 42 mA; final pain rating: M=8.5, SD=0.8, range: 6 to 10). In a second step, subjects were presented with at least three countdowns from 8 to 0 while expecting to receive a pain stimulus at any time during the countdown with a probability of 25%. An actual stimulation occurred in the second countdown. Current amplitude was considered appropriate when the fear of receiving such a stimulus was rated with 50 or higher on a scale from 0 (no fear) to 100 (high level of fear), as assessed verbally after the countdown (final rating: M=68, SD=16, range: 30 to 100). If the fear level was too low, either the current amplitude was increased, repeating the first step of calibration, or another countdown with stimulation was applied.

Subjects were then instructed that, during the task, they would receive a pain stimulus upon presentation of the yellow triangle with the exclamation mark in 25% of cases at any time during the trial (T trials), while the blue triangle with the minus sign would signal safety (NT trials). When seeing the strategy cue word “Distanzieren” (“distance”) on the screen, they should apply the distancing strategy during the following trial, with the help of the chosen self-statement (R trials). When seeing “Belassen” (“leave” or “do not regulate”), they should not apply the strategy (NR trials).

At the beginning of the task, the instructions were again presented for 40 s, in written form on the screen. In the task, there were a total of 20 trials, 5 per condition (NT/NR, NT/R, T/NR, T/R). Each trial started with the 2-seconds strategy cue word, followed by an 8- seconds presentation of the T or NT cue, during which subjects were to apply the strategy (see *Figure 5*). Within 6.5 s after cue offset, subjects had to rate their fear, anxiety and tension during the preceding trial on a VAS from 0 (no fear) to 100 (high fear) presented on the screen. Mean ITI was 7 s (range: 5 to 9 s). No pain stimulation was given in any of the trials. Trial order was pseudo-random, such that maximally two consecutive trials contained the same level of any of the two experimental factors. Two trial orders were used, interchanging R and NR.

We collected fear ratings and SCL as behavioral and physiological outcome measures. Perceived success of strategy use was assessed after the task via VAS from 0 (not successfully applied) to 100 (successfully applied).

#### Task 5: situation-focused volitional reappraisal

Emotional responses can volitionally be changed by reinterpreting or reframing the meaning of an emotional situation. In Task 5, subjects had to apply such a situation-focused form of reappraisal of emotional pictures, to explicitly achieve a more positive emotional state (see Figure 6). The paradigm is an adaptation of a previous design (Kanske, Heissler, Schonfelder, Bongers, & Wessa, 2011).

**Figure 6.**
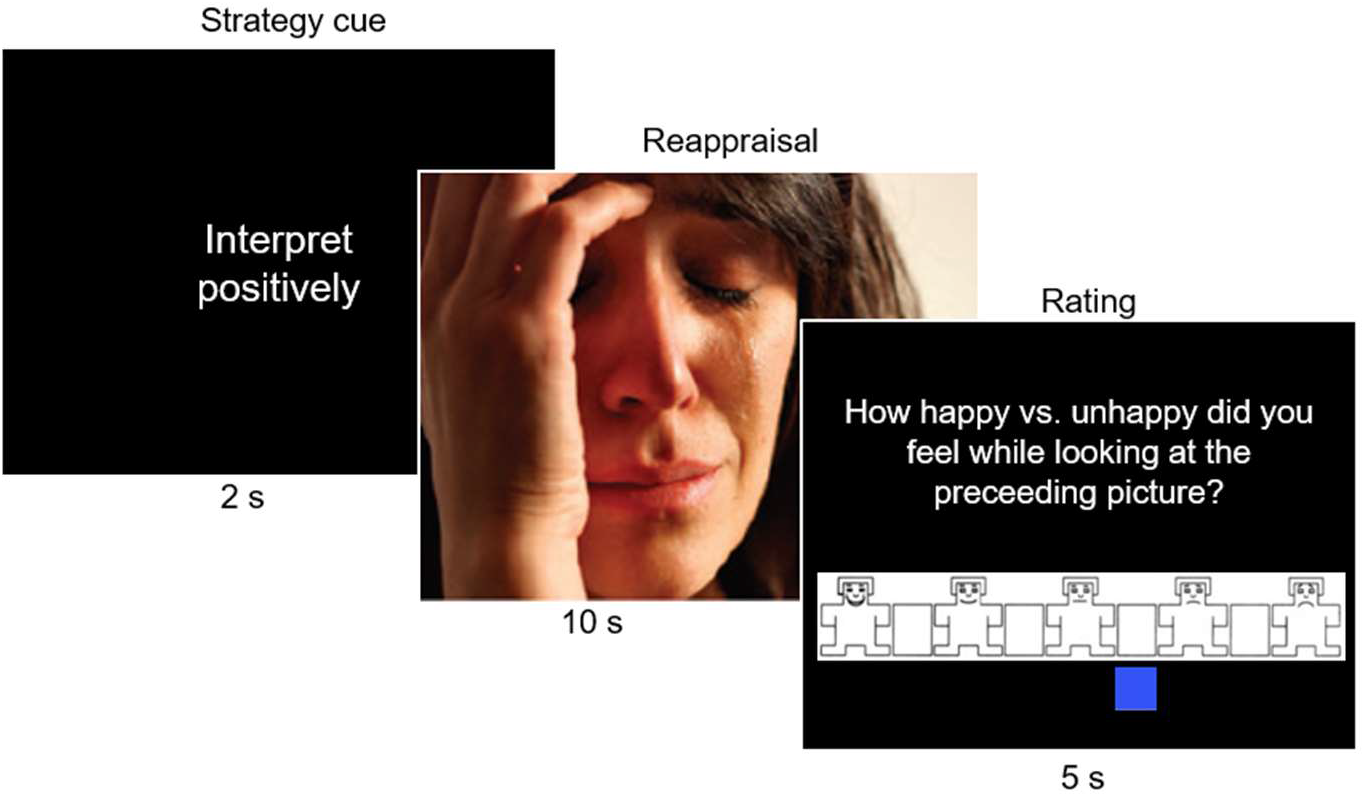
Design of Task 5: Situation-focused volitional reappraisal. Subjects had to either make their emotional state more positive by positively reinterpreting a depicted situation (Reappraisal or R trials) or not (No Reappraisal or NR trials). Each trial started with a 2-seconds strategy cue word (“interpret positively” in R or “view image” in NR trials), followed by a 10-seconds presentation of a positive (Pos), neutral (Neu) or negative (Neg) picture, during which the strategy was to be applied. Each trial ended with an emotional state rating from 0 to 5 for 5 s. As an example, a Neg/R trial is shown. The depicted image is retrieved from the EmoPicS (Wessa et al., 2010) and was already published in Kanske et al. (2011). It is used as a placeholder for emotional images here, but is not part of the stimulus set of Task 5.

We used a fully balanced, three-by-two factorial design, combining three types of picture valences (positive [Pos], neutral [Neu], negative [Neg]) with either situation-focused reappraisal (R) or viewing the image as a control condition (No Reappraisal, NR). Emotional pictures were selected from the EmoPicS (Wessa et al., 2010) and IAPS (Lang et al., 2008) picture sets, based on normative ratings of valence and arousal^3^. We only considered images of humans and social situations, excluding erotic contents to avoid gender biases. Positive pictures had high valence (>6.5) and were highly arousing (>4), negative pictures had low valence (<3.5) and were equally arousing (>4); neutral images had moderate valence (range: 4.5 to 5.5) and were only mildly arousing (<2.5). We created two sets of pictures for each valence matched by content and normative ratings. Each picture was only used once, to avoid habituation or transfer of emotion across conditions.

The reappraisal strategy in this task was to find a positive interpretation for a depicted situation by either focusing on positive aspects of the situation or by imagining a positive outcome. For example, when seeing an injured person, subjects could either think about an ambulance that was depicted in the background or they could imagine that the person would recover quickly. The control condition was to simply view the image and take in the impressions. Subjects had received psychoeducation on emotion regulation and had practiced situation-focused reappraisal at visit 1. To assure compliance, subjects provided literal picture interpretations for the last image of each valence shown in the reappraisal condition at the end of the practice at visit 1 via keyboard (results not reported here).

Upon start of the task, instructions were given, in written form on the screen for 40 s. Each trial began with a 2-seconds presentation of a strategy cue word on reappraisal, followed by 10-seconds presentation of a picture. The cue word “Positiv Interpretieren” (“interpret positively”) signaled R trials, while the cue word “Bild ansehen” (“view image”) signaled NR trials. The total number of trials was 30. All trials ended with an emotional state rating (5 s), in which subjects indicated how happy they felt while watching the picture on a scale from 1 (very happy) to 9 (very unhappy) with the IAPS self-assessment manikin scale (Lang et al., 2008). Emotional state ratings were recoded, so that high values indeed indicated high valence. ITIs were randomly generated between 5 and 9 s, with a mean ITI of 7.5 s. A white fixation cross on black background was presented during the ITIs. Picture set assignment was counterbalanced across subjects. We used two pseudorandom trial orders with inverse order for the NR and R trials. We ensured that there were no more than two R or NR trials or two trials of equal valence following each other.

Our behavioral outcome measures were emotional state ratings and post-experimental ratings of success of strategy use on a VAS from 0 (not successfully applied) to 100 (successfully applied).

#### Task 6: emotional interference and motor response inhibition

The Hybrid Response Inhibition task (HRI; Sebastian et al., 2013) was modified to test the inhibition of the interference induced by emotional picture stimuli with performance on a motor response inhibition task (Figure 7). The HRI is a paradigm that allows for testing subcomponents of motor response inhibition within the same paradigm with the same stimulus material. Two HRI-subtasks were included in Task 6: the Simon task, capturing spatial interference inhibition, and the stop signal task, assessing action cancellation, both in comparison to a simple go task as control condition (i.e., a two-choice response time task). The stimulus material for both tasks consisted of a white fixation cross on black background and a white arrow positioned either on the left or right side of the fixation cross. Fixation cross and arrow were encircled by an either white or blue ellipse. The arrow pointed either to the left or the right side. During the task, subjects had to indicate the pointing direction of the arrow by button press and to suppress responses if the ellipse turned blue (stop signal). 50% of the trials were congruent (Con Go), 25% incongruent (Incon Go) and 25% stop trials, which were all congruent trials. During congruent go trials, there was no spatial interference: the position of the arrow relative to the fixation cross corresponded to the pointing direction of the arrow. During incongruent go trials, the position of the arrow on the screen was incongruent with its pointing direction (e.g., a right pointing arrow was displayed left to the fixation cross), inducing spatial interference. In stop trials, the white ellipse turned blue after a stop signal delay (SSD), requiring subjects to cancel their prepared or ongoing action.

**Figure 7.**
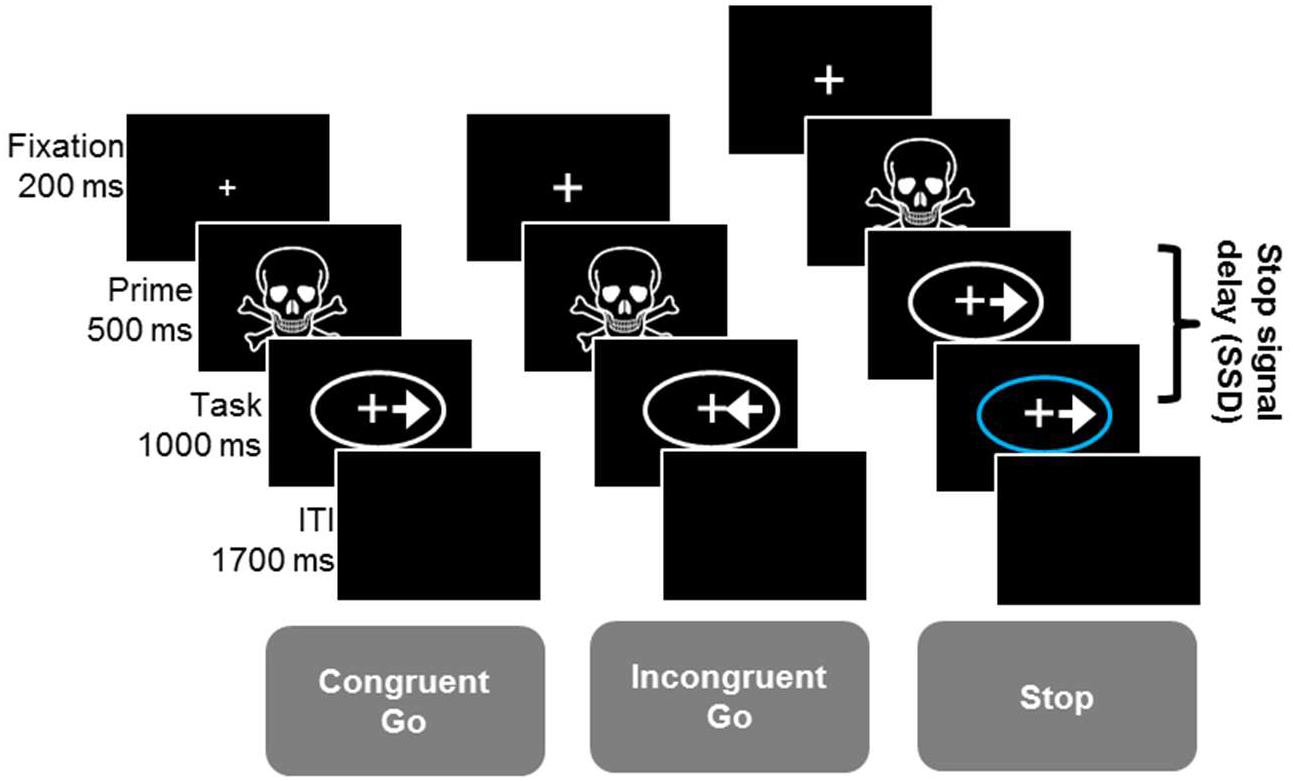
Design of Task 6: Emotional interference and motor response inhibition. Subjects had to indicate the pointing direction of the arrow by button press. During congruent go (Con Go) trials, pointing direction and position of the arrow relative to the fixation cross corresponded to each other, while in incongruent go (Incon Go) trials, pointing direction and spatial position of the arrow were incongruent with each other (Simon task testing spatial interference inhibition). In stop trials, the white ellipse turned blue (stop signal) after a stop signal delay (SSD), indicating to subjects not to respond (stop signal task testing action cancellation). A negative or neutral picture was presented as prime between fixation and task, in order to test emotional interference with performance in the two motor response inhibition tasks. Each trial lasted 1700 ms, followed by an ITI of 1700 ms. Only negatively primed trials are depicted as examples. The skull is used as a placeholder for negative images, but was not included in the stimulus data set of Task 6.

To additionally induce emotional interference with these two motor tasks, negative (Neg) and neutral (Neu) pictures were used as primes. We selected 216 photographs from three different picture sets [IAPS (Lang et al., 2008), NAPS (Marchewka, Żurawski, Jednoróg, & Grabowska, 2014) and EmoPicS (Wessa et al., 2010)], based on their normative arousal and valence ratings^4^. Negative images were negative (valence<3) and highly arousing (>5), neutral images were neutral (valence: 4 to 5.5) and not arousing (<4). Neutral and negative images were matched for depicted content. Emotional images were used twice during the course of the task, but in two separate runs and for different experimental conditions.

Each of the three runs of the task consisted of 144 trials. The first run started with an 8-seconds instruction. A fixation cross was presented at the start and the end of each run for 5 s. A black screen was presented during the ITI. Mean ITI was 1.7 s (range: 1 to 2.5 s). Each trial started with the presentation of a fixation cross for 200 ms, followed by the presentation of an either neutral or negative prime for 500 ms. The actual task (i.e., arrow) was presented immediately after the prime and lasted 1000 ms. The SSD was adapted using a staircase tracking procedure based on past performance to achieve a correct stopping rate of 50%. In case of correct stopping, the SSD was increased by 30 ms in the subsequent stop trial, while the SSD was reduced by 30 ms following incorrect stops. The SSD was adapted separately for Neg and Neu trials. The initial SSD was set to 200 ms; in the second and third runs, the final SSD of the preceding run was used as a starting value. The range of possible SSDs was between 30 and 540 ms.

Trials were presented in pseudo-randomized order. In the set-up of trial order we ensured that incongruent and stop trials were not followed by a trial of the same kind. Subjects were instructed to respond as quickly and accurately as possible.

Error rates and reaction times (RTs) served as behavioral outcome measures. Subjects with less than 44% correct stops were excluded from all analyses.

### Data acquisition

#### Electro-tactile stimulation

The Digitimer DS7A stimulator (Welwyn Garden City, UK) was used for painful electro-tactile stimulation in Tasks 3 and 4. Stimulation was given via a surface electrode with platinum pin (Specialty Developments, Bexley, UK) fixed to the right dorsal hand. An electro-tactile stimulus consisted of a train of three square-wave pulses of 2 ms duration each with an ITI of 50 ms. Current amplitude for stimulation was determined for each subject in standardized calibration procedures (see the respective task descriptions).

#### Physiological data

Electrocardiography (ECG) in Task 1 and skin conductance (SC) in Tasks 1, 3 and 4 were acquired with the Biopac MP150 recording system (BIOPAC Systems Inc., Goleta, California, USA) with a sampling rate of 1000 Hz, using the ECG 100C as well as the GSR 100C amplifiers in the behavioral laboratory and the EDA 100C amplifier in the MRI set-up. ECG data were registered with two pre-gelled ECG electrodes attached to the participant’s inner wrists and one ground electrode attached to the inner right ankle. Before attaching the electrodes, the skin surface was abraded. To avoid artefacts in the ECG data, subjects were instructed to sit very still during measurement. ECG data were band-pass filtered to frequencies between 0.5 and 35 Hz during data acquisition.

SC was measured using two self-adhesive Ag/AgACl electrodes prefilled with an isotonic electrolyte medium attached to the thenar and hypo-thenar of the non-dominant, left hand. The raw SC signal was amplified and low-pass filtered with a cut-off frequency of 1 Hz.

#### Behavioral data

Response data (hits, errors, omissions, RTs, in Tasks 2 and 6) and ratings of emotional state (stress, fear, valence, in Tasks 1, 3, 4 and 5) were collected via a computer keyboard in the behavioral laboratory (visit 1) and via an optical response keypad for the right hand (LUMItouch, Photon Control, Burnaby, Canada) during the MRI sessions (visits 2 and 3). Response data were logged by Presentation (NBS, San Francisco, USA). Rating scales were always presented visually on the computer screen. Answers were indicated by moving a blue box that was always initially displayed at a random position on the scale. For details on the specific questions and response poles, see the respective task descriptions.

#### Saliva samples

Saliva samples were collected at visit 1 with the “Salivette Cortisol, code blue” sampling device (Sarstedt, Nümbrecht, Germany). Saliva production was triggered by letting subjects chew on the synthetic swab of the salivette. There was only one saliva sample per sampling time point. Saliva samples were kept at room temperature until the end of the testing session and then stored at −18° C until analysis. Salivary cortisol and α-amylase were assayed in an external laboratory (Dresden LabService GmbH, Germany). Salivary cortisol is a biomarker for HPA activation (Kirschbaum & Hellhammer, 1989), since it correlates with cortisol levels in serum and plasma. Salivary α-amylase is a digestive enzyme that is used as an indirect marker of SNS activation (Bosch, Veerman, de Geus, & Proctor, 2011). Salivary cortisol was assayed with the cortisol saliva luminescence immunoassay (CLIA, IBL international, Hamburg, Germany). α-amylase was determined using an enzyme kinetic method previously described in Rohleder, Wolf, Maldonado, & Kirschbaum (2006), using the “Calibrator f.a.s.” solution (Roche Diagnostics, Mannheim, Germany) as standard and the α-amylase EPS Sys substrate reagent (Roche Diagnostics, Mannheim, Germany). Samples were assayed on four plates; inter-assay variation was below 3% for alpha-amylase and below 8% for cortisol.

### MRI data acquisition

Images were acquired at visits 2 and 3 on a Siemens 3 Tesla-Magnetom Tim Trio system (Siemens, Germany) running on software version Vb17, using a 32-channel head coil. Foam pads restricted head movement. Visual stimuli were presented on a screen at the head end of the scanner bore and projected to the subject’s visual field via a mirror that was fixed on the head coil. An infrared-light-camera-based eye tracking system (SMI, Teltow, Germany) was used to monitor subjects’ wakefulness. A multiband echoplanar imaging (EPI) sequence (TR=1000 ms, TE=29 ms, flip angle=56°, FOV=210 mm, voxel size=2.5×2.5×2.5 mm^3^, 60 slices, MB acceleration factor=4) was used for blood oxygen-level dependent (BOLD) fMRI, including resting-state scans, to achieve a fine-grained temporal resolution (Feinberg et al., 2010). fMRI was complemented by a T1-weighted MPRAGE-sequence (TR=1900 ms, TE=2.52 ms, flip angle=9°, FOV=250 mm, voxel size=1×1×1 mm^3^) as well as a T2–weighted (TR=6100ms, TE=79 ms, flip angle=120°, FOV=210mm, voxel size=0.6×0.5×3.0 mm^3^, 39 slices) and a diffusion-weighted (DTI) imaging (30 diffusion gradient directions, b1 = 0 s/mm^2^, b2=1300 s/mm^2^, TR=16900 ms, TE=117 ms, FOV=243 mm, voxel size=1.9×1.9×1.9mm^3^, 70 slices) sequence. Results for T2 and diffusion imaging are not reported here.

### Data analysis

Physiological, behavioral and rating data were read out and preprocessed in Matlab 2013b and 2015 (The Mathworks Inc., Natick, Massachusetts, USA). Statistical tests were performed in SPSS Statistics 23 for Windows (Armonk, NY: IBM Corp.).

#### Electrocardiography (ECG)

Electrocardiography (ECG) data recorded in Task 1 were analyzed with an costum-made Matlab script that discarded periods of missing data, if missing periods for a subject and phase did not exceed 30% of the time of an experimental phase. Otherwise, the phase was considered missing and the subject was excluded from further analysis. ECG data were first band-pass filtered to frequencies between 30 to 300 bpm. Subsequently, the Pan-Tompkins algorithm (Pan & Tompkins, 1983) was applied for QRS detection. Successful R-peak detection was scored by visual inspection. Heart rates (HR) were estimated on a beat-to-beat basis. HRs were averaged over the 5.15 to 5.75 minutes of each experimental phase (baseline, stress, recovery).

#### Skin conductance (SC)

Subjects were excluded from further analysis if recording during any experimental phase had failed. SCRs in Task 1 were detected and counted with an automated custom-made Matlab script. SCRs in Task 3 were manually scored with a half-automated custom-made Matlab script. SCRs were estimated as the largest trough-to-peak-amplitude (in μS) occurring after stimulus onset. Estimation of the trough-to-peak-amplitude was based on the approach by (Green, Kragel, Fecteau, & LaBar, 2014). Troughs were allowed from 0.9-4.0 s after cue onset, while peaks could occur until stimulus offset (6 s). SCRs larger than 0.02 μS were scored as responses, otherwise they were set to zero. Before statistical analysis, SCR amplitude data (Task 3) were log-transformed and range-corrected to assure normal distribution and to adjust for inter-individual differences in peak level.

SCL was calculated as the mean SC (in μS) over a 5.15 or 5.75-minutes experimental phase (Task 1) or an 8-seconds experimental condition (R or NR, Task 4). In Task 4, the SCL was baseline-corrected by subtracting the mean SCL 2 s prior to trial onset. SCL data were z-standardized within each subject for both Task 1 and Task 4 before statistical analysis (SCLz), to eliminate inter-individual variation in SCL.

#### Behavioral data

RTs in Tasks 2 and 6 were calculated for each experimental condition by averaging across trials. Stop signal reaction times (SSRT) in Task 6 were calculated by subtracting the mean stop signal delay (SSD) from the median RT in correct go trials. Hit rates and errors errors are given in percentage relative to the total number of trials per condition.

Ratings were generally averaged across experimental conditions, except for Tasks 1 and 3, where we were interested in experimentally induced changes over time. If the rate of missing values for the ratings was below 30%, missing values were replaced by the mean value of a given subject, else subjects were excluded from analysis. To deal with missing data for the baseline phase of Task 1, we averaged across the three baseline ratings.

### MRI data analysis

#### Image preprocessing

MRI data were analyzed in SPM8 (Wellcome Trust Centre for Neuroimaging, London, UK) implemented in the Matlab environment. Prior to data analysis, the first five EPI images of each run were discarded to establish equilibrium magnetization. In Task 3, run 1, the EPI images taken during the pre-experimental instruction and the habituation phase were discarded after preprocessing. Preprocessing included spatial realignment to the first image, co-registration of the mean functional image to the anatomical MPRAGE image, normalization to the MNI template via segmentation and spatial smoothing with a Gaussian filter with 8 mm FWHM. Estimated realignment parameters were used to screen for head movement exceeding translations of 3 mm or rotations of 2°. Subjects with excessive movement in an experimental task were discarded from the respective analysis. Additionally, the residual images resulting from the single-subject analysis were screened to control for artifacts.

#### Single-subject analysis (first level)

EPI images were temporally high-pass filtered with a cut-off of 128 s. A general linear model (GLM) was fitted for each subject and task to model the individual BOLD signal changes induced by experimental conditions. The three runs of Task 3 each corresponded to different experimental phases and were therefore analyzed in separate GLMs. The three runs of Task 6 were concatenated and analyzed within a single GLM. Regressors were boxcar functions corresponding in length to the experimental condition (block-type regressors), except for the painful electro-tactile stimulation (UCS) in Task 3 (CD, task run 1) and the trials in Task 6, for which we used stick functions (event-related regressors). To capture habituation of amygdala activity in threat-related tasks (Plichta et al., 2014), the categorical regressors in Tasks 3 and 4 were parametrically modulated with a decay function. For Task 3, we modelled a linear decay across trials of the same type, following previous work (e.g., Haaker et al., 2013); for Task 4, where there was no experiential learning of cue-pain contingencies (instructed fear), we modelled a steeper (quadratic) decay. The single-subject design matrices always included the regressors of interest plus some additional regressors of no interest (e.g., task instructions, strategy cues). Table 4 gives an overview of the regressors used in the tasks. All regressors were convolved with the canonical hemodynamic response function (HRF).

**Table 4.** Overview of regressors used in the single-subject fMRI analyses

| Task | Regressors of interest |  | Regressors of no interest |
| --- | --- | --- | --- |
|  | categorical | parametric modulators |  |
| 2 | Reward sensitivity<br>*Anticipation:*<br>+3 €, +0.5 €, ±0 €, -0.5 €, -3 €<br>(duration: 2-2.5 s)<br>*Outcome (feedback):*<br>Hits separately for +3 €, +0.5 €, ±0 €, -0.5 €, -3 € (2 s)<br>Misses separately for +3€, +0.5€, ±0€, -0.5€, -3€ (2 s) | — | Start instruction (7 s) |
| 3 | Safety learning and memory<br>*Conditioning (run 1):*<br>CS-, CS+e, CS+u (all trials) (6 s)<br>*Extinction (run 2):*<br>CS-, CS+e (6s) in bins of 2 trials<br>*Memory test (run 3):*<br>CS-, CS+e, CS+u (6s) in bins of 2 trials | Linear decay<br>—<br>— | UCS (0 s)<br>—<br>Start instruction (20 s) |
| 4 | Self-focused volitional reappraisal<br>No Threat/No Reappraisal (NT/NR),<br>No Threat/Reappraisal (NT/R),<br>Threat/No Reappraisal (T/NR),<br>Threat/Reappraisal (T/R) (8 s) | Quadratic decay | Start instruction (40 s)<br>Strategy cue (2 s)<br>Ratings (5 s) |
| 5 | Situation-focused volitional reappraisal<br>Positive/No Reappraisal (Pos/NR),<br>Positive/Reappraisal (Pos/R),<br>Neutral/No Reappraisal (Neu/NR),<br>Neutral/Reappraisal (Neu/R),<br>Negative/No Reappraisal (Neg/NR),<br>Negative/Reappraisal (Neg/R) (10 s) | — | Start instruction (40 s)<br>Strategy cue (2 s)<br>Ratings (5 s) |
| 6 | Emotional interference and motor response inhibition<br>Congruent go/Neutral (Con Go/Neu),<br>Congruent go/Negative (Con Go/Neg),<br>Incongruent go/Neutral (Incon Go/Neu),<br>Incongruent go/Negative (InconGo/Neg)<br>Stop/Neutral (Stop/Neu),<br>Stop/Negative (Stop/Neg) (0 s) |  | Start instruction (run 1) (8 s),<br>Incorrect trials |

#### Group analysis (second level)

Random-effects analyses were performed on the group level using single-subject beta or (in Task 6) contrast images of interest in SPM’s flexible factorial design. The contrast images of three runs of Task 6 were averaged before being entered into group analysis. Hypothesis-driven small volume analyses in predefined regions of interest (ROIs) used an initial uncorrected inclusion threshold of p_unc_<0.01. Within each ROI, correction for multiple comparisons was performed using the family-wise error (FWE) method at a threshold of p_SVC_<0.05 (SVC, small volume correction). Additional exploratory whole-brain analyses were conducted with a threshold of p_unc_<0.001. These are reported in the supplementary materials. Anatomical nomenclature follows the Harvard Oxford Brain Atlas (HOBA; Desikan et al., 2006; Frazier et al., 2005; Goldstein et al., 2007; Makris et al., 2006) and the Probablistic Cerebellum Atlas by Diedrichsen, Balsters, Flavell, Cussans, & Ramnani (2009), both thresholded at 25% with a 1-mm resolution. For the high-throughput anatomical labeling of whole brain activation peaks in the exploratory analyses, an automated costum-made Matlab script was applied that searched peak coordinates in the two atlases.

### Regions of interest (ROIs)

Table 5 gives an overview of our a priori ROIs and the employed literature sources. ROIs were based on peak MNI-coordinates retrieved from task-specific meta-analyses (Bartra, McGuire, & Kable, 2013; Buhle et al., 2013; Cieslik, Mueller, Eickhoff, Langner, & Eickhoff, 2015; Fullana et al., 2016; García-García et al., 2016; Lindquist, Satpute, Wager, Weber, & Barrett, 2016; Mechias, Etkin, & Kalisch, 2010). Where no suitable meta-analyses were available for a given task or effect within a task, informal reviews or suitable single studies were consulted. This was the case for vmPFC activity related to safety in Tasks 3 and 4 (Lonsdorf, Haaker, & Kalisch, 2014); for rostral dorsal anterior cingulate cortex/dorsomedial prefrontal cortex (rdACC/dmPFC) activity related to instructed fear in Task 4 (Kalisch & Gerlicher, 2014); and for subthalamic nucleus (STN) activity related to motor response inhibition in Task 6 (Aron & Poldrack, 2006). To assess amygdala activation in the fear-related Tasks 3 and 4, we used an anatomical mask from the HOBA, because recent meta-analyses did not report significant peak effects in this area (Fullana et al., 2018, 2016). We also used HOBA masks for the anterior and posterior hippocampus (aHPC, pHPC), to take into account the specific shape of this brain area. The a-p boundary was set at y=-25, approximately dividing the HPC mask in the middle of the y-plane. ROI masks were created with the SPM toolbox MARSBAR (Brett, Anton, Valabregue, & Poline, 2002). Subcortical ROIs were spheres of 6 mm diameter centered around the coordinates given in Table 5. Lateral cortical ROIs were spheres of 12 mm radius. Where above studies reported bilateral activations, we averaged the coordinates to achieve a symmetric distribution of the ROIs. Midline-cortical ROIs were midline-centered (x=0) boxes with dimensions 20 x 16 x 16 mm^3^ (as in Paret et al., 2011), and later work from our group). HOBA masks were thresholded at 50% with a 2-mm resolution.

**Table 5.** Overview of ROIs

| Task | Contrast | ROI | MNI |  |  | Reference |
| --- | --- | --- | --- | --- | --- | --- |
| 2 | Reward sensitivity |  |  |  |  |  |
|  | *Anticipation:* |  |  |  |  |  |
|  | Gain>Zero, | R + L NAcc | ±12 | 11 | -6 | Bartra et al. (2013) |
|  | Loss>Zero | R + L AI | ±31 | 21 | -7 | Bartra et al. (2013),<br>suppl. fig. 6A |
|  | Gain>Loss | R + L NAcc | ±12 | 11 | -6 | Bartra et al. (2013) |
|  |  | B vmPFC | 0 | 46 | -8 | Bartra et al. (2013) |
|  | *Outcome (feedback):* |  |  |  |  |  |
|  | Gain>No Gain | R + L NAcc | ±12 | 11 | -6 | Bartra et al. (2013) |
|  |  | B vmPFC | 0 | 46 | -8 | Bartra et al. (2013) |
|  | No Loss>Loss | R + L NAcc | ±12 | 11 | -6 | Bartra et al. (2013) |
|  | No Gain>Gain,<br>Loss>No Loss | R + L AI | ±36 | 20 | -3 | Bartra et al. (2013),<br>suppl. fig. 7 |
|  |  | B dACC | 0 | 22 | 44 |  |
| 3 | Safety learning and<br>memory |  |  |  |  |  |
|  | *Threat contrasts:* |  |  |  |  |  |
|  | CS+s>CS- (CD), | R + L AI | ±40 | 17 | 0 | Fullana et al. (2016) |
|  | CS+e>CS- (EX), | B dACC | 0 | 18 | 42 | Fullana et al. (2016) |
|  | CS+u>CS- (SR), | B pre-SMA | 0 | 16 | 54 | Fullana et al. (2016) |
|  | CS+u>CS+e (SR), | R + L thalamus | ±2 | -18 | 5 | Fullana et al. (2016) |
|  | CS+s>CS- (RN) | R + L BG | ±12 | 2 | -6 | Fullana et al. (2016) |
|  |  | R + L amygdala | – | – | – | HOBA |
|  | *Safety contrasts:* |  |  |  |  |  |
|  | CS->CS+s (CD), | B vmPFC | 0 | 40 | -12 | Lonsdorf et al. (2014) |
|  | CS+e>CS- (EX), | R + L lat. OFC | ±43 | 47 | -13 | Fullana et al. (2016) |
|  | CS+e>CS+u (SR), | R + L aHPC | – | – | – | HOBA |
|  | CS->CS+u (SR, RN) | R + L pHPC | – | – | – | HOBA |
| 4 | Self-focused<br>volitional reappraisal |  |  |  |  |  |
|  | T>NT | B rdACC/<br>dmPFC | 0 | 38 | 28 | Kalisch & Gerlicher<br>(2014)<sup>1</sup> |
|  |  | B dACC | 0 | 16 | 36 | Mechias et al. (2010) |
|  |  | R + L AI | ±34 | 20 | 3 | Mechias et al. (2010) |
|  |  | R + L BG,<br>thalamus<br>(see Task 3) | – | – | – | Fullana et al. (2016) |
|  |  | R + L amygdala | – | – | – | HOBA |
|  | NT>T | B vmPFC | 0 | 40 | -12 | Lonsdorf et al. (2014) |
|  |  | R + L lat. OFC | ±43 | 47 | -13 | Fullana et al. (2016) |
|  |  | R + L aHPC | – | – | – | HOBA |
|  |  | R + L pHPC | – | – | – | HOBA |
|  | R>NR | B dmPFC | 0 | 21 | 54 | Buhle et al. (2013) |
|  |  | B SMA | 0 | 6 | 63 | Buhle et al. (2013) |
|  |  | B ACC | 0 | 24 | 30 | Buhle et al. (2013) |
|  |  | R + L dlPFC | ±42 | 9 | 51 | Buhle et al. (2013) |
|  |  | L AI | -36 | 21 | -3 | Buhle et al. (2013) |
|  |  | R + L IFG | ±56 | 23 | 6 | Buhle et al. (2013) |
|  |  | L MTG | -51 | -39 | 3 | Buhle et al. (2013) |
|  |  | L SPL | -42 | -66 | 42 | Buhle et al. (2013) |
| 5 | Situation-focused volitional reappraisal |  |  |  |  |  |
|  | Pos/NR>Neu/NR | B vmPFC | 0 | 39 | -9 | Lindquist et al. (2016) |
|  |  | R + L NAcc | ±12 | 11 | -6 | Bartra et al. (2013) |
|  | Neg/NR>Neu/NR | R + L amygdala | ±21 | -4 | -15 | García-García et al. (2016) |
|  |  | R IFG | 50 | 16 | 22 | García-García et al. |
|  |  | R + L ILOC | ±50 | -68 | 6 | García-García et al. |
|  |  | R + L Brainstem<sup>2</sup> | ±4 | -28 | -6 | García-García et al. |
|  | R>NR | see Task 4 |  |  |  | Buhle et al. (2013) |
|  |  | R + L NAcc | ±12 | 11 | -6 | Bartra et al. (2013) |
| 6 | Emotional interference and motor response |  |  |  |  |  |
|  | *Inhibition common:* |  |  |  |  |  |
|  | Incon Go>Con Go, Stop>Con Go | B pre-SMA | 0 | 16 | 48 | Cieslik et al. (2015) |
|  |  | R + L AI | ±35 | 21 | -4 | Cieslik et al. (2015) |
|  |  | R IFG (pars opercularis) | 48 | 12 | 30 | Cieslik et al. (2015) |
|  | *Inhibition separate:* |  |  |  |  |  |
|  | Incon Go>Con Go | R + L dlPFC | ±28 | 3 | 53 | Cieslik et al. (2015) |
|  |  | R + L SPL | ±26 | -60 | 54 | Cieslik et al. (2015) |
|  | Stop>Con Go | R ACC | 0 | 30 | 32 | Cieslik et al. (2015) |
|  |  | R IFG (pars triangularis) | 56 | 20 | 4 | Cieslik et al. (2015) |
|  |  | R + L STN | ±10 | -15 | -5 | Aron & Poldrack |
|  | Neg>Neu | see Task 5 | – | – | – | García-García et al. |
|  | Interaction 1<sup>3</sup> | all inhibition |  |  |  | Cieslik et al. (2015) |
|  | Interaction 2<sup>4</sup> | R + L amygdala | ±21 | -4 | -15 | García-García et al. (2016) |

MNI, Montreal Neurological Institute; L, left; R, right; B, midline-centered bilateral box for midline-cortical ROIs; HOBA, Harvard-Oxford Brain Atlas. For abbreviations of experimental conditions, see task descriptions. Anatomical terms: ACC, anterior cingulate cortex; AI, anterior insula; BG, basal ganglia; dACC, dorsal ACC; dlPFC, dorsolateral PFC; dmPFC, dorsomedial PFC; HPC, hippocampus; IFG, inferior frontal gyrus; ILOC, inferior lateral occipital cortex; MTG, middle temporal gyrus; NAcc, Nucleus accumbens; OFC, orbitofrontal cortex; pre-SMA, pre-supplementary motor area; PFC, prefrontal cortex; rdACC, rostradorsal ACC; SMA, supplementary motor area; SPL, superior parietal lobule; STN, subthalamic nucleus; vmPFC, ventromedial PFC. <sup>1</sup>average of listed peaks. <sup>2</sup>García-García et al. (2016) reported a bilateral brainstem cluster with a right-sided (R) peak. We additionally included an equivalent left-sided (L) ROI. <sup>3</sup>((Incon Go, Stop)/2>Con Go)<sub>Neg</sub> > ((Incon Go, Stop)/2>Con Go)<sub>Neu</sub>. <sup>4</sup>(Con Go>(Incon Go, Stop)/2)<sub>Neg</sub> > (Con>(Incon Go, Stop)/2)<sub>Neu</sub>.

## Results

Table 6 gives an overview of the sample sizes available for the analysis of the different outcomes collected in the different tasks. Reasons for exclusion were errors in the experimental paradigm, missing data, failures in technical equipment, outliers, invalid subject performance and excessive head movement during MRI measurement. In general, exclusion rates were low (<10%), with the exception of the heart rate data in Task 1, which were affected by technical equipment failures. From the analysis of salivary α-amylase in Task 1 we excluded one outlier with a concentration of more than 3 standard deviations (SD) above the group mean. In Task 6, we had to exclude seven subjects because they did not achieve the predefined criterion of a stopping rate of at least 44%.

**Table 6.** Sample sizes for the different outcomes

| Task | Outcome | N | % valid |
| --- | --- | --- | --- |
| 1 | Stress rating | 51<sup>1,2</sup> | 94 |
|  | HR | 41<sup>1,2,3</sup> | 76 |
|  | nSCR, SCLz | 49<sup>1,2</sup> | 90 |
|  | Salivary cortisol | 52<sup>1</sup> | 96 |
| | Salivary α-amylase | 51<sup>1,4</sup> | 94 |
| 2 | RT | 53<sup>5</sup> | 98 |
|  | fMRI | 52<sup>5,6</sup> | 96 |
| 3 | Fear rating | 49<sup>1,2</sup> | 90 |
|  | SCR | 51<sup>1</sup> | 94 |
|  | fMRI | 50<sup>6</sup> | 93 |
| 4 | Fear rating | 54 | 100 |
|  | SCLz | 54 | 100 |
|  | fMRI | 54 | 100 |
| 5 | Emotional state rating | 53<sup>1</sup> | 98 |
|  | fMRI | 52<sup>1,6</sup> | 96 |
| 6 | RT, SSRT, error rates | 47<sup>5,3</sup> | 87 |
|  | fMRI | 47<sup>5,6</sup> | 87 |

<sup>1</sup>Error in experimental paradigm, <sup>2</sup>missing data, <sup>3</sup>technical equipment failure, <sup>4</sup>discarded outliers ( α-amylase > mean + 3SD), <sup>5</sup>invalid subject performance, <sup>6</sup>excessive head movement. HR, heart rate; nSCR, number of skin conductance responses; RT, reaction time; SCLz, skin conductance level, z-standardized; SCR, skin conductance response; SSRT, stop signal reaction time.

### Task 1: Stress reactivity and recovery

In the stress experience questionnaire, 55.6% of subjects listed the mental arithmetic task and the explosion sounds among the three most stressful components of the test; further listed components were negative performance feedback (44.4%), loud noise (35.2%), negative pictures (29.6%), speed of the task (27.8%), and threat of monetary loss (14.8%). Figure 8 shows average time courses of stress responses in the different outcome measures. The rate of stress responders, i.e., subjects that showed a numerical increase in a given outcome measure from baseline to the stress-induced group peak shown in Figure 8, varied between 65.4% for cortisol and 100% for stress ratings and skin conductance (see Table 7). Significant stress reactivity (baseline vs. peak) was observed in all measures; full stress recovery, defined as a non-significant difference between baseline and the recovery phase (or the last measurement for cortisol and α-amylase), was observed in all measures but skin conductance level (SCL) (see Table 7; comp. Figure 8 C).

**Figure 8.**
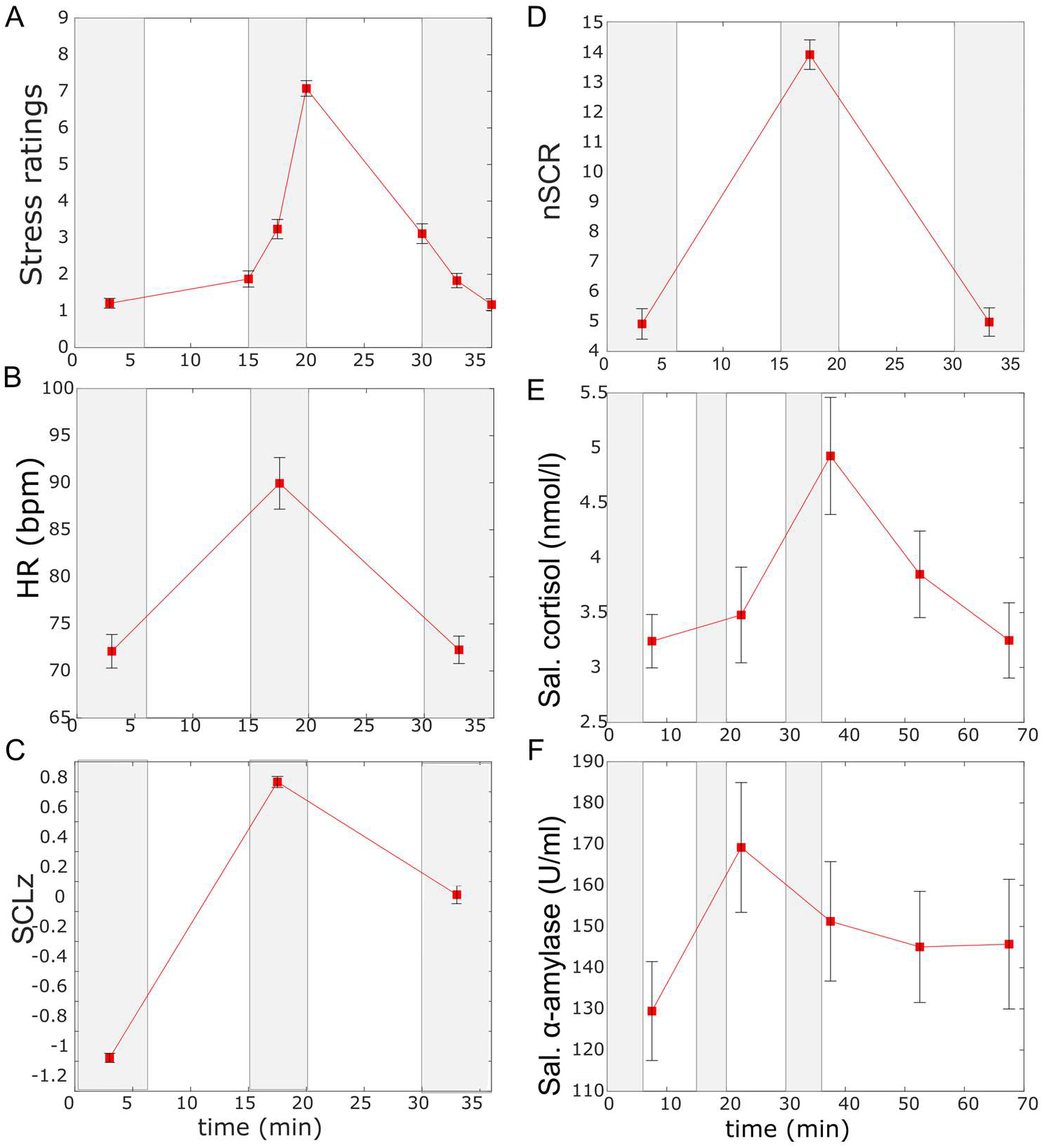
Results of Task 1 (stress reactivity and recovery): Response time courses. Grey bars indicate experimental phases: rest (baseline), stress, rest (recovery). (For experimental time line, see Figure 2) Plotted are group means±standard error of the mean (SEM). For Ns, see Table 7. HR, heart rate; SCLz, SCL z-standardized within each subject; nSCR, number of SCRs; Sal., salivary.

**Table 7.** Results of Task 1 (stress reactivity and recovery): Statistics

| Outcome | N | Response<br>rate | Baseline vs.<br>peak |  | Baseline vs.<br>recovery |  | Time effect |  |  |  |
| --- | --- | --- | --- | --- | --- | --- | --- | --- | --- | --- |
|  |  | in % | *t-score* | *p* | *t-score* | *p* | *F-* | *df<sub>time</sub>* | *df<sub>err</sub>* | *p* |
| Stress | 5 | 100 | -25.5 | <0.001 | 0.2 | 0.85 | 137.38 | 4.16 | 208 | <0.001 |
| HR | 4 | 97.6 | -7.23 | <0.001 | -0.24 | 0.81 | 49.47 | 1.1 | 44. | <0.001 |
| SCLz | 4 | 100 | -57.95 | <0.001 | -12.4 | <0.001 | 291.49 | 1.18 | 56. | <0.001 |
| nSCR | 4 | 100 | -18.97 | <0.001 | 0.21 | 0.83 | 313.33 | 1.7 | 81. | <0.001 |
| Sal. | 5 | 65.4 | -4.04 | <0.001 | -0.03 | 0.98 | 10.43 | 2.29 | 117 | <0.001 |
| Sal. α-<br>amylase | 5<br>1 | 78.4 | -4.27 | <0.001 | -1.49 | 0.14 | 4.26 | 4 | 200 | <0.001 |
|  |  |  |  | * |  |  |  |  |  | * |

\* significant at $p < 0.05$ . HR, heart rate; nSCR, number of skin conductance responses; SCLz, skin conductance level, z-standardized; SCR, skin conductance response.

### Task 2: Reward sensitivity

#### Behavioral data

The targeted hit rate (66%) was well approximated, with subjects hitting on average on 63.7% of trials (SD=6%, range: 44.4 to 75.5%; Table 8). Incentive magnitude had a significant effect on RTs (F_(3.4;176.81)_=3.31, p=0.01; N=53). This effect was driven by reduced RTs for Gain (+3 €, +0.5 €) compared to Zero (±0 €) trials (t_+3>0 €_=-2.77, p=0.007; t_+0.5>0 €_=-2.69, p=0.01). Loss trials (−3 €, −0.5€) did not differ significantly from Gain or Zero trials.

**Table 8.** Results of Task 2 (reward sensitivity): Behavior

| Incentive<br>magnitude | Hit rates (%) |  | RT (ms) |  |
| --- | --- | --- | --- | --- |
|  | *M* | *SD* | *M* | *SD* |
| +3 € | 66.46 | 8.85 | 297.74 | 56.99 |
| +0.5 € | 64.78 | 10.39 | 296.00 | 61.20 |
| ±0 € | 61.22 | 12.64 | 313.82 | 55.48 |
| -0.5 € | 62.90 | 10.44 | 304.45 | 54.03 |
| -3 € | 63.32 | 11.02 | 307.01 | 50.31 |

#### fMRI data

##### Anticipation

ROI analyses yielded the expected strong bilateral nucleus accumbens (NAcc) and anterior insula (AI) activations for both the contrasts Gain>Zero and Loss>Zero (Table 9, Figure 9 A,B), with the corresponding U-shaped response profile across the five incentive conditions (Figure 9 C). Gain>Loss induced stronger activation in bilateral NAcc and left vmPFC (Table 9). Here and in further sections, the results tables list only those ROIs containing voxels that survived the initial inclusion threshold for small volume correction (SVC) of p_unc_<0.01 (see the description of MRI data analysis). For those ROIs, all peaks are listed. Peaks surviving SVC at p_SVC_<0.05 are marked with * in the tables. The activation maps in the results figures are thresholded at p_unc_<0.01, to provide unbiased information about the spatial distribution of the effects. Exploratory whole-brain analyses for the anticipation phase at a threshold of p_unc_<0.001 are reported in the supplementary materials.

**Table 9.**
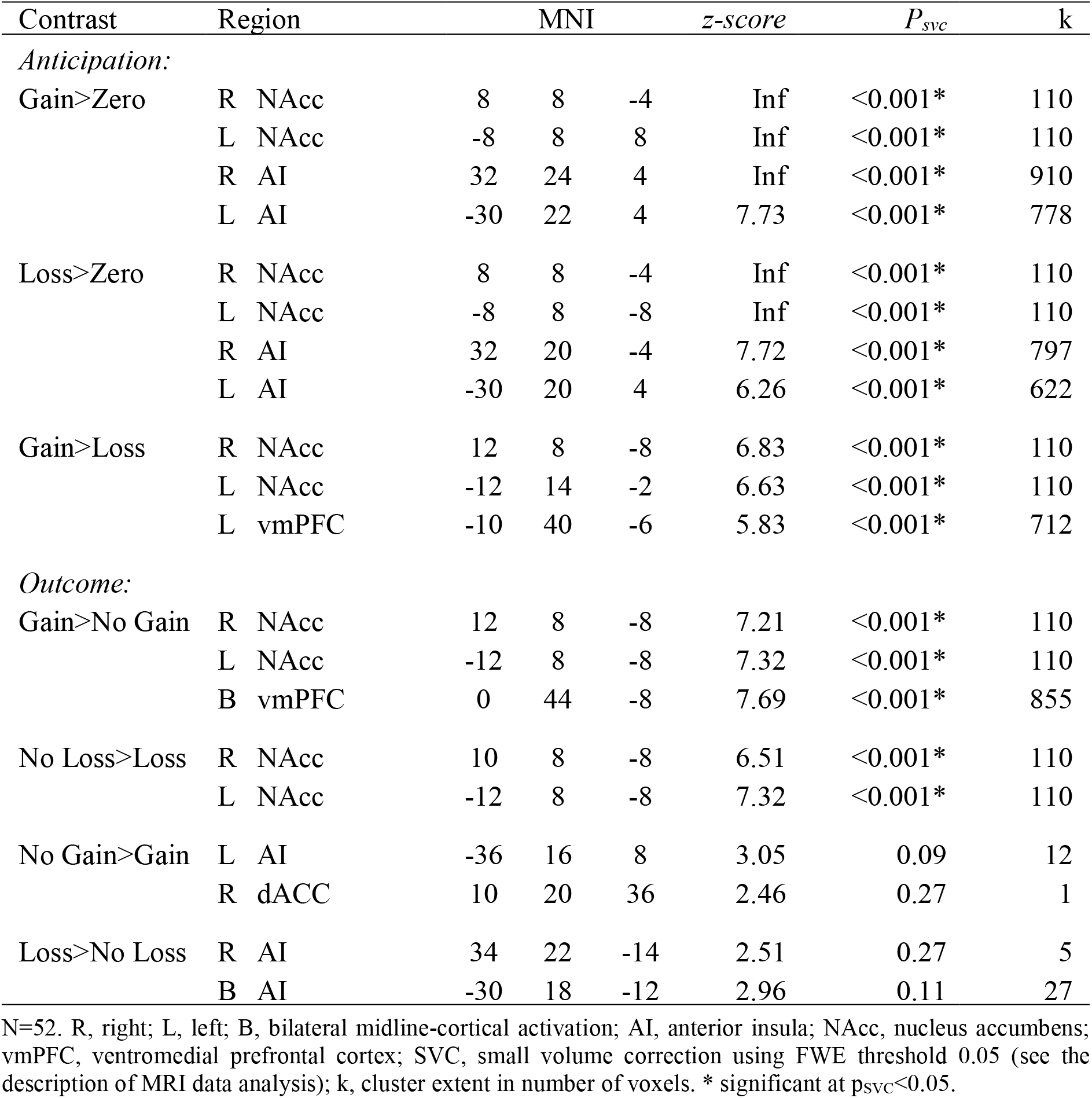
Results of Task 2 (reward sensitivity): fMRI ROI analyses.

**Figure 9.**
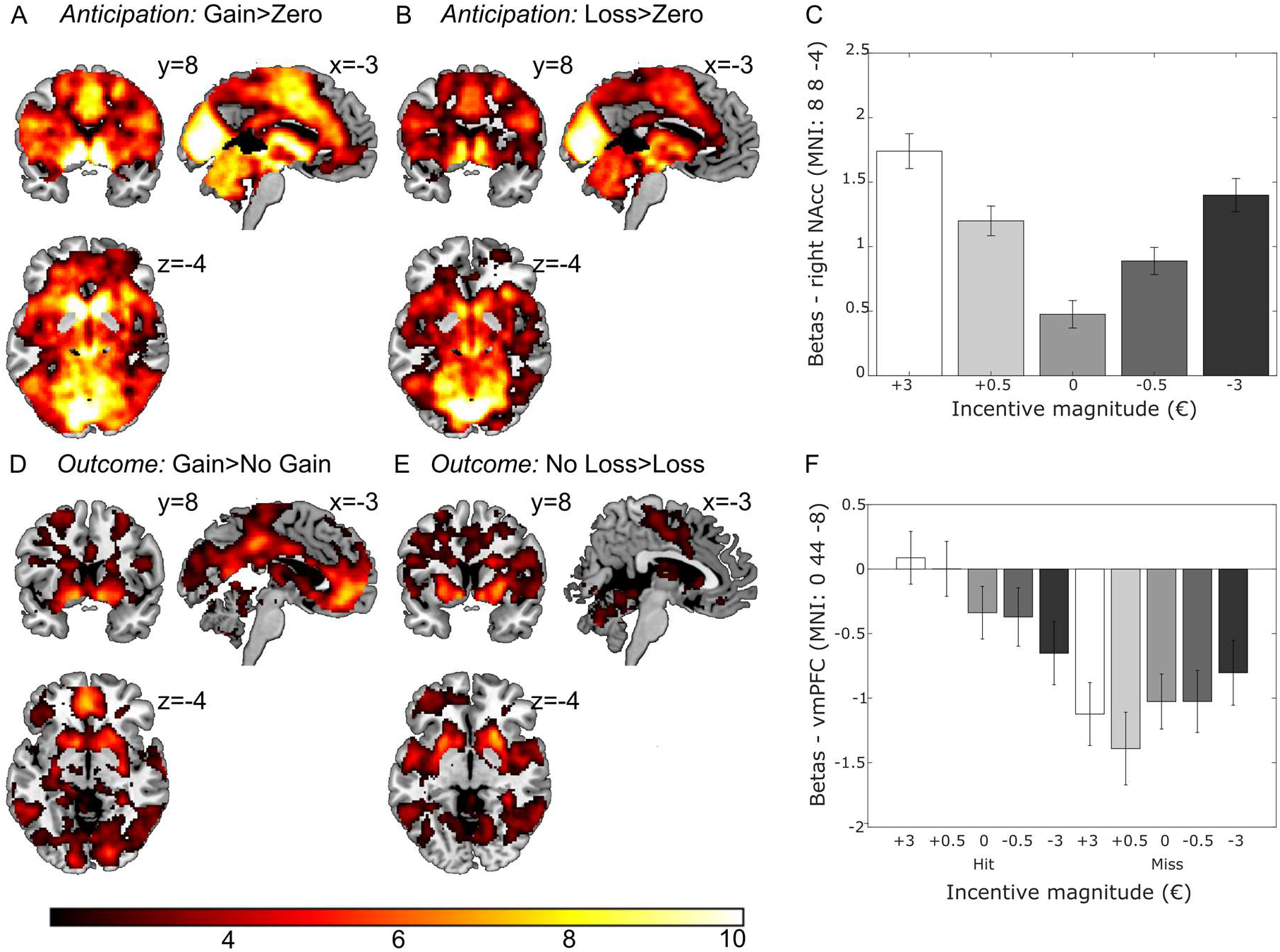
Results of Task 2 (reward sensitivity): fMRI activations. N=52. Display threshold: p_unc_<0.01. Color bar shows t-values. Bar graphs show means±SEM.

#### Outcome (feedback)

Realizing relative to not realizing a gain (hit vs. missed +3 € and +0.5 € trials; contrast: Gain>No Gain) as well as avoiding relative to not avoiding a loss (hit vs. missed −3 € and −0.5 € trials; contrast: No Loss>Loss) both activated the bilateral NAcc ROI (Table 9, Figure 9 D, E). Gain>No Gain additionally recruited the bilateral ventromedial prefrontal cortex (vmPFC). This activation was characterized by strong relative deactivations when missing a gain (No Gain; Figure 9 F). Notably, there was no significant ROI activation for missed gains (No Gain>Gain) and for losses (Loss>No Loss). Hence, to summarize, subjects reacted strongest to realized gains (see also Figure 9 F). Additional exploratory whole-brain results are reported in suppl. materials.

### Task 3: Safety learning and memory

#### Contingency ratings

Contingency awareness was assessed after each experimental phase. All subjects reported that they had received pain stimulation (UCS) during conditioning (CD, task run 1) and no stimulation during extinction (EX, task run 2) and the memory retrieval tests (SN and RN, task run 3 the next day). The rate of subjects who reported to know when they would receive a UCS during conditioning was 86.3%. After conditioning, 98% (CS+e) and 100% (CS+u) of subjects, respectively, could tell that the respective CS+ had been coupled with a UCS, while 94.1% of subjects had understood that the CS− had never been followed by a UCS. After extinction and the memory tests, the rate of subjects indicating that they had not received UCSs with the presented CSs was high, ranging from 92.1 to 100%. 94.1% correctly identified the background picture (context) used during conditioning, and all subjects identified the picture used during extinction. Hence, overall, contingency awareness was highly developed.

#### Fear ratings and SCRs

Fear and SCRs were assessed at every trial. Figure 10 shows responses per phase. In conditioning, subjects successfully learned to discriminate between the two CS+s and the CS− in both measures (see also Table 10), indicating the CS+s were identified as threatening and the CS− as safe. CS+e/CS− discrimination was still present in early extinction (E-EX) in ratings but not SCRs, an effect that can be explained by generalized heightened arousal to any stimuli presented at the beginning of a new task run, after an experimental break. Accordingly, a reduction in CS+e/CS− discrimination during extinction, indicating safety learning to the CS+e, could only be observed in ratings (stimulus by time: F_(1;48)_=14.77, p<0.001; stimulus: F_(1;48)_=21.89, p<0.001; time: F_(1;48)_=101.06, p<0.001; see also Table 10; cf. SCRs: stimulus by time: F_(1;50)_=0.005, p=0.944; stimulus: F_(1;50)_=0.52, p=0.474; time; F_(1;50)_=42.79, p<0.001).

**Figure 10.**
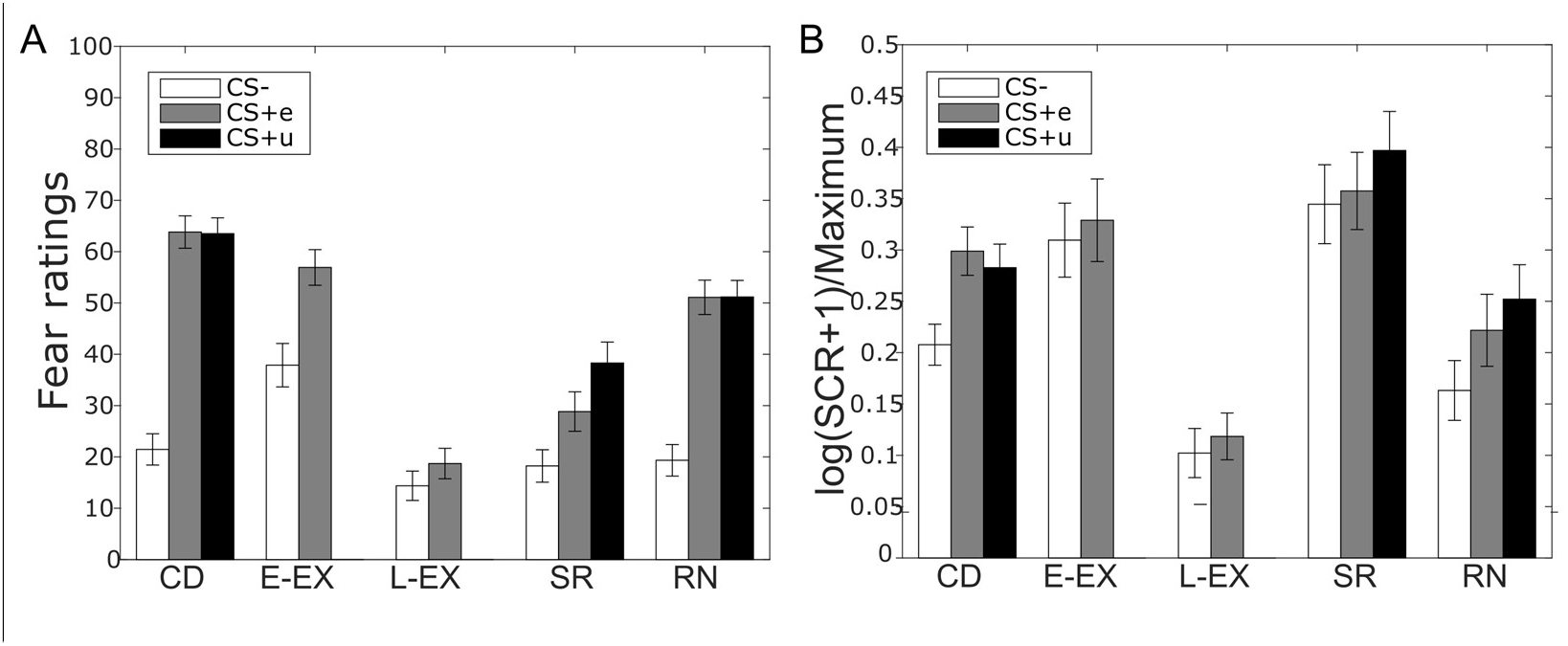
Results of Task 3 (safety learning and memory): Fear ratings and SCRs. Plotted are group means±SEM. N=49 for fear ratings; N=51 for SCR. Conditioning: average of all trials; all other bars: average of first two trials (E-EX, SR, RN) or last two trials (L-EX). CD, conditioning; E-EX, early extinction; L-EX, late extinction; RN, renewal; SR, spontaneous recovery.

**Table 10.**
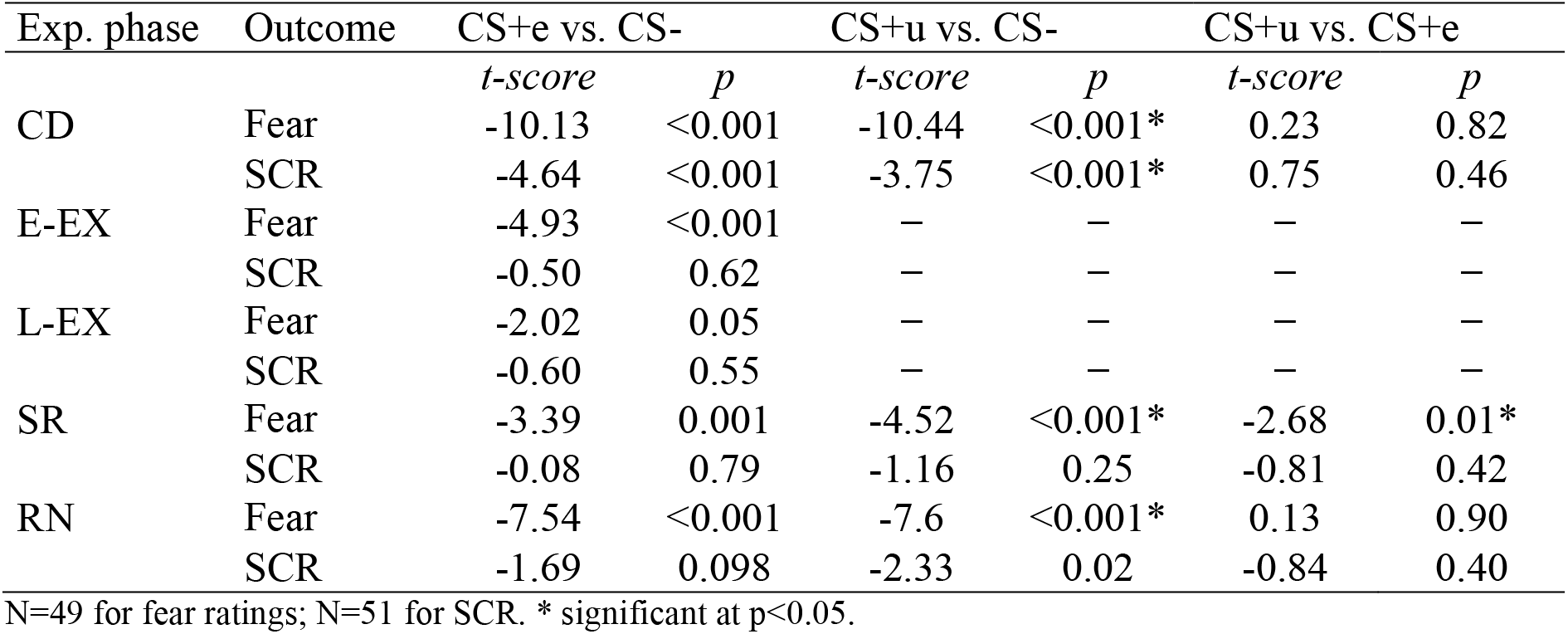
Results of Task 3 (safety learning and memory): Statistics

Discrimination in the fear ratings was re-established during spontaneous recovery for both the CS+e and the CS+u, i.e., fear also returned to the extinguished CS+ (CS+e). However, responding was significantly more pronounced to the unextinguished CS+u than to the extinguished CS+e (Table 10), reflecting retrieval of the safety memory associated with the CS+e during extinction (“extinction memory”). No discrimination was observed in SCRs, perhaps again reflecting generalized arousal. Renewal was generalized, leading to comparable CS+/CS− discrimination for both CS+s, and also extended to SCRs (Table 10). Hence, no remaining safety memory expression was observable during renewal.

#### fMRI data

##### Conditioning

ROI analyses yielded the expected strong activation in the canonical fear network in the threat-related categorical contrast CS+s>CS− (Table 11, Figure 11 A); the parametrically modulated CS+s>CS− contrast detected linearly decaying activation in an amygdalar-hippocampal transition zone (Figure 11 B). Activation in the bilateral vmPFC and the orbitofrontal cortex (OFC) was found in the safety-related contrast CS−>CS+s (Figure 11 A). Additional exploratory whole-brain results are reported in the suppl. materials.

**Table 11.**
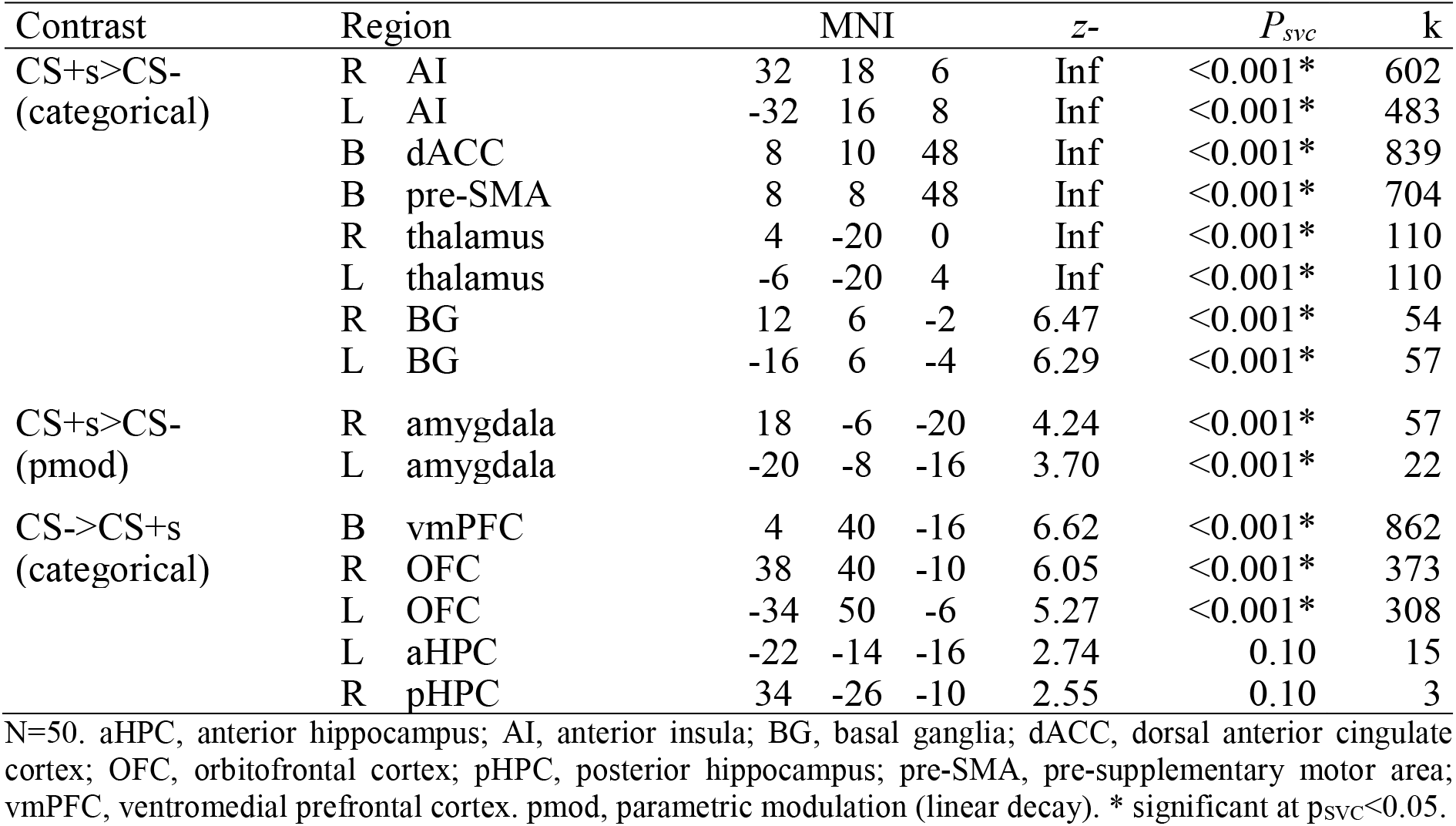
Results of Task 3 (safety learning and memory): fMRI ROI analyses for conditioning.

**Figure 11.**
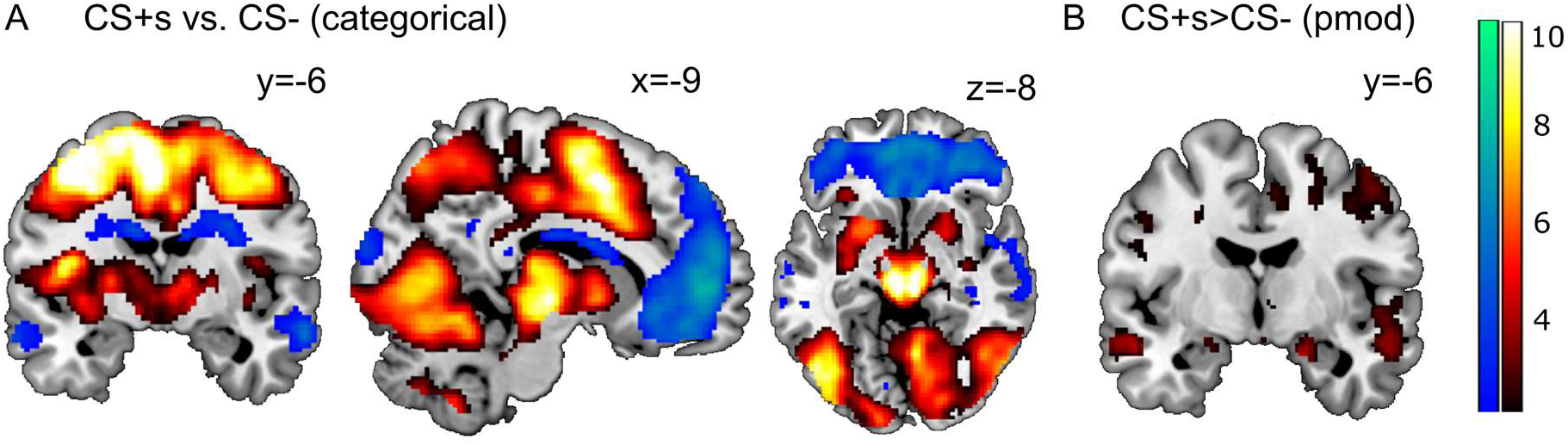
Results of Task 3 (safety learning and memory): fMRI activations for conditioning. N=50. pmod, parametric modulation (linear decay). Display threshold: p_unc_<0.01. Hot color bar (right) shows t-values for activations; cold color bar (left) shows t-values for deactivations.

##### Extinction

As in SCRs, there was no detectable CS+e/CS− discrimination in BOLD responses. However, also mirroring the SCR results, activation globally decreased from early to late extinction (E-EX>L-EX) in the major fear network ROIs (Table 12, Figure 12 A). Global activation increases in the posterior hippocampus (pHPC) ROI (Table 12) could be attributed as per whole-brain analysis to an adjacent bilateral lingual gyrus activation (Figure 12 B). For additional whole-brain results, see suppl. materials.

**Table 12.** Results of Task 3 (safety learning and memory): fMRI ROI analyses for extinction, spontaneous recovery, renewal.

| Contrast | Region |  | MNI |  | *z-score* | *P<sub>svc</sub>* | k |  |
| --- | --- | --- | --- | --- | --- | --- | --- | --- |
| *Extinction* |  |  |  |  |  |  |  |  |
| CS+e>CS- | - |  |  |  |  |  |  |  |
| E-EX>L-EX<sup>1</sup> | R | AI | 34 | 18 | -10 | 5.25 | <0.001* | 408 |
|  | L | AI | -30 | 18 | -6 | 3.73 | 0.01* | 179 |
|  | R | pre-SMA/dACC | 4 | 20 | 50 | 3.48 | 0.02* | 170 |
|  | R | amygdala | 28 | 2 | -20 | 3.29 | 0.02* | 20 |
| L-EX>E-EX<sup>1</sup> | R | aHPC | 36 | -30 | -10 | 2.37 | 0.21 | 1 |
|  | L | aHPC | -22 | -14 | -16 | 2.90 | 0.06 | 21 |
|  | R | pHPC | 30 | -38 | -4 | 4.14 | <0.001* | 27 |
|  | L | pHPC | -30 | -36 | -8 | 3.27 | 0.02* | 43 |
| *Spontaneous recovery:* |  |  |  |  |  |  |  |  |
| CS+u>CS- | R | AI | 36 | 16 | -8 | 4.75 | <0.001* | 682 |
|  | L | AI | -34 | 16 | 10 | 4.05 | <0.001* | 271 |
|  | B | pre-SMA/dACC | 0 | 14 | 50 | 3.30 | 0.05* | 175 |
|  | R | BG | 10 | -2 | -2 | 3.59 | <0.001* | 15 |
|  | R | amygdala | 18 | -4 | -12 | 3.66 | 0.01* | 16 |
|  | L | amygdala | -22 | -4 | -16 | 4.05 | <0.001* | 78 |
| CS+u>CS+e | R | AI | 36 | 16 | -8 | 3.82 | 0.01* | 418 |
|  | L | AI | -34 | 20 | -6 | 3.17 | 0.07 | 52 |
|  | L | thalamus | -6 | -14 | 4 | 2.88 | 0.03* | 9 |
|  | R | BG | 10 | -2 | -4 | 3.22 | 0.01* | 12 |
|  | L | amygdala | -24 | -10 | -14 | 3.33 | 0.02* | 45 |
| CS+e>CS+u | - |  |  |  |  |  |  |  |
| CS->CS+u | L | vmPFC | -8 | 44 | -14 | 3.55 | 0.02* | 57 |
|  | R | aHPC | 28 | -14 | -18 | 2.95 | 0.07 | 12 |
|  | L | aHPC | -30 | -16 | -20 | 2.65 | 0.13 | 1 |
|  | R | pHPC | 30 | -38 | -2 | 2.73 | 0.07 | 2 |
| *Renewal:* |  |  |  |  |  |  |  |  |
| CS+s>CS- | R | AI | 36 | 26 | -6 | 2.34 | 0.39 | 1 |
|  | L | AI | -36 | 18 | 10 | 3.27 | 0.05* | 96 |
| CS+u>CS+e | - |  |  |  |  |  |  |  |
| CS+e>CS+u | - |  |  |  |  |  |  |  |
| CS->CS+s | B | vmPFC | 0 | 36 | -20 | 3.42 | 0.03* | 177 |
|  | R | OFC | 36 | 38 | -10 | 3.81 | 0.01* | 67 |
|  | L | OFC | -40 | 36 | -14 | 2.92 | 0.14 | 9 |
|  | R | aHPC | 24 | -12 | -20 | 3.22 | 0.04* | 101 |
|  | L | aHPC | -22 | -14 | -20 | 2.86 | 0.08 | 18 |
|  | R | pHPC | 30 | -32 | -10 | 2.98 | 0.04* | 12 |

N=50. aHPC, anterior hippocampus; AI, anterior insula; BG, basal ganglia; dACC, dorsal anterior cingulate cortex; OFC, orbitofrontal cortex; pHPC, posterior hippocampus; pre-SMA, pre-supplementary motor area; vmPFC, ventromedial prefrontal cortex. E-EX, early extinction (first two trials); L-EX, late extinction (last two

**Figure 12.**
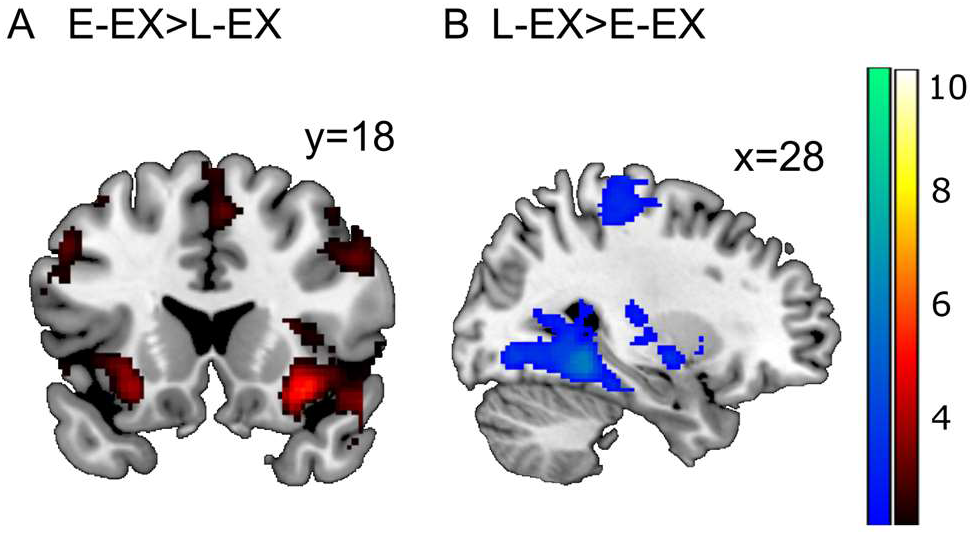
Results of Task 3 (safety learning and memory): fMRI activations for extinction. N=50. Display threshold: p_unc_<0.01. Hot color bar (right) shows t-values for activations; cold color bar (left) shows t-values for deactivations.

##### Spontaneous recovery

ROI analyses showed significant fear network activation in the contrast CS+u>CS−. Additionnally, fear network activation was significantly increased for the CS+u compared to the CS+e. (CS+u>CS+e) (Table 12, Figure 13 A-C). While this supports stronger fear memory retrieval to the CS+u than to the CS+e (see also rating results), we did not observe corresponding correlates of safety (extinction) memory retrieval to the CS+e (inverse contrast CS+e>CS+u) at the group level. However, the CS− relative to the CS+u activated the vmPFC ROI (Table 12, Figure 13 D,E). For whole-brain results, see suppl. materials.

**Figure 13.**
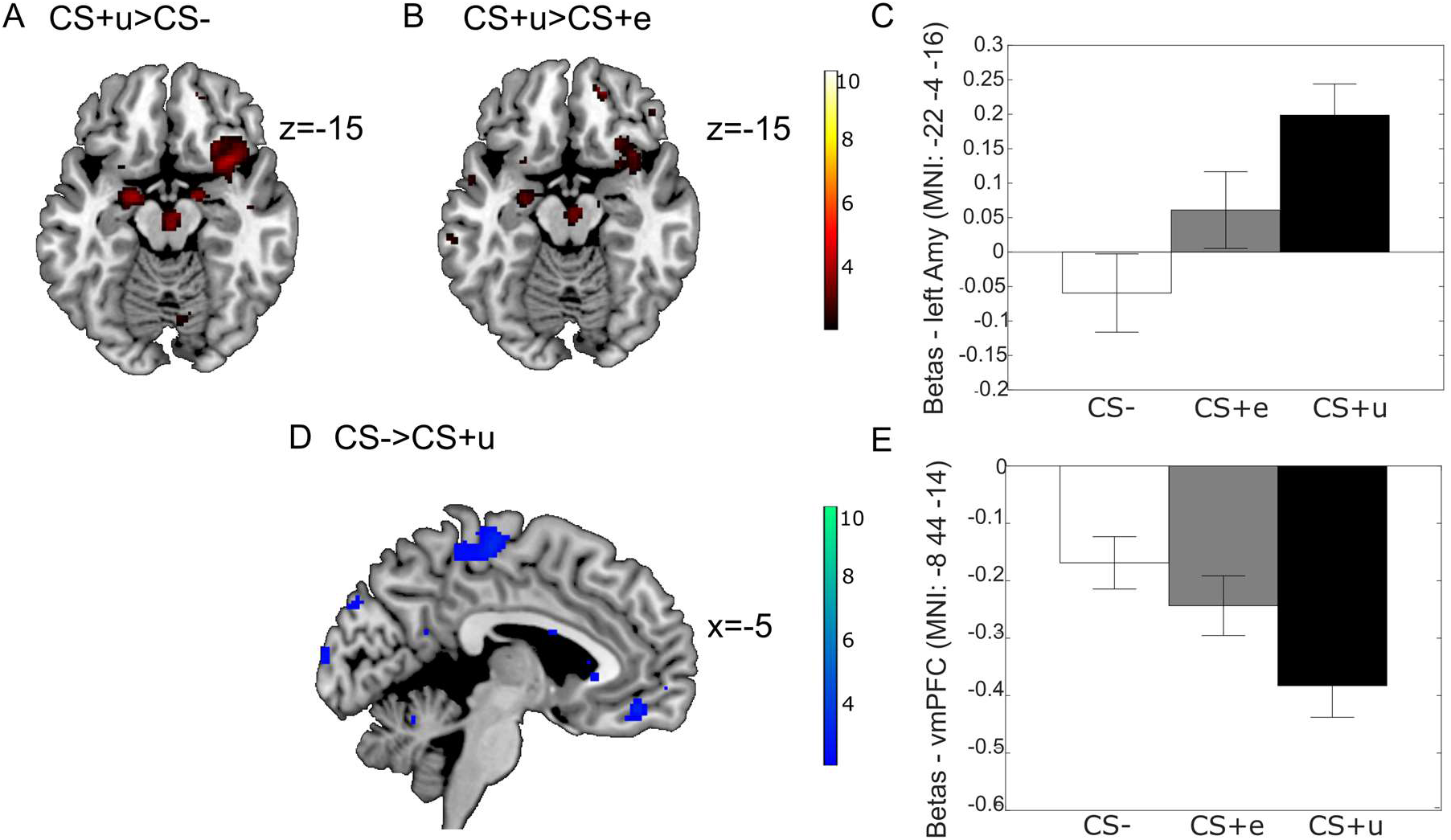
Results for Task 3 (safety learning and memory): fMRI activations for spontaneous recovery. N=50. Display threshold: p_unc_<0.01. Hot color bar (right) shows t-values for activations; cold color bar (left) shows t-values for deactivations. Bar graphs show means±SEM.

##### Renewal

In line with the behavioral data showing generalized renewal (4.3.2), the contrast CS+s>CS− led to increased activation in parts of the fear network (left AI; Table 12, Figure 14 A), with both CS+s showing similar activation (Figure 14 B). The CS− compared to the CS+s recruited the vmPFC, right OFC, right anterior and posterior HPC and left aHPC. See suppl. materials for whole-brain results.

**Figure 14.**
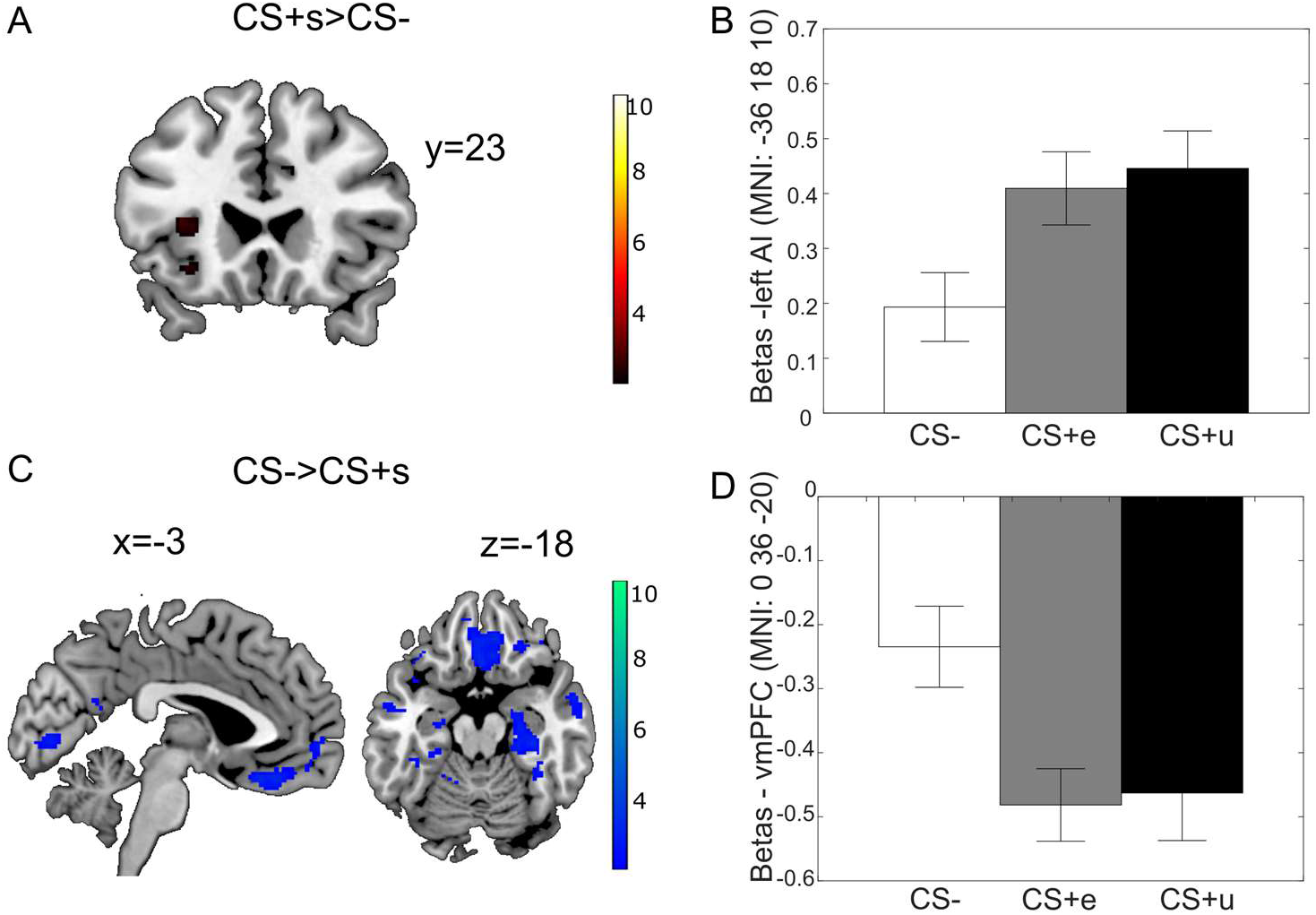
Results for Task 3 (safety learning and memory): fMRI activations for renewal. N=50. Display threshold: p_unc_<0.01. Hot color bar (A) shows t-values for activations; cold color bar (C) shows t-values for deactivations. Bar graphs show means±SEM.

### Task 4: Self-focused volitional reappraisal

#### Ratings and SCL

After the experiment, subjects reported to have successfully applied the reappraisal strategy (M=70 out of 100, SD=16.3, range: 28 to 100, N=52). Trial-by-trial fear ratings showed a significant main effect of threat (T>NT: F_(1;53)_=281.97, p<0.001), indicating successful fear induction by the threat instruction. The impact of reappraisal consisted in a significant main effect of reappraisal (R>NR: F_(1;53)_=37.59, p<0.001) and a significant threat by reappraisal interaction (F_(1;53)_=29.31, p<0.001). The interaction was characterized by an anxiolytic effect of reappraisal that was specific to the threat condition ((T/NR>T/R)>(NT/NR>NT/R); post-hoc test for T/NR>T/R: t=-6.30, p<0.001; *Figure 15* A). Successful threat induction was also indicated by a significant main effect of threat on SCLz (F_(1;53)_=35.18, p<0.001). However, reappraisal did not reduce sympathetic arousal (no R>NR main effect: F_(1;53)_=2.79, p=0.10), but rather tended to increase it (*Figure 15* B), suggesting application of the strategy might involve mental effort. Also, there was no significant threat by reappraisal interaction on SCLz (F_(1;53)_=1.06, p=0.308).

**Figure 15.**
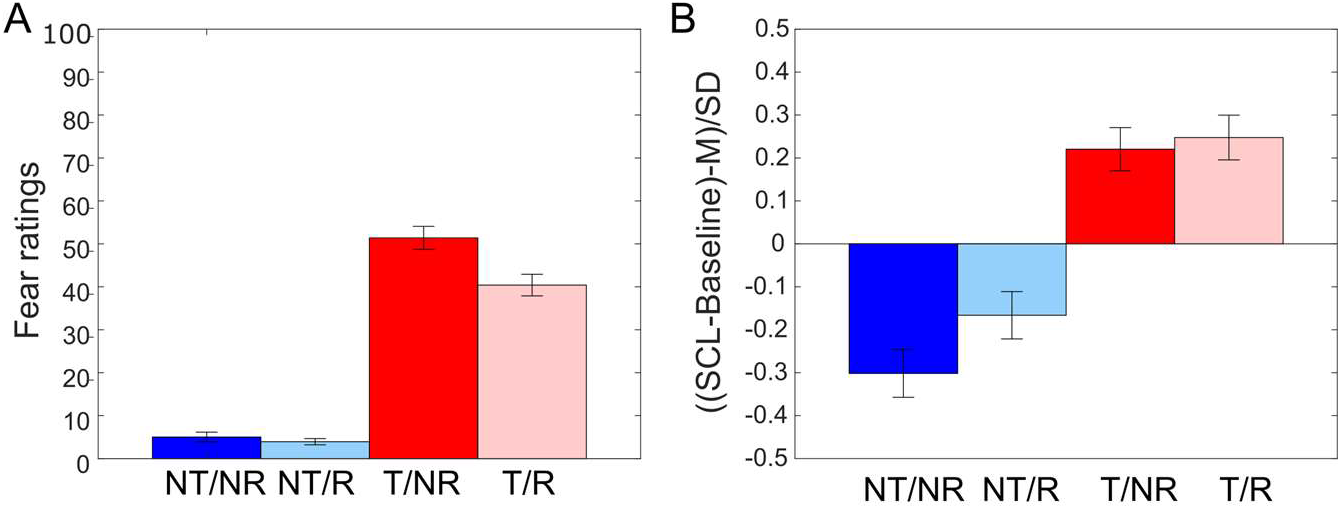
Results of Task 4 (self-focused volitional reappraisal): Fear ratings and SCL. N=54. NT, No Threat; NR, No Reappraisal; R, Reappraisal; SCL, skin conductance level; T, Threat.

#### fMRI data

ROI analyses yielded the expected strong activation in the fear network in the threat contrast T>NT and also detected typical activity in the rostral dorsal ACC/dorsomedial PFC (rdACC/ dmPFC),previously implicated in conscious threat appraisal (Kalisch & Gerlicher, 2014) (Table 13, Figure 16 A). Threat-related bilateral amygdala activation decayed quickly (parametrically modulated T>NT contrast). Safety-related activation (NT>T, categorical) was observed in the vmPFC and bilateral aHPC as well as pHPC. Reappraisal (R>NR, categorical) was associated with the typical left-lateralized (pre)frontal and temporo-parietal activation (Figure 16 B). We did not find any significant threat by reappraisal interaction that would have indicated enhanced reappraisal-related activation under threat or reduction of threat-related activation under reappraisal. We, however, report a significant simple main effect of reappraisal under threat (T/NR>T/R, categorical) in the right thalamus (Figure 16 C), in line with a dampening effect of reappraisal on threat-related processing. Additional exploratory whole-brain results are reported in the suppl. materials.

**Table 13.** Results of Task 4 (self-focused volitional reappraisal): fMRI ROI analyses.

| Contrast | Region | MNI |  |  | *z*-score | *P*<sub>svc</sub> | k |
| --- | --- | --- | --- | --- | --- | --- | --- |
| T>NT | R dACC | 2 | 14 | 28 | 5.89 | <0.001* | 866 |
|  | R rdACC/dmPFC | 6 | 34 | 22 | Inf | <0.001* | 819 |
|  | R AI | 36 | 24 | 6 | Inf | <0.001* | 878 |
|  | L AI | -32 | 20 | 6 | Inf | <0.001* | 863 |
|  | R thalamus | 4 | -20 | 0 | 6.76 | <0.001* | 110 |
|  | L thalamus | -4 | -20 | 0 | 6.24 | <0.001* | 110 |
|  | R BG | 14 | 6 | -2 | Inf | <0.001* | 51 |
|  | L BG | -12 | 2 | 0 | 7.52 | <0.001* | 36 |
| T>NT (pmod) | R amygdala | 18 | -6 | -18 | 3.59 | 0.01* | 102 |
|  | L amygdala | -22 | -6 | -18 | 3.57 | 0.01* | 97 |
| NT>T | B vmPFC | 0 | 34 | -20 | 5.18 | <0.001* | 487 |
|  | R aHPC | 36 | -24 | -14 | 5.16 | <0.001* | 254 |
|  | L aHPC | -32 | -24 | -12 | 3.57 | 0.01* | 209 |
|  | R pHPC | 34 | -28 | -12 | 4.28 | <0.001* | 59 |
|  | L pHPC | -30 | -36 | -4 | 4.25 | <0.001* | 98 |
| R>NR | L dmPFC | -4 | 14 | 54 | 4.34 | <0.001* | 176 |
|  | L SMA | -4 | 4 | 62 | 6.18 | <0.001* | 338 |
|  | L ACC | -4 | 22 | 38 | 2.82 | 0.14 | 13 |
|  | L dlPFC | -48 | 0 | 52 | 6.15 | <0.001* | 262 |
|  | L AI | -42 | 24 | -4 | 4.87 | <0.001* | 504 |
|  | L IFG | -50 | 18 | 2 | 4.36 | <0.001* | 309 |
|  | L MTG | -56 | -30 | -2 | 5.76 | <0.001* | 700 |
|  | L SPL | -40 | -56 | 36 | 2.70 | 0.21 | 8 |
| T/NR>T/R<br>(categorical) | R thalamus | 6 | -22 | 4 | 2.80 | 0.03* | 70 |

N=54. ACC, anterior cingulate cortex; AI, anterior insula; BG, basal ganglia; dlPFC, dorsolateral PFC; dmPFC, dorsomedial PFC; IFG, inferior frontal gyrus; MTG, middle temporal gyrus; rdACC, rostradorsal ACC; SMA, supplementary motor area; vmPFC, ventromedial PFC. \* significant at $p_{svc}<0.05$ .

**Figure 16.**
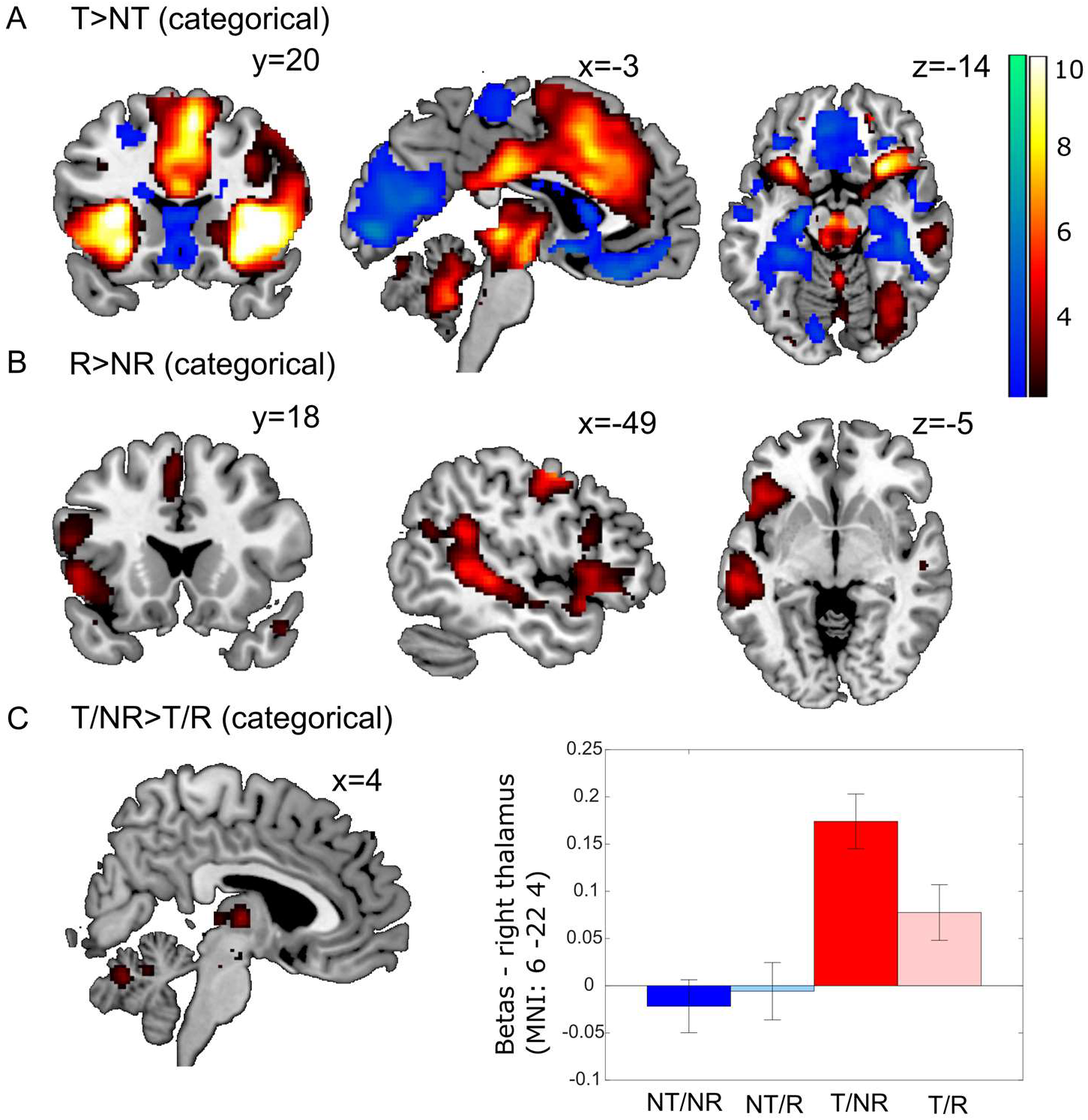
Results for Task 4 (self-focused volitional reappraisal): fMRI activations. N=54. Display threshold: p_unc_<0.01. Hot color bar (right) shows t-values for activations; cold color bar (left) shows t-values for deactivations. Bar graph shows means±SEM.

### Task 5: Situation-focused volitional reappraisal

#### Ratings

After the experiment, subjects reported to have successfully applied the reappraisal strategy (M=74 out of 100, SD=15.34, range: 24 to 100, N=53). Trial-by-trial emotional state ratings showed a main effect of valence (F_(1.24;64.43)_=292.09, p<0.001, N=53) in the expected order, with subjects feeling more happy when seeing positive [Pos] than neutral [Neu] than negative [Neg] pictures (Figure 17). The impact of reappraisal consisted in a significant main effect of reappraisal (R>NR: F_(1;52)_=295.50, p<0.001) and a significant valence by reappraisal interaction ((R>NR)Neg>(R>NR)Neu>(R>NR)Pos: F_(1.59;82.81)_=46.54, p<0.001). The interaction was mainly characterized by a higher increase in emotional state ratings for negative than for positive pictures by reappraisal (post-hoc test for (R>NR)Neg>(R>NR)Pos: F_(1;52)_=63.59, p<0.001).

**Figure 17.**
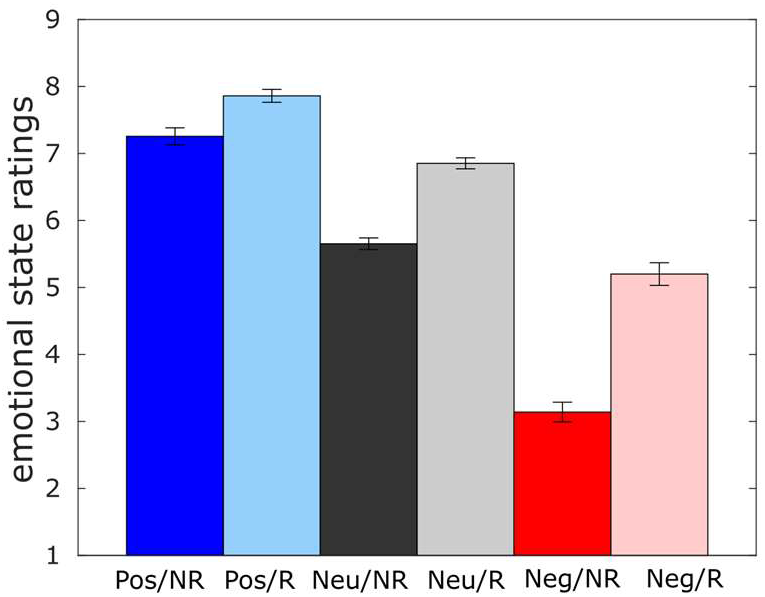
Results of Task 5 (situation-focused volitional reappraisal): Emotional state ratings. N=53. Neg, negative pictures; Neu, neutral pictures; NR, No Reappraisal; Pos, positive pictures; R, Reappraisal.

#### fMRI data

ROI analyses yielded the expected typical activation for positive picture viewing (Pos/NR>Neu/NR) in the vmPFC, but not in the NAcc; negative picture viewing (Neg/NR>Neu/NR) activated the typical network including bilateral amygdala and right inferior frontal gyrus (IFG) (Table 14, Figure 18 A, B). Across picture valences, reappraisal (R>NR) was associated with the typical left-lateralized (pre)frontal and temporo-parietal activation and also included the right NAcc (Table 14, Figure 18 C). The NAcc activation pattern (Figure 18 C) mirrored the emotional state ratings (Figure 17). We did not find significant activation reductions by reappraisal (NR>R), nor a significant valence by reappraisal interaction. Hence, in concordance with the task instructions, the reappraisal task might mainly have involved up-regulation of positive affect. Additional exploratory whole-brain results are reported in the suppl. materials.

**Table 14.** Results of Task 5 (situation-focused volitional reappraisal): fMRI ROI analyses.

| Contrast | Region | MNI |  |  | *z*-score | *P*<sub>svc</sub> | k |
| --- | --- | --- | --- | --- | --- | --- | --- |
| Pos/NR>Neu/NR | L vmPFC | -2 | 32 | -12 | 3.48 | 0.02* | 299 |
| Neg/NR>Neu/NR | R amygdala | 18 | -4 | -12 | 5.50 | <0.001 | 92 |
|  | L amygdala | -22 | -6 | -12 | 5.16 | <0.001 | 84 |
|  | R IFG | 46 | 24 | 14 | 3.88 | 0.01* | 510 |
|  | R ILOC | 50 | -60 | -2 | 5.36 | <0.001 | 607 |
|  | L ILOC | -44 | -70 | -4 | 5.70 | <0.001 | 666 |
|  | R brainstem | 2 | -30 | -2 | 4.49 | <0.001 | 95 |
|  | L brainstem | -8 | -28 | -10 | 5.47 | <0.001 | 113 |
| R>NR | L dmPFC | -10 | 24 | 62 | 7.25 | <0.001 | 575 |
|  | L SMA | -4 | 8 | 62 | Inf | <0.001 | 623 |
|  | R ACC | 10 | 18 | 32 | 3.50 | 0.02* | 829 |
|  | L dlPFC | -44 | 2 | 54 | Inf | <0.001 | 559 |
|  | L AI | -34 | 10 | 0 | 4.56 | <0.001 | 844 |
|  | R IFG | 54 | 26 | -4 | 3.61 | 0.02* | 114 |
|  | L IFG | -54 | 18 | 16 | 6.22 | <0.001 | 478 |
|  | L MTG | -50 | -34 | -2 | Inf | <0.001 | 778 |
|  | L SPL | -42 | -60 | 32 | 5.74 | <0.001 | 393 |
|  | R NAcc | 12 | 6 | -4 | 5.05 | <0.001 | 40 |

N=52. ACC, anterior cingulate cortex; AI, anterior insula; IFG, inferior frontal gyrus; dlPFC, dorsolateral PFC; dmPFC, dorsomedial PFC; ILOC, inferior lateral occipital cortex; MTG, middle temporal gyrus; NAcc, nucleus accumbens; SPL, superior parietal lobule; vmPFC, ventromedial PFC. \* significant at *p*<sub>svc</sub><0.05.

**Figure 18.**
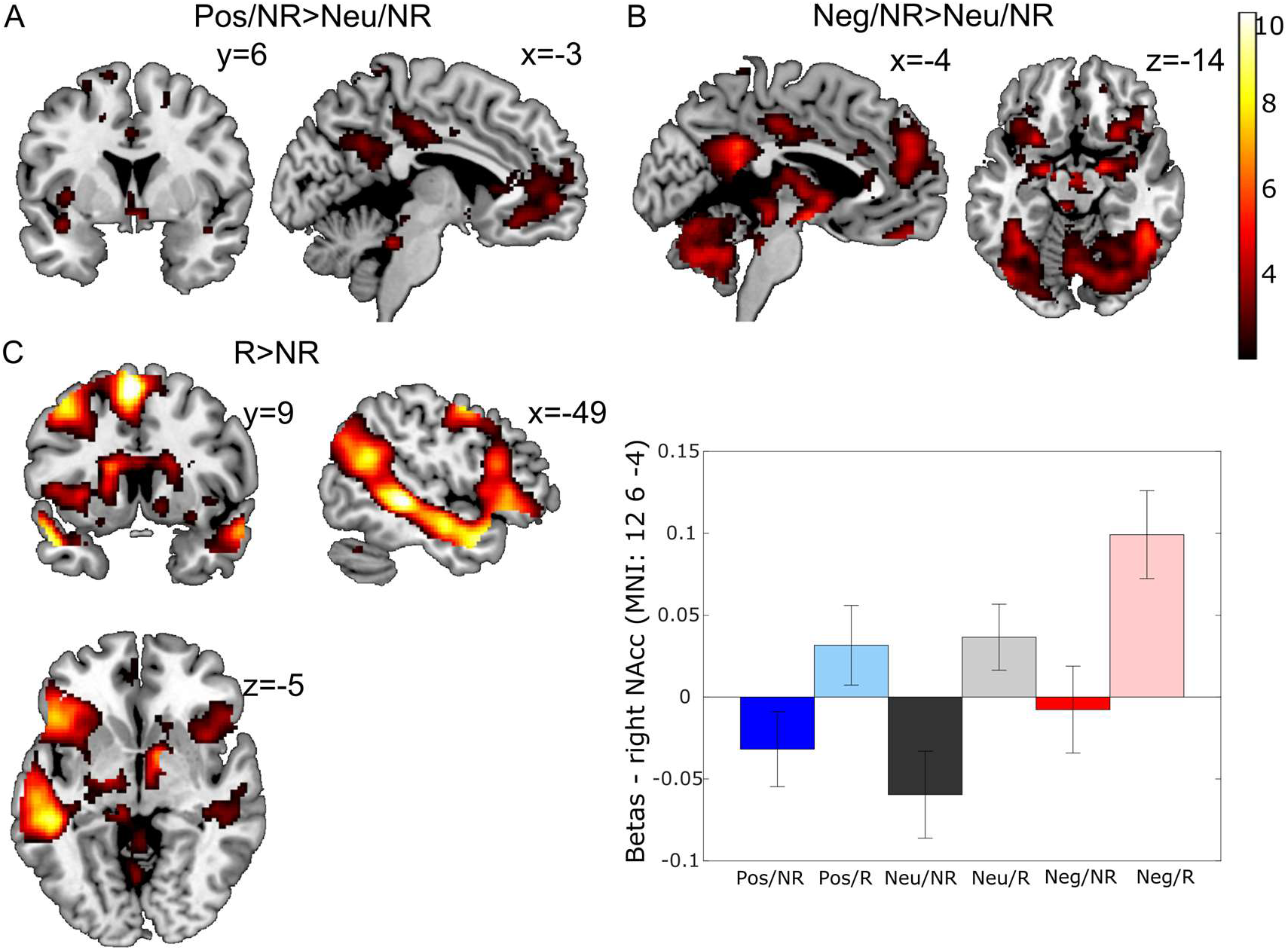
Results for Task 5 (situation-focused volitional reappraisal): fMRI activations. N=52. Display threshold: p_unc_<0.01. Color bar shows t-values. Bar graph shows means±SEM.

### Task 6: Emotional interference and motor response inhibition

#### Behavioral data

In the spatial interference inhibition (Simon) task, the rates of omissions and errors (wrong answers) during Incongruent Go (Incon Go) trials were generally low (<2%), and the rates did not differ significantly between negatively (Neg) and neutrally (Neu) primed trials (t_omission_=1.07, p=0.290; t_errors_=0.39, p=0.696; N=47). Repeated-measures ANOVA on RTs (Table 15) revealed a significant main effect of spatial interference (Incon Go>Con Go) (F=133.71, p<0.001; N=47), a significant main effect of negative emotion (Neg>Neu) (F=7.56, p=0.009), but no significant interaction between spatial interference and negative emotion (F=0.23, p=0.637). Hence, there was no evidence that the emotional primes interfered with motor response inhibition in this task. In the action cancellation (stop signal) task, the targeted rate of commission errors (50%) during Stop trials was well approximated, with 52% of subjects succeeding to stop (correct stops: SD=0.5, range: 44 to 68%). Subjects made thus more commission errors after negative primes (see Table 15). In addition, negative primes prolonged SSRTs (Table 15), indicating that negative emotion interfered with (slowed down) both going and stopping processes in the stop signal task.

**Table 15.** Results of Task 6 (emotional interference and motor response inhibition): Behavior

|  | Neu<br>*M* | *SD* | Neg<br>*M* | *SD* | *t-score* | *p* |
| --- | --- | --- | --- | --- | --- | --- |
| RT (Con Go) | 618.68 | 86.88 | 623.42 | 84.68 | 2.61 | 0.01* |
| RT (Incon Go) | 669.49 | 85.42 | 673.28 | 85.09 | 2.03 | 0.05* |
| Correct stops | 52.72% | 5.26% | 51.93% | 5.45% | -2.31 | 0.03* |
| SSRT | 372.95 | 54.98 | 383.83 | 58.85 | 3.16 | 0.00* |

N=47. Neg, negatively primed trials; Neu, neutrally primed trials; SSRT, stop signal reaction time.
\* significant at $p < 0.05$ .

#### fMRI data

Spatial interference inhibition (Incon Go>Con Go) and action cancellation (Stop>Con Go) both activated the typical common motor response inhibition network including pre-SMA, bilateral AI and right IFG, pars opercularis (Table 16, Figure 19 A,B). Spatial interference inhibition-specific activations were present bilaterally in the dlPFC and superior parietal lobule (SPL); action cancellation-specific activations were found in the right ACC and IFG, pars triangularis, as well as the bilateral subthalamic nucleus (STN; Table 16). Negative primes (Neg>Neu) induced comparable activation as the Neg/NR>Neu/NR contrast in Task 5 (Table 16, Figure 19 C). The right amygdala showed an interaction of motor response inhibition with emotion in that it was significantly more reactive to the negative emotional primes in the control (Con Go) condition as compared to the two motor response inhibition conditions (Incon Go, Stop) (Table 16, Figure 19 D). This suggests that motor response inhibition dampened emotional processing, in line with the employment of cognitive control. There was, however, no motor response inhibition-related area activated more strongly on Incon Go and Stop than Con Go trials specifically after negative primes (i.e., inverse interaction). For exploratory whole-brain results, see suppl. materials.

**Table 16.**
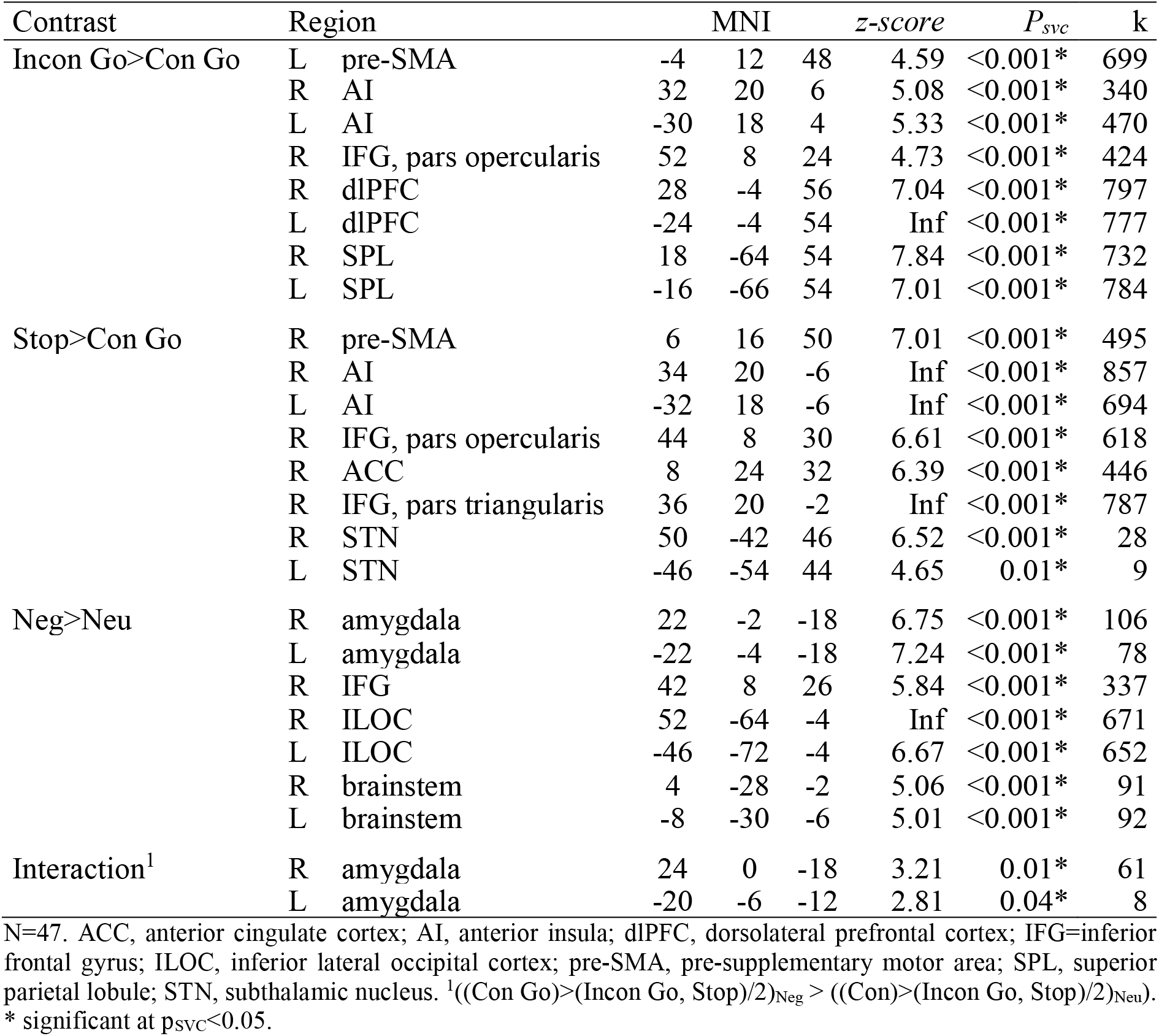
Results of Task 6 (emotional interference and motor response inhibition): fMRI ROI analyses

| Contrast | Region | MNI | | | z-score | $P_{svc}$ | k |
| --- | --- | --- | --- | --- | --- | --- | --- |
| Incon Go>Con Go | L pre-SMA | -4 | 12 | 48 | 4.59 | <0.001* | 699 |
|  | R AI | 32 | 20 | 6 | 5.08 | <0.001* | 340 |
|  | L AI | -30 | 18 | 4 | 5.33 | <0.001* | 470 |
|  | R IFG, pars opercularis | 52 | 8 | 24 | 4.73 | <0.001* | 424 |
|  | R dlPFC | 28 | -4 | 56 | 7.04 | <0.001* | 797 |
|  | L dlPFC | -24 | -4 | 54 | Inf | <0.001* | 777 |
|  | R SPL | 18 | -64 | 54 | 7.84 | <0.001* | 732 |
|  | L SPL | -16 | -66 | 54 | 7.01 | <0.001* | 784 |
| Stop>Con Go | R pre-SMA | 6 | 16 | 50 | 7.01 | <0.001* | 495 |
|  | R AI | 34 | 20 | -6 | Inf | <0.001* | 857 |
|  | L AI | -32 | 18 | -6 | Inf | <0.001* | 694 |
|  | R IFG, pars opercularis | 44 | 8 | 30 | 6.61 | <0.001* | 618 |
|  | R ACC | 8 | 24 | 32 | 6.39 | <0.001* | 446 |
|  | R IFG, pars triangularis | 36 | 20 | -2 | Inf | <0.001* | 787 |
|  | R STN | 50 | -42 | 46 | 6.52 | <0.001* | 28 |
|  | L STN | -46 | -54 | 44 | 4.65 | 0.01* | 9 |
| Neg>Neu | R amygdala | 22 | -2 | -18 | 6.75 | <0.001* | 106 |
|  | L amygdala | -22 | -4 | -18 | 7.24 | <0.001* | 78 |
|  | R IFG | 42 | 8 | 26 | 5.84 | <0.001* | 337 |
|  | R ILOC | 52 | -64 | -4 | Inf | <0.001* | 671 |
|  | L ILOC | -46 | -72 | -4 | 6.67 | <0.001* | 652 |
|  | R brainstem | 4 | -28 | -2 | 5.06 | <0.001* | 91 |
|  | L brainstem | -8 | -30 | -6 | 5.01 | <0.001* | 92 |
| Interaction<sup>1</sup> | R amygdala | 24 | 0 | -18 | 3.21 | 0.01* | 61 |
|  | L amygdala | -20 | -6 | -12 | 2.81 | 0.04* | 8 |

N=47. ACC, anterior cingulate cortex; AI, anterior insula; dlPFC, dorsolateral prefrontal cortex; IFG=inferior frontal gyrus; ILOC, inferior lateral occipital cortex; pre-SMA, pre-supplementary motor area; SPL, superior parietal lobule; STN, subthalamic nucleus. <sup>1</sup> $((\text{Con Go}) > (\text{Incon Go, Stop})/2)_{\text{Neg}} > ((\text{Con}) > (\text{Incon Go, Stop})/2)_{\text{Neu}}$ .
\* significant at $p_{svc} < 0.05$ .

**Figure 19.**
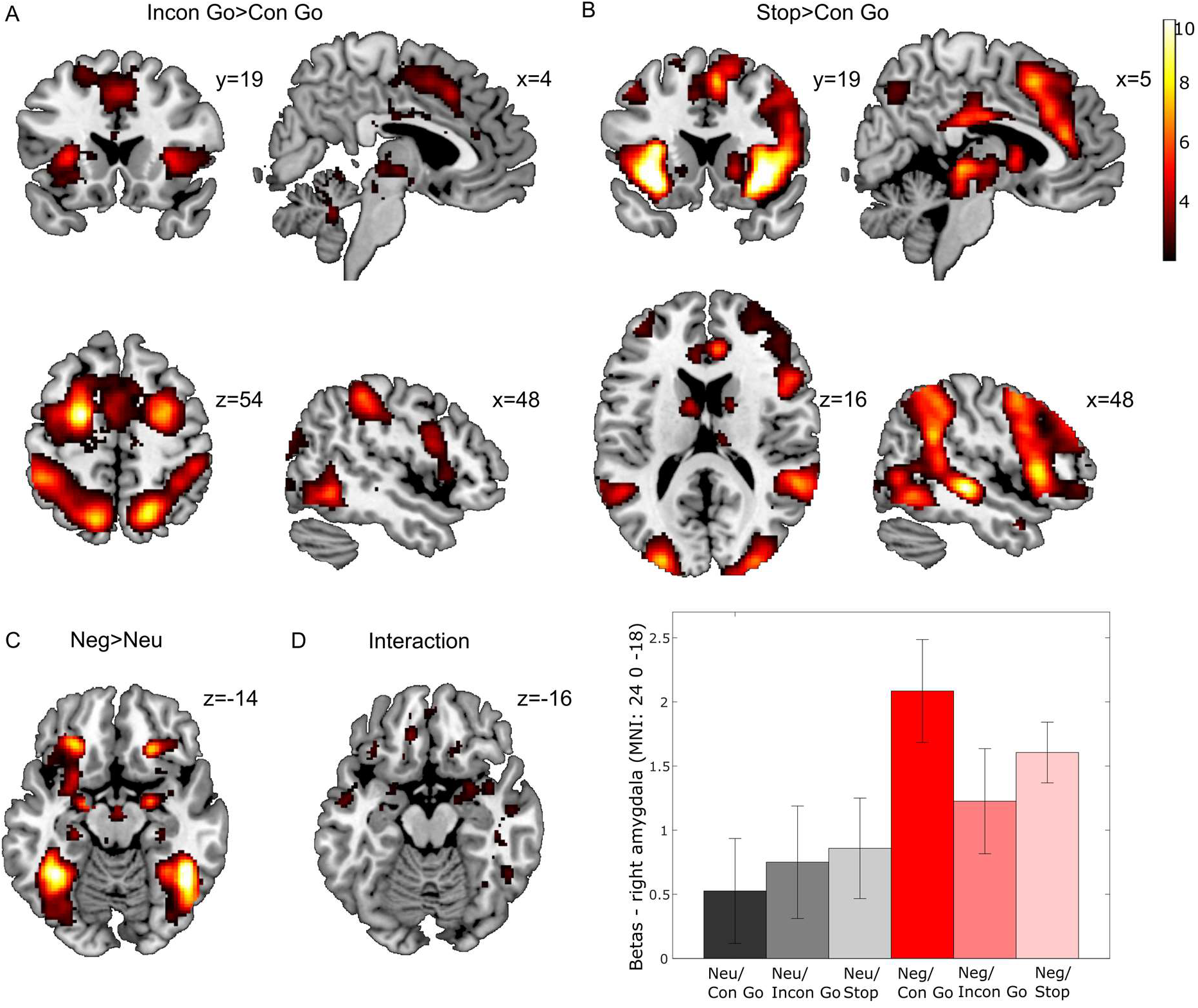
Results for Task 6 (emotional interference and motor response inhibition): fMRI activations. N=47. Display threshold: p_unc_<0.01. Color bar shows t-values. Bar graph shows means±SEM.

## Discussion

### Developing a test battery for the longitudinal study of resilience

Based on a theoretical framework of resilience (Kalisch et al., 2015), a multi-modal test battery was developed that measures subjective-experiential, behavioral, physiological and neural reactions to various types of aversive stimulation, with a focus on scenarios that involve a change of stimulus meaning (e.g., because a stimulus terminates or is no longer threatening), require subjects to deliberately change stimulus meaning (volitional reappraisal), or necessitate the inhibition of irrelevant aversive stimuli. By thus assessing the efficiency and effectiveness of subjects’ stress response regulation, the battery is assumed to inform about the efficiency and effectiveness of the underlying positive (i.e., non-negative) (re)appraisal processes in three broad classes of neuro-cognitive processes (positive situation classification, reappraisal proper, interference inhibition). Different aversive stimulus types are employed (performance pressure, negative pictures, unpleasant and distracting sensory stimulation, pain) to help average out potential idiosyncratic responses a subject might show to a specific stimulus type, based for instance on previous experience with such stimuli. Factor analysis of the battery should reveal latent hidden variables that would ideally span more than one task, type of aversive stimulation and measurement modality and should be interpretable as reflecting key (re)appraisal processes. These in turn should explain interindividual differences in resilience in longitudinal studies.

We here for the first time applied the battery in a sample of healthy young participants (N=55) to test its feasibility and usefulness. We consider the battery to be feasible, based on overall acceptable dropout rates. One subject dropped out before MRI due to claustrophobia. Restriction to subjects without MRI-related fears is an inherent feature of neuroimaging studies and constitutes a limitation in so far as such studies may be blind for a group of individuals with a specific psychological vulnerability. All 54 remaining subjects finished the battery. Between 47 to 54 of the fMRI task data sets could be analyzed (see Table 6). The dropout rates are in a range reported by other studies using neuroimaging batteries with emotional tasks in healthy normal subjects (e.g., Roalf et al., 2014; Rohr et al., 2015). Given the long duration of our procedures and the stressful nature of several of the tasks, it is however conceivable that dropout rates might be higher in older subjects or in subjects selected for particular stress-related vulnerabilities. This would motivate a more condensed battery. A pragmatic approach for future applications could be to use information about the most predictive battery factors in the on-going prospective-longitudinal resilience study MARP to select only those tasks or task components that are strongly related to resilience. Dropout rates in non-neural measures ranged from 13 subjects (heart rate in Task 1) to 0 (ratings in Task 4) and were mostly in the order of 5 or less subjects. Since each task involves several outcomes measures (several non-neural outcomes in Task 1, fMRI and at least one non-neural outcome in all other tasks), we deem these rates acceptable.

For the purpose of testing the battery’s usefulness, we asked whether task-based fMRI activations are in agreement with results from previous studies with comparable tasks, as published in recent meta-analyses, informal reviews and relevant single studies. We did not attempt to exactly replicate identical earlier experiments; therefore, any judgement given here is qualitative and served us to decide whether or not the identical battery should be used in MARP. (By contrast, quantitative judgments will be possible based on an exact replication of the present results in N=103 MARP subjects, which will be reported elsewhere.) In the following, we briefly discuss the results from each task in this respect. On the basis of this discussion, we judge the battery to globally replicate the existing literature, and we consequentially decided to use the identical battery in MARP. The employment of an identical battery in MARP has the significant advantage that we can use information about the factor structure of the battery to be derived from the current sample to also analyze battery data from the MARP sample.

Task 1 (MMST; adapted from Reinhardt et al., 2012) of the battery investigates stress reactivity and recovery. We replicated the findings by Reinhardt et al. (2012) showing substantial stress responses across different outcome measures including stress ratings, heart rate, skin conductance level (SCL) and responses (SCR), and salivary cortisol and α-amylase. Except for SCL we captured a full recovery within our experimental session for all outcome measures.

Task 2 (MID; adapted from Knutson et al., 2001; Wu et al., 2014) assesses reward sensitivity. As expected, the chance to win money led to reduced reaction times and marginally enhanced hit rates. Congruent with our predictions, the NAcc was activated by both gain and loss anticipation as well as by the realization of gains and the prevention of losses during the outcome phase (Bartra et al., 2013). Also in congruence with former results (Wu et al., 2014), the anticipation of gains and losses also recruited the bilateral AI. Note that gain and loss anticipation activated multiple brain networks outside our ROIs, spanning from prefrontal to occipital cortex, which can be explained by the diversity of processes (attention, outcome prediction, motor response preparation) co-occurring during this phase of the task (Jia et al., 2016). In line with Bartra et al. (2013), we found the vmPFC to specifically activate when subjects anticipated or received a gain. Interestingly, whole-brain activation maps for gain outcomes closely resembled the profile of the default mode network (DMN) (Greicius, Krasnow, Reiss, & Menon, 2003), including not only the vmPFC, but also posterior cingulate cortex and precuneus. This corresponds to a meta-analysis by Liu, Hairston, Schrier, & Fan (2011), who have shown that the activation pattern for the receipt of a reward closely resembles the DMN. Unexpectedly, both no gain and loss outcomes did not recruit bilateral AI and dACC (Bartra et al., 2013; Liu et al., 2011), the brain areas most consistently activated by aversive stimuli (Mechias et al., 2010), in our sample. In general, a low reactivity to negative outcomes in this task might be related to the relatively low aversiveness of these stimuli compared to the aversiveness of the salient negative pictures or the threat of pain also present in the battery.

Safety learning is tested in Task 3 by means of differential Pavlovian fear conditioning, subsequent extinction and – on the next day - a memory test (adapted from Milad et al., 2007). Fear ratings and SCR data proved successful conditioning, i.e., CS+>CS− stimulus discrimination. Activation patterns largely corresponded to the results of a recent meta-analysis (Fullana et al., 2016), revealing activations in bilateral AI, pre-SMA, dACC, bilateral thalamus and basal ganglia for the CS+. Additionally, we observed CS+ related activation in a amygdala-hippocampal transition zone that gradually decreased over the course of conditioning (Plichta et al., 2014). vmPFC and OFC were deactivated relative to the CS− (Fullana et al., 2016).

Successful extinction was indicated by initial stimulus discrimination in the fear ratings that dissolved until the end of extinction (stimulus by time interaction). Contrary to our prediction, we did not find stimulus discrimination or stimulus by time effects in the SCR and fMRI data, but merely a general effect of time. From the start to the end of extinction, activation in bilateral AI, dACC and right amygdala decreased. One explanation for the lack of stimulus discrimination in extinction could be a general increase in arousal, obscuring differential stimulus effects, when starting a new task run after a break used for resting-state fMRI that further involved a change of context (background picture) relative to the previous conditioning run. Additionally, the abrupt change from a 100% to a no reinforcement schedule might have accelerated extinction learning (Lonsdorf et al., 2017; Vansteenwegen, Crombez, Baeyens, & Eelen, 1998), making detection of relatively larger CS+ responses even more difficult. Nonetheless, in the spontaneous recovery test the next day, subjects showed higher fear ratings as well as higher activation in regions of the fear network, namely left amygdala, right AI, right basal ganglia and left thalamus, to the unextinguished (CS+u) than to the extinguished CS+ (CS+e). A relative reduction in responding to the extinguished CS implicates successful retrieval of extinction memory.

The re-occurrence of the threat-related conditioning context A in the subsequent renewal test led to a general return of fear, indicated by increased fear ratings and SCRs and a higher activation in left AI to both CS+s compared to the CS−. As in conditioning, the CS− was characterized by a significant relative activation in vmPFC, right OFC and now also right anterior and posterior hippocampus. A loss of discrimination between the extinguished and the unextinguished CS may reflect the difficulty to retrieve an extinction memory in the threat-related conditioning context. A test order effect might play an additional role: since spontaneous recovery served as opportunity for extinction learning to the CS+u, initial stimulus discrimination might have been lost by the start of renewal.

In Task 4, subjects have to apply self-focused volitional reappraisal in an instructed fear (“threat of shock”) paradigm (adapted from Paret et al., 2011). Threat induced activations and deactivations in the same canonical fear and safety networks that were already observed in Task 3 (see Fullana et al., 2016; Mechias et al., 2010) and also included decaying amygdala activity. Additionally, threat recruited the rdACC/dmPFC, a region known to be involved in conscious threat appraisal (Kalisch & Gerlicher, 2014). Applying reappraisal by distancing oneself from the experimental situation had an anxiolytic effect, as indicated by reduced fear ratings. Contrary to our prediction, it did not lead to a reduction in SCL. Former tasks that succeeded in modulating SCL by reappraisal used trials of longer duration (Kalisch et al., 2005; Paret et al., 2011), thus giving subjects more time to apply the strategy and also reducing the chance that consecutive trials confound SCL measurement. An alternative explanation is that the cognitive effort likely to be associated with volitional reappraisal (as suggested by the numerically positive reappraisal main effect on SCL) might have masked SCL reductions in the threat condition. Corresponding to former meta-analyses (Buhle et al., 2013; Frank et al., 2014; Kalisch, 2009; Kohn et al., 2014), applying cognitive volitional reappraisal activated a left-sided network including ACC, pre-SMA, dmPFC, left dlPFC, left AI, left IFG, left middle temporal gyrus and left superior parietal lobule. Note that we did not predict right-sided activation because of the short trial duration (Kalisch, 2009). Unexpectedly, there were no activity reductions in threat-related areas by reappraisal, with the exception of reduced thalamus activity, the latter presumably reflecting a globally dampening effect of the distancing strategy, irrespective of whether subjects are threatened or not. Globally dampening effects are generally observed in behavioral distancing studies and include the effectiveness of distancing in also down-regulating positive emotion (e.g., Park et al., 2014; Verduyn, Van Mechelen, Kross, Chezzi, & Van Bever, 2012). In situation-focused volitional reappraisal in Task 5, subjects have to positively reinterpret visually displayed social situations of positive, neutral and negative valence. Subjects showed the typical subjective and fMRI responses to the pictures (Carretié, 2014; García-García et al., 2016). As expected, reappraisal made subjects feel more positive when viewing the pictures, an effect that was most pronounced for negative pictures. Reappraisal-related activations were comparable to Task 4, with an additional NAcc activation that we predicted based on the nature of the reappraisal strategy that emphasized the generation of positive interpretations. The unexpected absence of activity reductions in negative picture-responsive brain areas by reappraisal might also be related to the nature of the strategy. In line with this, a recent metaanalysis revealed that amygdala activation was only reduced when subjects were instructed to downregulate emotion (Buhle et al., 2013).

In Task 6, negative visual distractors are used as primes to affect performance on a well-established motor response inhibition task (HRI; Sebastian et al., 2013). The spatial interference inhibition (Simon) component of the HRI yielded the described slowing of reactions in incongruent relative to congruent go trials (Sebastian et al., 2013) as well as the described recruitment of AI, dlPFC, IFG, superior parietal lobule and pre-SMA (Cieslik et al., 2015). The action cancellation (stop signal) component activated a largely overlapping set of brain regions, with both tasks also showing a limited number of expected task-specific activations (Cieslik et al., 2015; Sebastian et al., 2013, 2016; Aron & Poldrack, 2006). Negative primes slowed both going and stopping processes (Carretié, 2014; Etkin, Egner, Peraza, Kandel, & Hirsch, 2006; Kalanthroff, Cohen, & Henik, 2013; Verbruggen & De Houwer, 2007) and induced activity comparable to the negative pictures in Task 5, including in the amygdala. Amygdala reactivity to negative pictures was higher in the congruent go than in the incongruent go and stop trials. This emotion by motor response inhibition interaction effect can be interpreted as dampening of emotional reactivity by the concurrent performance of a response inhibition task. This is in line with former studies that have shown a reduction in emotional reactivity in cognitive tasks by increased cognitive load (Erk, Kleczar, & Walter, 2007; Kalisch, Wiech, Critchley, & Dolan, 2006; Kellermann et al., 2012; Wessa, Heissler, Schönfelder, & Kanske, 2013). The pattern of reduced emotional processing can be interpreted as an effort to stabilize cognitive functioning. Unexpectedly, we did not find compensatory changes in prefrontal activation (Iordan & Dolcos, 2015; Iordan, Dolcos, & Dolcos, 2013; Wessa et al., 2013).

### Conclusion and Outlook

Resilience research, in particular in humans, is still mainly phenomenological and restricted to collecting factors that are statistically related to resilient outcomes. Reviews enumerate long lists of such resilience factors. Apart from intensity and quality of the stressors, these lists include external factors, such as social support or socioeconomic status, as well as internal or personality factors (e.g., life history, certain character traits, coping style, age, sex, ethnicity, (epi)genetics, spirituality, cognitive abilities, hormonal factors etc.); these factors are often complemented by their interactions (e.g., Feder, Charney, & Abe, 2011; Mancini & Bonanno, 2009; Sapienza & Masten, 2011; Southwick & Charney, 2012; Stewart & Yuen, 2011). These lists are lengthy and cumbersome, and it has been pointed out that many of the listed factors overlap conceptually and presumably mediate or depend on each other (Stewart & Yuen, 2011). It is therefore necessary for resilience research to proceed from collecting a never ending list of resilience factors to identifying and understanding the mediating mechanisms, i.e., a presumably limited number of shared cognitive and/or neural pathways that provide protection against stress-related impairments (Kalisch et al., 2015). From this mechanistic perspective, the psychosocial, genetic and other resilience factors listed above would only affect resilience via these mediating mechanisms, which we term ‘resilience mechanisms’. The core of resilience research should therefore be the identification, understanding, and exploitation of these resilience mechanisms. It can be hoped that interventions targeting these mechanisms might be particularly efficient in preventing stress-related dysfunction. The current investigation is intended to contributing to this long-term goal.

Another issue in current resilience research is that the majority of existing resilience studies use retrospective cross-sectional designs, e.g., comparing traumatized subjects with psychopathology to traumatized subjects without psychopathology (i.e., resilient subjects) in terms of brain or cognitive function or other potential resilience mechanisms. To infer causality and rule out pre-existing group differences, prospective-longitudinal studies are needed that assess possible resilience mechanisms before and after stressor exposure, as well as ideally also during stressor exposure. Such studies also require a thorough monitoring of stressor exposure and psychological health, again ideally at multiple time points, to eventually yield insights into the dynamic adaptation processes underlying resilience (Kalisch et al., 2017). To this end, we currently apply the described task battery repeatedly every 1.75 years in a longitudinal study, the Mainz Resilience Project (MARP), in a group of young adults with significant previous adverse life experiences who find themselves in the transition from adolescence and family and school life to adulthood and work or academic life. This critical life transition (“leaving home”) is known to be associated in many individuals with a range of new challenges and with the onset or exacerbation of stress-related mental problems (Compas, Wagner, Slavin, & Vannatta, 1986; Craske et al., 2012; Herbst, Voeth, Eidhoff, Müller, & Stief, 2016; Kessler et al., 2005; Reavley & Jorm, 2010; Rotenstein et al., 2016). In addition to the test battery, stressor exposure and mental health is assessed every 3 months using online tools. This dense assessment and thorough phenotyping renders it possible to appropriately quantify resilience (Kalisch et al., 2017) and relate it to predictors from the test battery.

The latter is, however, complicated by the fact that our test battery is extended and multi-modal and therefore contains a large number of potential predictor variables that far exceed the number of subjects that can be reasonably tested with the battery. This requires statistical methods for data integration, data reduction and evenutally identification of a limited number of latent battery variables, or factors. These on-going analysis efforts might not only yield good predictors but also permit in the future to prune the battery, leaving only those tasks or elements that strongly contribute to the predictor variables. We again emphasize that we expect those predictor variables to change over time, reflecting adaptation.

Concluding, we have developed a test battery that we hope will help uncover the underlying structure of resilience mechanisms. It can be applied in longitudinal stress resilience studies for a systematic investigation of resilience mechanisms. One major strength of the test battery is its theoretical foundation. Finally, with its neuro-cognitive focus, it promises to contribute to the identifican of new targets for prevention.

## Acknowledgement

We wish to thank Petra Seyfarth and Manuela Götz for their engagement in subject recruitment, data acquisition and study organization. Further help was provided by Thomas Bauermann, Hanno Burger, Samira Christmann, Haakon Engen, Sarah Mohr, Alexander Schüler, Goran Vucurevic, Merle Wachendörfer and Vanessa Zörrer.

This work was funded by Stiftung Rheinland-Pfalz für Innovation (MARP program, No 961-386261/1080), the Ministry of Science of the state of Rhineland-Palatinate (DRZ program) and the European Union’s Horizon 2020 research and innovation program (grant agreement No 777084).

## Footnotes

1 The following contingency questions were asked after all phases: Did you receive a pain stimulation in this phase (yes/no)? Specifically after CD: Did you know when you would receive a pain stimulus (yes/no)? For each symbol, after all phases: Was this symbol coupled with a pain stimulus (yes/no)? Specifically after CD and EX: In which of these two contexts [shown below] were the symbols presented to you?

2 The four self statements were as follows (translated into English): “The situation does not directly concern me.”, “I observe my reactions in a distant and neutral fashion,”, “I am observing how the experiment develops.”, “I experience the situation from far away.”

3 The following pictures were selected for Experiment 5: EmoPicS: 55, 56, 57, 66, 74, 75, 93, 106, 108, 109, 114, 129, 133, 160, 162, 211, 233, 235,239, 244, 251, 252, IAPS: 2411, 3350, 4597, 4599, 8158, 8496, 9520, 9530

4 The following pictures were selected for Experiment 6: EmoPicS 83, 84, 86, 87, 88, 90, 91, 92, 104, 105, 110, 112, 113, 116, 120, 121, 122, 123, 124, 127, 130, 131, 132, 135, 137, 139, 142, 143, 157, 158, 159, 167, 172, 175, 180, 181, 183, 185, 186, 187, 188, 190, 191, 192, 193, 195, 199, 201, 202, 236, 237, 240, 241, 242, 243, 269, 276, 278, 280, 281, 285, 286, 289, 301, 302, 304, 306, 309, 310, 311, 312, 313, 319, IAPS: 2002, 2026, 2101, 2102, 2104, 2190, 2191, 2200, 2206, 2210, 2221, 2272, 2273, 2305, 2308, 2375.1, 2377, 2385, 2390, 2397, 2440, 2441, 2445, 2446, 2484, 2499, 2512, 2516, 2525, 2750, 2811, 3001, 3016, 3017, 3030, 3051, 3053, 3059, 3060, 3061, 3063, 3064, 3069, 3080, 3100, 3101, 3102, 3103, 3110, 3120, 3130, 3131, 3170, 3185, 3195, 3225, 3300, 5731, 6230, 6250, 6260, 6415, 6563, 6831, 7001, 7014, 7018, 7019, 7033, 7043, 7045, 7052, 7055, 7057, 7058, 7059, 7061, 7062, 7081, 7500, 7512, 7513, 7546, 7547, 7710, 8312, 9000, 9040, 9070, 9075, 9183, 9187, 9252, 9253, 9260, 9265, 9412, 9413, 9414, 9420, 9432, 9433, 9500, 9570, 9571, 9600, 9610, 9611, 9620, 9700, 9900, 9901, 9903, 9904, 9905, 9908, 9910, 9911, 9920, NAPS: Animals_001_h. 008_v, 013_h, 016_h, 019_h, 024_h, 025_h, 027_h, 032_h, 033_h, 038_h, 039_h, 048_h, 054_h, 056_h, 057_h, 062_h, 063_h, 064_v, 067_h, 068_h, 071_h, 074_h, 077_h, 078_h/ Faces_007_h, 010_h, 016_h, 018_h, 019_h, 028_h, 143_v, 145_v, 147_v, 149_v, 152_h, 153_v, 159_h, 170_h, 172_h, 363_v, 364_v, 365_v, 366_h, 367_h, 368_h, 371_v/ Landscapes_002_h, 007_h, 022_h, 026_h, 139_h/ 0bjects_001_h, 002_h, 003_h, 283_h, 285_h/ People_003_h, 004_h, 008_h, 009_h, 013_v, 016_h, 020_h, 021_h, 022_h, 031_v, 038_h, 039_v, 128_h, 140_h, 198_h, 200_h, 201_v, 205_v, 211_v, 218_v, 221_h, 225_h, 226_h, 227_h, 233_h, 235_h, 238_h, 240_h, 241_h, 242_v, 243_h

## References

Adler, N. E., Epel, E. S., Castellazzo, G., & Ickovics, J. R. (2000). Relationship of subjective and objective social status with psychological and physiological functioning: Preliminary data in healthy, White women. Health Psychology, 19(6), 586–592. https://doi.org/10.1037/0278-6133.19.6.586

Arnold, M. B. (1969). Emotion, motivation, and the limbic system. Annals of the New York Academy of Sciences, 159(3 Experimental), 1041–1058. https://doi.org/10.1111/j.1749-6632.1969.tb12996.x

Aron, A. R., & Poldrack, R. A. (2006). Cortical and Subcortical Contributions to Stop Signal Response Inhibition: Role of the Subthalamic Nucleus. Journal of Neuroscience, 26(9), 2424–2433. https://doi.org/10.1523/JNEUROSCI.4682-05.2006

Bartra, O., McGuire, J. T., & Kable, J. W. (2013). The valuation system: A coordinate-based meta-analysis of BOLD fMRI experiments examining neural correlates of subjective value. NeuroImage, 76, 412–427. https://doi.org/10.1016/j.neuroimage.2013.02.063

Boks, M. P., Mierlo, H. C. van, Rutten, B. P. F., Radstake, T. R. D. J., De Witte, L., Geuze, E., … Vermetten, E. (2015). Longitudinal changes of telomere length and epigenetic age related to traumatic stress and post-traumatic stress disorder. Psychoneuroendocrinology, 51, 506–512. https://doi.org/10.1016/j.psyneuen.2014.07.011

Bonanno, G. A., Kennedy, P., Galatzer-Levy, I. R., Lude, P., & Elfström, M. L. (2012). Trajectories of resilience, depression, and anxiety following spinal cord injury. Rehabilitation Psychology, 57(3), 236–247. https://doi.org/10.1037/a0029256

Bonanno, G. A., Mancini, A. D., Horton, J. L., Powell, T. M., LeardMann, C. A., Boyko, E. J., … Smith, T. C. (2012). Trajectories of trauma symptoms and resilience in deployed US military service members: prospective cohort study. The British Journal of Psychiatry, 200(4), 317–323. https://doi.org/10.1192/bjp.bp.111.096552

Bonanno, G. A., Romero, S. A., & Klein, S. I. (2015). The Temporal Elements of Psychological Resilience: An Integrative Framework for the Study of Individuals, Families, and Communities. Psychological Inquiry, 26(2), 139–169. https://doi.org/10.1080/1047840X.2015.992677

Bosch, J. A., Veerman, E. C. I., de Geus, E. J., & Proctor, G. B. (2011). α-Amylase as a reliable and convenient measure of sympathetic activity: don’t start salivating just yet! Psychoneuroendocrinology, 36(4), 449–453. https://doi.org/10.1016/j.psyneuen.2010.12.019

Breen, M. S., Maihofer, A. X., Glatt, S. J., Tylee, D. S., Chandler, S. D., Tsuang, M. T., … Woelk, C. H. (2015). Gene networks specific for innate immunity define post-traumatic stress disorder. Molecular Psychiatry, 20(12), 1538–1545. https://doi.org/10.1038/mp.2015.9

Brett, M., Anton, J.-L., Valabregue, R., & Poline, J.-B. (2002). Region of interest analysis using an SPM toolbox [abstract]. Presented at the 8th International Conference on Functional Mapping of the Human Brain, Sendai, Japan: Available on CD-ROM in NeuroImage, Vol 16, No 2.

Buhle, J. T., Silvers, J. A., Wager, T. D., Lopez, R., Onyemekwu, C., Kober, H., … Ochsner, K. N. (2013). Cognitive Reappraisal of Emotion: A Meta-Analysis of Human Neuroimaging Studies. Cerebral Cortex. https://doi.org/10.1093/cercor/bht154

Carretié, L. (2014). Exogenous (automatic) attention to emotional stimuli: a review. Cognitive, Affective, & Behavioral Neuroscience, 14(4), 1228–1258. https://doi.org/10.3758/s13415-014-0270-2

Carver, C. S. (1997). You want to measure coping but your protocol’s too long: Consider the Brief COPE. I. International Journal of Behavioral Medicine, 4, 92–100.

Caspi, A., Moffit, T. E., & Thorton, A. (1996). The Life History Calender: A research and clinical assessment method for collecting retrospective event-history data. International Journal of Methods in Psychiatric Research, 6(2), 101–114.

Chmitorz, A., Kunzler, A., Helmreich, I., Tüscher, O., Kalisch, R., Kubiak, T., … Lieb, K. (2018). Intervention studies to foster resilience – A systematic review and proposal for a resilience framework in future intervention studies. Clinical Psychology Review, 59, 78–100. https://doi.org/10.1016/j.cpr.2017.11.002

Chmitorz, A., Wenzel, M., Stieglitz, R.-D., Kunzler, A., Bagusat, C., Helmreich, I., … Tüscher, O. (2018). Population-based validation of a German version of the Brief Resilience Scale. PLOS ONE, 13(2), e0192761. https://doi.org/10.1371/journal.pone.0192761

Cieslik, E. C., Mueller, V. I., Eickhoff, C. R., Langner, R., & Eickhoff, S. B. (2015). Three key regions for supervisory attentional control: Evidence from neuroimaging meta-analyses. Neuroscience & Biobehavioral Reviews, 48, 22–34. https://doi.org/10.1016/j.neubiorev.2014.11.003

Compas, B. E., Wagner, B. M., Slavin, L. A., & Vannatta, K. (1986). A prospective study of life events, social support, and psychological symptomatology during the transition from high school to college. American Journal of Community Psychology, 14(3), 241–257.

Craske, M. G., Wolitzky-Taylor, K. B., Mineka, S., Zinbarg, R., Waters, A. M., Vrshek-Schallhorn, S., … Ornitz, E. (2012). Elevated responding to safe conditions as a specific risk factor for anxiety versus depressive disorders: Evidence from a longitudinal investigation. Journal of Abnormal Psychology, 121(2), 315–324. https://doi.org/10.1037/a0025738

Dalgard, O. S., Bjork, S., & Tambs, K. (1995). Social support, negative life events and mental health. The British Journal of Psychiatry, 166, 29–34.

de Vries, G.-J., & Olff, M. (2009). The lifetime prevalence of traumatic events and posttraumatic stress disorder in the Netherlands. Journal of Traumatic Stress, 22(4), 259–267. https://doi.org/10.1002/jts.20429

Desikan, R. S., Ségonne, F., Fischl, B., Quinn, B. T., Dickerson, B. C., Blacker, D., … Killiany, R. J. (2006). An automated labeling system for subdividing the human cerebral cortex on MRI scans into gyral based regions of interest. NeuroImage, 31(3), 968–980. https://doi.org/10.1016/j.neuroimage.2006.01.021

Dickinson, A., & Pearce, J. M. (1977). Inhibitory interactions between appetitive and aversive stimuli. Psychological Bulletin, 84, 690–711.

Diedrichsen, J., Balsters, J. H., Flavell, J., Cussans, E., & Ramnani, N. (2009). A probabilistic MR atlas of the human cerebellum. NeuroImage, 46(1), 39–46. https://doi.org/10.1016/j.neuroimage.2009.01.045

Erk, S., Kleczar, A., & Walter, H. (2007). Valence-specific regulation effects in a working memory task with emotional context. NeuroImage, 37(2), 623–632. https://doi.org/10.1016/j.neuroimage.2007.05.006

Etkin, A., Egner, T., Peraza, D. M., Kandel, E. R., & Hirsch, J. (2006). Resolving Emotional Conflict: A Role for the Rostral Anterior Cingulate Cortex in Modulating Activity in the Amygdala. Neuron, 51(6), 871–882. https://doi.org/10.1016/j.neuron.2006.07.029

Euteneuer, F., Süßenbach, P., Schäfer, S. J., & Rief, W. (2014). Subjektiver sozialer Status. MacArthur-Skalen zur Erfassung des wahrgenommenen sozialen Status im sozialen Umfeld (SSS-U) und in Deutschland (SSS-D). Verhaltenstherapie, 25(3), 229–232. https://doi.org/10.1159/000371558

Feder, A., Charney, D., & Abe, K. (2011). Neurobiology of resilience. In S. M. Southwick (Ed.), Resilience and mental health: challenges across the lifespan (pp. 1–29). Cambridge [England]: Cambridge University Press.

Feinberg, D. A., Moeller, S., Smith, S. M., Auerbach, E., Ramanna, S., Glasser, M. F., … Yacoub, E. (2010). Multiplexed Echo Planar Imaging for Sub-Second Whole Brain FMRI and Fast Diffusion Imaging. PLoS ONE, 5(12), e15710. https://doi.org/10.1371/journal.pone.0015710

Frank, D. W., Dewitt, M., Hudgens-Haney, M., Schaeffer, D. J., Ball, B. H., Schwarz, N. F., … Sabatinelli, D. (2014). Emotion regulation: Quantitative meta-analysis of functional activation and deactivation. Neuroscience & Biobehavioral Reviews, 45, 202–211. https://doi.org/10.1016/j.neubiorev.2014.06.010

Frazier, J. A., Chiu, S., Breeze, J. L., Makris, N., Lange, N., Kennedy, D. N., … Biederman, J. (2005). Structural Brain Magnetic Resonance Imaging of Limbic and Thalamic Volumes in Pediatric Bipolar Disorder. American Journal of Psychiatry, 162(7), 1256–1265. https://doi.org/10.1176/appi.ajp.162.7.1256

Friedman, A. K., Walsh, J. J., Juarez, B., Ku, S. M., Chaudhury, D., Wang, J., … Han, M.-H. (2014). Enhancing Depression Mechanisms in Midbrain Dopamine Neurons Achieves Homeostatic Resilience. Science, 344(6181), 313–319. https://doi.org/10.1126/science.1249240

Fullana, M. A., Albajes-Eizagirre, A., Soriano-Mas, C., Vervliet, B., Cardoner, N., Benet, O., … Harrison, B. J. (2018). Fear extinction in the human brain: A meta-analysis of fMRI studies in healthy participants. Neuroscience & Biobehavioral Reviews, 88, 16–25. https://doi.org/10.1016/j.neubiorev.2018.03.002

Fullana, M. A., Harrison, B. J., Soriano-Mas, C., Vervliet, B., Cardoner, N., Àvila-Parcet, A., & Radua, J. (2016). Neural signatures of human fear conditioning: an updated and extended meta-analysis of fMRI studies. Molecular Psychiatry, 21(4), 500–508. https://doi.org/10.1038/mp.2015.88

García-García, I., Kube, J., Gaebler, M., Horstmann, A., Villringer, A., & Neumann, J. (2016). Neural processing of negative emotional stimuli and the influence of age, sex and task-related characteristics. Neuroscience & Biobehavioral Reviews, 68, 773–793. https://doi.org/10.1016/j.neubiorev.2016.04.020

Garnefski, N., Kraaij, V., & Spinhoven, P. (2001). Negative life events, cognitive emotion regulation and depression. Personality and Individual Differences, 30, 1311–1327.

Glaesmer, H., Hoyer, J., Klotsche, J., & Herzberg, P. Y. (2008). Die deutsche Version des Life-Orientation-Tests (LOT-R) zum dispositionellen Optimismus und Pessimismus. Zeitschrift für Gesundheitspsychologie, 16(1), 26–31. https://doi.org/10.1026/0943-8149.16.1.26

Goldberg, D. P., & Hillier, V. F. (1979). A scaled version of the General Health Questionnaire. Psychological Medicine, 9(1), 139–145.

Goldstein, J. M., Seidman, L. J., Makris, N., Ahern, T., O’Brien, L. M., Caviness, V. S., … Tsuang, M. T. (2007). Hypothalamic Abnormalities in Schizophrenia: Sex Effects and Genetic Vulnerability. Biological Psychiatry, 61(8), 935–945. https://doi.org/10.1016/j.biopsych.2006.06.027

Green, S. R., Kragel, P. A., Fecteau, M. E., & LaBar, K. S. (2014). Development and validation of an unsupervised scoring system (Autonomate) for skin conductance response analysis. International Journal of Psychophysiology, 91(3), 186–193. https://doi.org/10.1016/j.ijpsycho.2013.10.015

Greicius, M. D., Krasnow, B., Reiss, A. L., & Menon, V. (2003). Functional connectivity in the resting brain: A network analysis of the default mode hypothesis. Proceedings of the National Academy of Sciences, 100(1), 253–258. https://doi.org/10.1073/pnas.0135058100

Gross, J. J. (1998). The emerging field of emotion regulation: an integrative review. Review of General Psychology, 2(3), 271–299.

Gross, J. J. (2001). Emotion regulation in adulthood: Timing is everything. Current Directions in Psychological Science, 10, 214–219.

Haaker, J., Gaburro, S., Sah, A., Gartmann, N., Lonsdorf, T. B., Meier, K., … Kalisch, R. (2013). Single dose of L-dopa makes extinction memories context-independent and prevents the return of fear. Proceedings of the National Academy of Sciences, 110(26), E2428–E2436. https://doi.org/10.1073/pnas.1303061110

Herbst, U., Voeth, M., Eidhoff, A. T., Müller, M., & Stief, S. (2016). Studierendenstress in Deutschland - eine empirische Untersuchung.

Iordan, A. D., & Dolcos, F. (2015). Brain Activity and Network Interactions Linked to Valence-Related Differences in the Impact of Emotional Distraction. Cerebral Cortex, bhv242. https://doi.org/10.1093/cercor/bhv242

Iordan, A. D., Dolcos, S., & Dolcos, F. (2013). Neural signatures of the response to emotional distraction: a review of evidence from brain imaging investigations. Frontiers in Human Neuroscience, 7. https://doi.org/10.3389/fnhum.2013.00200

Jia, T., Macare, C., Desrivières, S., Gonzalez, D. A., Tao, C., Ji, X., … the IMAGEN Consortium. (2016). Neural basis of reward anticipation and its genetic determinants. Proceedings of the National Academy of Sciences, 113(14), 3879–3884. https://doi.org/10.1073/pnas.1503252113

Joseph, S., & Linley, P. A. (2006). Growth following adversity: Theoretical perspectives and implications for clinical practice. Clinical Psychology Review, 26(8), 1041–1053. https://doi.org/10.1016/j.cpr.2005.12.006

Kalanthroff, E., Cohen, N., & Henik, A. (2013). Stop feeling: inhibition of emotional interference following stop-signal trials. Frontiers in Human Neuroscience, 7. https://doi.org/10.3389/fnhum.2013.00078

Kalisch, R. (2006). Context-Dependent Human Extinction Memory Is Mediated by a Ventromedial Prefrontal and Hippocampal Network. Journal of Neuroscience, 26(37), 9503–9511. https://doi.org/10.1523/JNEUROSCI.2021-06.2006

Kalisch, R. (2009). The functional neuroanatomy of reappraisal: Time matters. Neuroscience & Biobehavioral Reviews, 33(8), 1215–1226. https://doi.org/10.1016/j.neubiorev.2009.06.003

Kalisch, R., Baker, D. G., Basten, U., Boks, M. P., Bonanno, G. A., Brummelman, E., … Kleim, B. (2017). The resilience framework as a strategy to combat stress-related disorders. Nature Human Behaviour, 1(11), 784–790. https://doi.org/10.1038/s41562-017-0200-8

Kalisch, R., & Gerlicher, A. M. V. (2014). Making a mountain out of a molehill: On the role of the rostral dorsal anterior cingulate and dorsomedial prefrontal cortex in conscious threat appraisal, catastrophizing, and worrying. Neuroscience & Biobehavioral Reviews, 42, 1–8. https://doi.org/10.1016/j.neubiorev.2014.02.002

Kalisch, R., Müller, M. B., & Tüscher, O. (2015). A conceptual framework for the neurobiological study of resilience. Behavioral and Brain Sciences, 38(E92), 1–49. https://doi.org/10.1017/S0140525X1400082X

Kalisch, R., Wiech, K., Critchley, H. D., & Dolan, R. J. (2006). Levels of appraisal: A medial prefrontal role in high-level appraisal of emotional material. NeuroImage, 30(4), 1458–1466. https://doi.org/10.1016/j.neuroimage.2005.11.011

Kalisch, R., Wiech, K., Critchley, H. D., Seymour, B., O’Doherty, J. P., Oakley, D. A., … Dolan, R. J. (2005). Anxiety reduction through detachment: subjective, physiological, and neural effects. Journal of Cognitive Neuroscience, 17(6), 874–883. https://doi.org/10.1162/0898929054021184

Kanske, P., Heissler, J., Schonfelder, S., Bongers, A., & Wessa, M. (2011). How to Regulate Emotion? Neural Networks for Reappraisal and Distraction. Cerebral Cortex, 21(6), 1379–1388. https://doi.org/10.1093/cercor/bhq216

Kellermann, T. S., Sternkopf, M. A., Schneider, F., Habel, U., Turetsky, B. I., Zilles, K., & Eickhoff, S. B. (2012). Modulating the processing of emotional stimuli by cognitive demand. Social Cognitive and Affective Neuroscience, 7(3), 263–273. https://doi.org/10.1093/scan/nsq104

Kemper, C. J., Ziegler, M., & Taylor, S. (2009). Überprüfung der psychometrischen Qualität der deutschen Version des Angstsensitivitätsindex-3. Diagnostica, 55(4), 223–233. https://doi.org/10.1026/0012-1924.55.4.223

Kent, M., Davis, M. C., & Reich, J. W. (Eds.). (2014). The resilience handbook: approaches to stress and trauma. New York, NY: Routledge, Taylor & Francis Group.

Kessler, R. C., Angermeyer, M., Anthony, J. C., DE Graaf, R., Demyttenaere, K., Gasquet, I., … Ustün, T. B. (2007). Lifetime prevalence and age-of-onset distributions of mental disorders in the World Health Organization’s World Mental Health Survey Initiative. World Psychiatry: Official Journal of the World Psychiatric Association (WPA), 6(3), 168–176.

Kessler, R. C., Berglund, P., Demler, O., Jin, R., Merikangas, K. R., & Walters, E. E. (2005). Lifetime Prevalence and Age-of-Onset Distributions of DSM-IV Disorders in the National Comorbidity Survey Replication. Archives of General Psychiatry, 62(6), 593. https://doi.org/10.1001/archpsyc.62.6.593

Kilpatrick, D. G., Resnick, H. S., Milanak, M. E., Miller, M. W., Keyes, K. M., & Friedman, M. J. (2013). National Estimates of Exposure to Traumatic Events and PTSD Prevalence Using DSM-IV and DSM-5 Criteria: DSM-5 PTSD Prevalence. Journal of Traumatic Stress, 26(5), 537–547. https://doi.org/10.1002/jts.21848

Kirschbaum, C., & Hellhammer, D. H. (1989). Salivary Cortisol in Psychobiological Research: An Overview. Neuropsychobiology, 22(3), 150–169. https://doi.org/10.1159/000118611

Klaiberg, A., Schumacher, J., & Brähler, E. (2004). General Health Questionnaire 28 (GHQ-28): Teststatistische Überprüfung einer deutschen Version in einer bevölkerungsrepräsentativen Stichprobe. Zeitschrift Für Klinische Psychologie, Psychiatrie Und Psychotherapie, 52, 31–42.

Knoll, N., Rieckmann, N., & Schwarzerq, R. (2005). Coping as a mediator between personality and stress outcomes: A longitudinal study with cataract surgery patients. European Journal of Personality, 19(3), 229–247.

Knutson, B., Adams, C. M., Fong, G. W., & Hommer, D. (2001). Anticipation of Increasing Monetary Reward Selectively Recruits Nucleus Accumbens. The Journal of Neuroscience, 21.

Kohn, N., Eickhoff, S. B., Scheller, M., Laird, A. R., Fox, P. T., & Habel, U. (2014). Neural network of cognitive emotion regulation — An ALE meta-analysis and MACM analysis. NeuroImage, 87, 345–355. https://doi.org/10.1016/j.neuroimage.2013.11.001

Konorski. (1967). Integrative activity of the brain: an interdisciplinary approach. Chicago: The University of Chicago Press.

Krishnan, V., Han, M.-H., Graham, D. L., Berton, O., Renthal, W., Russo, S. J., … Nestler, E. J. (2007). Molecular Adaptations Underlying Susceptibility and Resistance to Social Defeat in Brain Reward Regions. Cell, 131(2), 391–404. https://doi.org/10.1016/j.cell.2007.09.018

Lang, P. J., Bradley, M. M., & Cuthbert, B. N. (2008). International affective picture system (IAPS): Affective ratings of pictures and instruction manual. Technical Report A-8. University of Florida, Gainesville, FL.

Lazarus, R. S., & Folkman, S. (1984). Stress, appraisal and coping. New York: Springer.

Lejuz, C., Kahler, C. W., & Brown, R. A. (2003). A modified computer version of the Paced Auditory Serial Addition Task (PASAT) as a laboratory-based stressor. Behavior Therapist, 290–293.

Leventhal, H., & Scherer, K. (1987). The Relationship of Emotion to Cognition: A Functional Approach to a Semantic Controversy. Cognition & Emotion, 1(1), 3–28. https://doi.org/10.1080/02699938708408361

Lindquist, K. A., Satpute, A. B., Wager, T. D., Weber, J., & Barrett, L. F. (2016). The Brain Basis of Positive and Negative Affect: Evidence from a Meta-Analysis of the Human Neuroimaging Literature. Cerebral Cortex, 26(5), 1910–1922. https://doi.org/10.1093/cercor/bhv001

Liu, X., Hairston, J., Schrier, M., & Fan, J. (2011). Common and distinct networks underlying reward valence and processing stages: A meta-analysis of functional neuroimaging studies. Neuroscience & Biobehavioral Reviews, 35(5), 1219–1236. https://doi.org/10.1016/j.neubiorev.2010.12.012

Loch, N., Hiller, W., & Witthöft, M. (2011). Der Cognitive Emotion Regulation Questionnaire (CERQ): Erste teststatistische Überprüfung einer deutschen Adaption. Zeitschrift Für Klinische Psychologie Und Psychotherapie, 40(2), 94–106. https://doi.org/10.1026/1616-3443/a000079

Lonsdorf, T. B., Haaker, J., & Kalisch, R. (2014). Long-term expression of human contextual fear and extinction memories involves amygdala, hippocampus and ventromedial prefrontal cortex: a reinstatement study in two independent samples. Social Cognitive and Affective Neuroscience, 9(12), 1973–1983. https://doi.org/10.1093/scan/nsu018

Lonsdorf, T. B., Menz, M. M., Andreatta, M., Fullana, M. A., Golkar, A., Haaker, J., … Merz, C. J. (2017). Don’t fear “fear conditioning”: Methodological considerations for the design and analysis of studies on human fear acquisition, extinction, and return of fear. Neuroscience & Biobehavioral Reviews, 77, 247–285. https://doi.org/10.1016/j.neubiorev.2017.02.026

Luthar, S. S., Cicchetti, D., & Becker, B. (2000). The construct of resilience: a critical evaluation and guidelines for future work. Child Development, 71(3), 543–562.

Maier, S. F. (2015). Behavioral control blunts reactions to contemporaneous and future adverse events: Medial prefrontal cortex plasticity and a corticostriatal network. Neurobiology of Stress, 1, 12–22. https://doi.org/10.1016/j.ynstr.2014.09.003

Makris, N., Goldstein, J. M., Kennedy, D., Hodge, S. M., Caviness, V. S., Faraone, S. V., … Seidman, L. J. (2006). Decreased volume of left and total anterior insular lobule in schizophrenia. Schizophrenia Research, 83(2–3), 155–171. https://doi.org/10.1016/j.schres.2005.11.020

Mancini, A. D., & Bonanno, G. A. (2009). Predictors and Parameters of Resilience to Loss: Toward an Individual Differences Model. Journal of Personality, 77(6), 1805–1832. https://doi.org/10.1111/j.1467-6494.2009.00601.x

Mancini, A. D., Bonanno, G. A., & Sinan, B. (2015). A Brief Retrospective Method for Identifying Longitudinal Trajectories of Adjustment Following Acute Stress. Assessment, 22(3), 298–308. https://doi.org/10.1177/1073191114550816

Marchewka, A., Żurawski, Ł., Jednoróg, K., & Grabowska, A. (2014). The Nencki Affective Picture System (NAPS): Introduction to a novel, standardized, wide-range, high-quality, realistic picture database. Behavior Research Methods, 46(2), 596–610. https://doi.org/10.3758/s13428-013-0379-1

McEwen, B. S. (1993). Stress and the Individual: Mechanisms Leading to Disease. Archives of Internal Medicine, 153(18), 2093. https://doi.org/10.1001/archinte.1993.00410180039004

Mechias, M.-L., Etkin, A., & Kalisch, R. (2010). A meta-analysis of instructed fear studies: Implications for conscious appraisal of threat. NeuroImage, 49(2), 1760–1768. https://doi.org/10.1016/j.neuroimage.2009.09.040

Milad, M. R., Wright, C. I., Orr, S. P., Pitman, R. K., Quirk, G. J., & Rauch, S. L. (2007). Recall of Fear Extinction in Humans Activates the Ventromedial Prefrontal Cortex and Hippocampus in Concert. Biological Psychiatry, 62(5), 446–454. https://doi.org/10.1016/j.biopsych.2006.10.011

Nasser, H. M., & McNally, G. P. (2012). Appetitive–aversive interactions in Pavlovian fear conditioning. Behavioral Neuroscience, 126(3), 404–422. https://doi.org/10.1037/a0028341

Pan, J., & Tompkins, W. J. (1983). A Real-Time QRS Detection Algorithm. IEEE TRANSACTIONS ON BIOMEDICAL ENGINEERING, VOL. BME-32(NO. 3).

Paret, C., Brenninkmeyer, J., Meyer, B., Yuen, K. S. L., Gartmann, N., Mechias, M.-L., & Kalisch, R. (2011). A Test for the Implementation? Maintenance Model of Reappraisal. Frontiers in Psychology, 2. https://doi.org/10.3389/fpsyg.2011.00216

Park, J., Ayduk, Ö., O’Donnell, L., Chun, J., Gruber, J., Kamali, M., … Kross, E. (2014). Regulating the High: Cognitive and Neural Processes Underlying Positive Emotion Regulation in Bipolar I Disorder. Clinical Psychological Science, 2(6), 661–674. https://doi.org/10.1177/2167702614527580

Plichta, M. M., Grimm, O., Morgen, K., Mier, D., Sauer, C., Haddad, L., … Meyer-Lindenberg, A. (2014). Amygdala habituation: A reliable fMRI phenotype. NeuroImage, 103, 383–390. https://doi.org/10.1016/j.neuroimage.2014.09.059

Ragland, G. B., & Shulkin, J. (2014). Introduction to allostasis and allostatic load. In M. Kent, M. C. Davis, & J. W. Reich (Eds.), The resilience handbook (pp. 44–52). New York, NY: Routledge, Taylor & Francis Group.

Reavley, N., & Jorm, A. F. (2010). Prevention and early intervention to improve mental health in higher education students: a review: Mental health in higher education students. Early Intervention in Psychiatry, 4(2), 132–142. https://doi.org/10.1111/j.1751-7893.2010.00167.x

Reinhardt, T., Schmahl, C., Wüst, S., & Bohus, M. (2012). Salivary cortisol, heart rate, electrodermal activity and subjective stress responses to the Mannheim Multicomponent Stress Test (MMST). Psychiatry Research, 198(1), 106–111. https://doi.org/10.1016/j.psychres.2011.12.009

Roalf, D. R., Ruparel, K., Gur, R. E., Bilker, W., Gerraty, R., Elliott, M. A., … Gur, R. C. (2014). Neuroimaging predictors of cognitive performance across a standardized neurocognitive battery. Neuropsychology, 28(2), 161–176. https://doi.org/10.1037/neu0000011

Robinson, M. D. (1998). Running from William James’ Bear: A Review of Preattentive Mechanisms and their Contributions to Emotional Experience. Cognition & Emotion, 12(5), 667–696. https://doi.org/10.1080/026999398379493

Rohleder, N., Wolf, J. M., Maldonado, E. F., & Kirschbaum, C. (2006). The psychosocial stress-induced increase in salivary alpha-amylase is independent of saliva flow rate. Psychophysiology, 43(6), 645–652. https://doi.org/10.1111/j.1469-8986.2006.00457.x

Rohr, C. S., Dreyer, F. R., Aderka, I. M., Margulies, D. S., Frisch, S., Villringer, A., & Okon-Singer, H. (2015). Individual differences in common factors of emotional traits and executive functions predict functional connectivity of the amygdala. NeuroImage, 120, 154–163. https://doi.org/10.1016/j.neuroimage.2015.06.049

Rotenstein, L. S., Ramos, M. A., Torre, M., Segal, J. B., Peluso, M. J., Guille, C., … Mata, D. A. (2016). Prevalence of Depression, Depressive Symptoms, and Suicidal Ideation Among Medical Students: A Systematic Review and Meta-Analysis. JAMA, 316(21), 2214. https://doi.org/10.1001/jama.2016.17324

Russo, S. J., Murrough, J. W., Han, M.-H., Charney, D. S., & Nestler, E. J. (2012). Neurobiology of resilience. Nature Neuroscience, 15(11), 1475–1484. https://doi.org/10.1038/nn.3234

Rutter, M. (2012). Resilience as a dynamic concept. Development and Psychopathology, 24(02), 335–344. https://doi.org/10.1017/S0954579412000028

Sander, D., Grandjean, D., & Scherer, K. R. (2005). A systems approach to appraisal mechanisms in emotion. Neural Networks, 18(4), 317–352. https://doi.org/10.1016/j.neunet.2005.03.001

Sapienza, J. K., & Masten, A. S. (2011). Understanding and promoting resilience in children and youth: Current Opinion in Psychiatry, 24(4), 267–273. https://doi.org/10.1097/YCO.0b013e32834776a8

Scheier, M. F., & Carver, C. S. (1985). Optimism, Coping, and Health: Assessment and Implications of Generalized Outcome Expectancies. Health Psychology, 4.

Scherer, K. R. (2001). Appraisal considered as a process of multilevel sequential checking. In K. R. Scherer, A. Schorr, & T. Johnstone (Eds.), Appraisal processes in emotion: theory, methods, research (pp. 92–120). New York: Oxford University Press.

Schwarzer, R., & Jerusalem, M. (Eds.). (1999). Skalen zur Erfassung von Lehrer- und Schülermerkmalen. Dokumentation der psychometrischen Verfahren im Rahmen der Wissenschaftlichen Begleitung des Modellversuchs Selbstwirksame Schulen. Berlin: Freie Universität.

Sebastian, A., Jung, P., Neuhoff, J., Wibral, M., Fox, P. T., Lieb, K., … Mobascher, A. (2016). Dissociable attentional and inhibitory networks of dorsal and ventral areas of the right inferior frontal cortex: a combined task-specific and coordinate-based meta-analytic fMRI study. Brain Structure and Function, 221(3), 1635–1651. https://doi.org/10.1007/s00429-015-0994-y

Sebastian, A., Pohl, M. F., Klöppel, S., Feige, B., Lange, T., Stahl, C., … Tüscher, O. (2013). Disentangling common and specific neural subprocesses of response inhibition. NeuroImage, 64, 601–615. https://doi.org/10.1016/j.neuroimage.2012.09.020

Seery, M. D., Holman, E. A., & Silver, R. C. (2010). Whatever does not kill us: Cumulative lifetime adversity, vulnerability, and resilience. Journal of Personality and Social Psychology, 99(6), 1025–1041. https://doi.org/10.1037/a0021344

Seery, M. D., Leo, R. J., Lupien, S. P., Kondrak, C. L., & Almonte, J. L. (2013). An Upside to Adversity?: Moderate Cumulative Lifetime Adversity Is Associated With Resilient Responses in the Face of Controlled Stressors. Psychological Science, 24(7), 1181–1189. https://doi.org/10.1177/0956797612469210

Smith, B. W., Dalen, J., Wiggins, K., Tooley, E., Christopher, P., & Bernard, J. (2008). The brief resilience scale: Assessing the ability to bounce back. International Journal of Behavioral Medicine, 15(3), 194–200. https://doi.org/10.1080/10705500802222972

Southwick, S. M., & Charney, D. S. (2012). The Science of Resilience: Implications for the Prevention and Treatment of Depression. Science, 338(6103), 79–82. https://doi.org/10.1126/science.1222942

Sterling, P., & Eyer, J. (1988). Allostasis: a new paradigm to explain arousal pathways. In Handbook of life stress, cognition and health (pp. 629–649). New York: Wiley.

Stewart, D. E., & Yuen, T. (2011). A Systematic Review of Resilience in the Physically Ill. Psychosomatics, 52(3), 199–209. https://doi.org/10.1016/j.psym.2011.01.036

Taylor, S., Zvolensky, M. J., Cox, B. J., Deacon, B., Heimberg, R. G., Ledley, D. R., … Cardenas, S. J. (2007). Robust dimensions of anxiety sensitivity: Development and initial validation of the Anxiety Sensitivity Index-3. Psychological Assessment, 19(2), 176–188. https://doi.org/10.1037/1040-3590.19.2.176

Tedeschi, R. G. (2011). Posttraumatic Growth in Combat Veterans. Journal of Clinical Psychology in Medical Settings, 18(2), 137–144. https://doi.org/10.1007/s10880-011-9255-2

Vanderwal, T., Kelly, C., Eilbott, J., Mayes, L. C., & Castellanos, F. X. (2015). Inscapes: A movie paradigm to improve compliance in functional magnetic resonance imaging. NeuroImage, 122, 222–232. https://doi.org/10.1016/j.neuroimage.2015.07.069

Vansteenwegen, D., Crombez, G., Baeyens, F., & Eelen, P. (1998). Extinction in fear conditioning: Effects on startle modulation and evaluative self-reports. Psychophysiology, 35(6), 729–736. https://doi.org/10.1111/1469-8986.3560729

Verbruggen, F., & De Houwer, J. (2007). Do emotional stimuli interfere with response inhibition? Evidence from the stop signal paradigm. Cognition & Emotion, 21(2), 391–403. https://doi.org/10.1080/02699930600625081

Verduyn, P., Van Mechelen, I., Kross, E., Chezzi, C., & Van Bever, F. (2012). The relationship between self-distancing and the duration of negative and positive emotional experiences in daily life. Emotion, 12(6), 1248–1263. https://doi.org/10.1037/a0028289

Wang, M., Perova, Z., Arenkiel, B. R., & Li, B. (2014). Synaptic Modifications in the Medial Prefrontal Cortex in Susceptibility and Resilience to Stress. Journal of Neuroscience, 34(22), 7485–7492. https://doi.org/10.1523/JNEUROSCI.5294-13.2014

Weiner, H. (1992). Perturbing the Organism: The Biology of Stressful Experience. Chicago: University of Chicago Press.

Werner, E. (1992). The children of Kauai: Resiliency and recovery in adolescence and adulthood1. Journal of Adolescent Health, 13(4), 262–268. https://doi.org/10.1016/1054-139X(92)90157-7

Wessa, M., Heissler, J., Schönfelder, S., & Kanske, P. (2013). Goal-directed behavior under emotional distraction is preserved by enhanced task-specific activation. Social Cognitive and Affective Neuroscience, 8(3), 305–312. https://doi.org/10.1093/scan/nsr098

Wessa, M., Kanske, P., Neumeister, P., Bode, K., Heissler, J., & Schönfelder, S. (2010). EmoPics: Subjektive und psychophysiologische Evaluationen neuen Bildmaterials für die klinisch-bio-psychologische Forschung. Zeitschrift Für Klinischer Psychologie Und Psychotherapie, Supplement, 1/11, 77.

Wittchen, H. U., Jacobi, F., Rehm, J., Gustavsson, A., Svensson, M., Jönsson, B., … Steinhausen, H.-C. (2011). The size and burden of mental disorders and other disorders of the brain in Europe 2010. European Neuropsychopharmacology, 21(9), 655–679. https://doi.org/10.1016/j.euroneuro.2011.07.018

Wu, C. C., Samanez-Larkin, G. R., Katovich, K., & Knutson, B. (2014). Affective traits link to reliable neural markers of incentive anticipation. NeuroImage, 84, 279–289. https://doi.org/10.1016/j.neuroimage.2013.08.055

